# A nano-liquid hub integrating growth and immune-evasion signaling to promote cell survival

**DOI:** 10.1101/2021.12.30.474523

**Authors:** Taka A. Tsunoyama, Christian Hoffmann, Daiki Sasaki, Cheng Tan, Bo Tang, Koichiro M. Hirosawa, Yuri L. Nemoto, Rinshi S. Kasai, Takahiro K. Fujiwara, Kenichi G. N. Suzuki, Yuji Sugita, Hiroki Ishikawa, Dragomir Milovanovic, Akihiro Kusumi

## Abstract

Cell survival in immune-competent organisms relies on both growth and immune evasion, yet these processes have been studied as independent cellular programs. Here, using super-resolution single-molecule imaging, we identify GEM (Growth and Evasion Metastable hub), a ≈34-nm, metastable, nano-liquid, plasma membrane hub that physically integrates receptor-type tyrosine kinase (RTK) inputs with CD59-mediated immune-evasion signaling within the same assembly. GEM is formed by LLPS-like protein clustering on PI(4,5)P_2_-containing raft-nanodomains. Upon concurrent RTK and CD59 stimulations, GEM recruits both receptors and downstream kinases, enabling reciprocal activation via nano-liquid confinement that supralinearly amplifies PLCγ–IP_3_–Ca^2+^ and PI3K–Akt survival signaling outputs. In mice, disrupting GEM suppresses tumor growth *in vivo*, establishing nano-liquid GEM as an integrator that physically couples growth and CD59-mediated immune-evasion signaling for cell survival.

## Main Text

Cell survival in multicellular organisms *with active immune systems* depends on both growth signaling and mechanisms that protect cells from self-directed immune attack, yet these processes have traditionally been studied as independent cellular programs. As a result, whether growth and immune-evasion signals can be coordinated, let alone organized within shared plasma-membrane (PM) architectures, has remained largely unexplored. To address this issue, we asked whether and how growth and immune-evasion signals might be physically coordinated by dedicated architectures.

Growth signaling, typically mediated by receptor-type tyrosine kinases (RTKs) at the plasma membrane (PM), has been extensively characterized as a central driver of cell proliferation and survival (*1*). Immune evasion can occur through diverse mechanisms; in this study, we focus on immune evasion in the form of resistance to complement-dependent cytotoxicity (CDC), because CDC, like RTK signaling, is initiated at the PM. CD59, a glycosylphosphatidylinositol (GPI)-anchored, raftophilic receptor expressed in virtually all cell types, is a central mediator of resistance to CDC. In the context of cancer biology, many tumors overexpress CD59 (*2*). CD59 prevents CDC by binding the complement C5b-8 complex, the membrane attack complex (MAC) precursor, to inhibit MAC formation (*3*). Beyond this canonical role, the engagement of CD59 by MAC precursors has been reported to elicit rapid oligomerization and recruitment of cytoplasmic signaling molecules, including trimeric G proteins, Src-family kinases (SFKs) and phospholipase Cγ (PLCγ), thereby triggering proliferative signaling outputs (*4–7*). PLCγ recruitment and activation occur at CD59-cluster-associated platforms (*4–6*). Because these signaling molecules also operate downstream of growth-factor RTKs, growth and immune evasion signals might converge at shared membrane-associated signaling sites. Because both RTKs (*8–10*) and GPI-anchored receptors (*11–15*) can interact with integrins, we asked whether integrin-associated assemblies could provide a physical basis for such convergence.

Here, using advanced super-resolution single-molecule imaging and tracking, we identify GEM (Growth and Evasion Metastable hub), a nanoscale (≈34 nm), liquid-like, metastable, integrin-associated assemblies that integrates CD59-mediated immune-evasion signaling and growth-factor signaling. GEM consists of integrin, talin, zyxin, and other proteins associated with integrin adhesion complexes (IACs), while remaining distinct from canonical IACs, because GEM is present in the apical PM and it lacks vinculin and paxillin, essential constituent molecules for canonical IACs. GEM assembly is driven primarily by zyxin’s intrinsically disordered region (IDR) and is supported by PI(4,5)P_2_-containing raft nanodomains and cortical actin, exhibiting hallmark behaviors of liquid-liquid phase separation (LLPS)-like clustering. The nano-liquid organization provides a physical basis for bringing together structurally diverse receptors and signaling molecules via weak, multivalent interactions. GEMs form constitutively at endogenous protein expression levels, at densities comparable to clathrin-coated pits (several hundred GEMs/cell), persists for ≈8.5 s, and exchange constituents with their cytoplasmic pools on second timescales.

Upon concurrent RTK and CD59 stimulations, GEM recruits both receptors together with their respective downstream kinases, SFKs and focal adhesion kinase (FAK), and cooperatively produces supralinear amplification of PLCγ-IP_3_-Ca^2+^ and PI3K-Akt signals, key signaling outputs essential for cell survival. We propose that confinement within the nano-liquid GEM environment enables cooperative kinase activation that underlies this supralinearity. GEM also enhances CD59-dependent immune evasion from CDC by facilitating encounters between CD59 and MAC precursors. In murine tumor models, restoration of GEM formation markedly enhances tumor growth, while disruption of GEM attenuates it.

Taken together, these findings identify GEM as a metastable, nano-liquid PM hub in which growth signaling and CD59-associated complement evasion, long treated as independent survival programs, are physically coordinated within a shared nano-liquid domain. This framework links nanoscale architecture on the PM to functional coupling of proliferative signaling and CDC resistance *in vivo*, and provides a basis for understanding how such organization can be leveraged in cancer.

## Results

### GPI-anchored receptors, engaged RTKs, and PLCγ1 are transiently recruited to talin clusters on the apical plasma membrane

Talin is a large, multidomain integrin-associated protein that links integrins to multiple interaction partners and is essential for the assembly and function of integrin–adhesion complexes (IACs; throughout this report, IACs include mature focal adhesions) (*16–18*). We therefore began by asking whether talin-centered assemblies at the plasma membrane (PM) engage both growth signaling and CD59-mediated immune-evasion (CDC-resistance) pathways.

Unexpectedly, we found that talin forms discrete clusters on the apical PM, where canonical IACs are absent. These talin clusters are immobile or slowly-diffusive and metastable, repeatedly assembling and disassembling (Fig. 1A and movie S1) with an exponential lifetime of 9.0 s (Fig. 1B; τ_2_; see materials and methods). These clusters were detected using Halo7-tagged-talin1 molecules expressed in talin1/2 double-knockout mouse embryonic fibroblasts (dKO-MEFs) (*19*) at levels comparable to endogenous talin1 (Fig. 1A, fig. S1, A to G, and data S1; see materials and methods). Unless otherwise stated, tagged talin1 is simply referred to as talin hereafter. Raw data and statistical parameters, including SEMs, *p*-values, and numerical original data, are summarized in data S2.

**Fig. 1.**
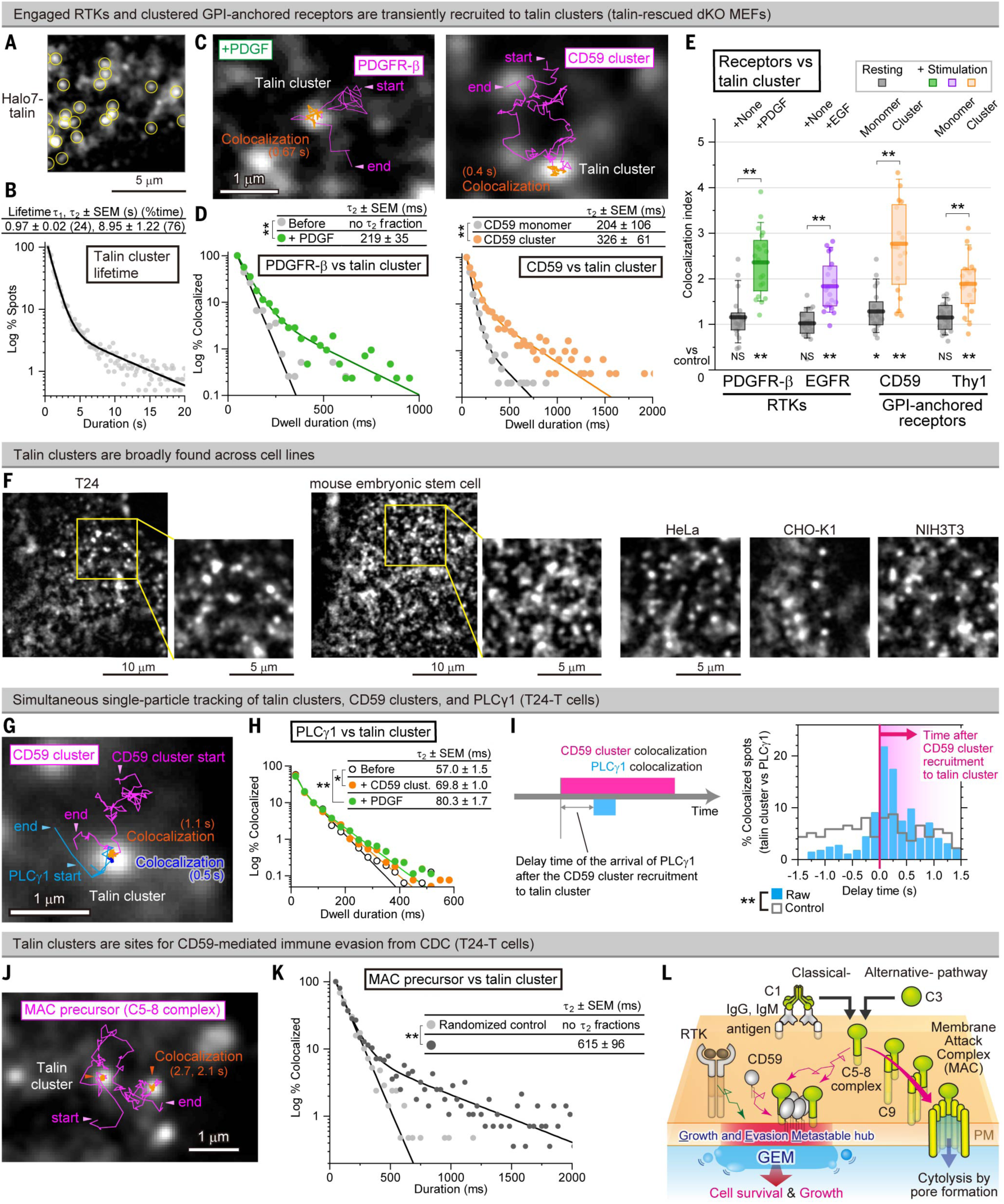
Single-molecule imaging reveals a nanoscale liquid-like signaling hub (growth and evasion metastable hub=GEM). (**A**) Single-molecule visualization of fluorescently labeled talin clusters located on the apical PM. (**B**) Distributions of talin cluster lifetimes. (**C and D**) Intermittent, transient recruitment of engaged PDGFR and CD59 (trajectories) to talin clusters (white spots) (C, snapshot from movie S2) and their dwell lifetimes at talin clusters (τ_2_; D). (**E**) Single-molecule colocalization indexes of various receptors with talin clusters. (**F**) Typical images of talin clusters (overexpressed Halo7-talin) in various cell lines. (**G to I**) Immediately following the recruitment of engaged CD59 clusters to talin clusters, PLCγ1(-Halo7) is recruited to the same talin clusters. (G) Representative talin cluster image and trajectories of a PLCγ1 molecule and a CD59 cluster (snapshot from movie S3). (H) Distribution of the dwell durations of PLCγ1 on talin clusters. (I) Timing of PLCγ1 recruitment relative to CD59-cluster recruitment (>3-frame colocalizations). (**J and K**) Transient recruitment of MAC precursors (trajectory; J) to talin clusters (white puncta) and their dwell lifetimes at talin clusters (K). (**L**) Schematic model illustrating that CD59, MAC precursors, and PDGFR are recruited to GEM (growth and evasion metastable hub). Throughout this study (unless otherwise noted), bars, boxes, and whiskers indicate mean values, interquartile ranges (25–75%), and 10–90% ranges, respectively, and NS, not significant; *, *P* < 0.05; and **, *P* < 0.01. Brunner-Munzel test for (D), two-tailed Welch’s t-test for (E), Steel-Dwass’ test for (H and K).

Single-molecule imaging revealed that ligated PDGFR molecules and signaling-active CD59 clusters (each consisting of an average of five CD59 molecules and recapitulating the signaling-competent oligomers described previously (*4*, *5*)) were transiently recruited to talin clusters on the apical PM (Fig. 1, C and D). Their dwell lifetimes were on the sub-second timescale (exponential time constant, τ_2_ (*20*); Fig. 1D and movie S2; see materials and methods). In contrast, non-ligated PDGFR molecules were not recruited to talin clusters, whereas non-clustered CD59 molecules were recruited less frequently and with shorter dwell lifetimes (Fig. 1D).

To quantify recruitment, we used a single-molecule colocalization index (*20*, *21*) (fig. S1H). Engaged PDGFR and CD59 showed robust recruitment, and similar recruitment was observed for another RTK, EGFR, and another GPI-anchored receptor, Thy1 (Fig. 1E), indicating that apical talin clusters can recruit receptors with highly diverse molecular architectures.

Talin clusters were detected in all five mammalian cell lines examined (Fig. 1F). Because apical talin clusters were most readily visualized in human epithelial T24 cells, owing to their large, flat apical PM regions, we generated a T24-derived line stably expressing SNAPf-talin at 0.7x the endogenous talin level (T24-T cells; fig. S2), and used this line extensively in subsequent experiments. As in MEFs, individual engaged CD59 clusters and ligand-bound PDGFR molecules were transiently recruited to talin clusters in T24-T cells (fig. S2, D and E).

CD59 clusters are known to recruit a downstream cytoplasmic signaling protein involved in cell growth, PLCγ, at CD59 cluster-associated platforms (*4–6*). We therefore examined whether talin clusters might serve as the physical sites where this signaling-related function of CD59 is executed. Using triple-color single-molecule imaging, we found that following the recruitment of an engaged CD59 cluster to a talin cluster, PLCγ1 molecules are transiently recruited to the same talin cluster (≈70 ms), predominantly immediately after CD59 arrival (Fig. 1, G to I, and movie S3). Because CD59 cannot directly bind PLCγ1, these observations suggest that, following CD59 engagement, tyrosine phosphorylated docking sites are generated at the talin cluster (*4–7*). Upon PDGF stimulation, PLCγ1 is likewise transiently recruited to talin clusters with a comparable dwell lifetime (≈80 ms; Fig. 1H).

### Talin clusters contribute to CD59-mediated immune evasion

Whereas the preceding subsection addressed CD59-dependent growth signaling at talin clusters, we next examined whether talin clusters also contribute to CD59’s canonical immune evasion function, resistance to CDC (*2*). Specifically, we tested whether the bona-fide ligands of CD59, MAC precursors (C5b-8 complexes), are recruited to talin clusters. Using single-molecule imaging, we found that MAC precursors were transiently recruited to talin clusters (Fig 1, J and K).

Because both ligated and non-ligated CD59 are recruited to talin clusters, these observations indicate that talin clusters can co-enrich CD59 and MAC precursors, enhancing local CD59-MAC precursor encounters and thereby suppressing C9 recruitment and MAC assembly (*3*). Therefore, talin clusters are well positioned to facilitate CD59-mediated immune evasion via resistance to CDC (Fig. 1L; also see Fig. 4M).

Taken together with the results described in the preceding subsection, talin clusters show three defining features: (1) they engage growth signaling downstream from both RTKs and CD59 (Fig. 1, C to E and G to I); (2) they are associated with CD59-mediated immune evasion via resistance to CDC (Fig. 1, J and K); and (3) they are metastable structures that repeatedly assemble and disassemble with short lifetimes (Fig. 1B). Based on these combined growth- and evasion-related functions, we refer to these talin clusters as GEMs (Growth and Evasion Metastable hubs) (Fig. 1L). GEM constituent molecules (including integrin), formation mechanisms, and functional consequences are described in subsequent subsections.

### GEM forms metastable liquid-like co-assemblies with zyxin

To further characterize GEM molecular composition and dynamics, we next asked whether another IAC protein, zyxin, is incorporated into GEMs. We employed zyxin-KO MEFs (*22*) rescued with Halo7-tagged zyxin, expressed at near-endogenous levels (fig. S1, D to F). Halo7-zyxin also formed clusters on the apical PM (fig. S1G and movie S1), and ≈70% of talin clusters colocalized with zyxin clusters (Fig. 2, A and B), indicating that zyxin is a major constituent of GEMs. Accordingly, we identify GEMs as clusters detected by talin or zyxin imaging, and report the mean of metrics obtained from talin-defined and zyxin-defined GEMs in the main text (and thus these values differ slightly in the raw data shown in individual figures).

**Fig. 2.**
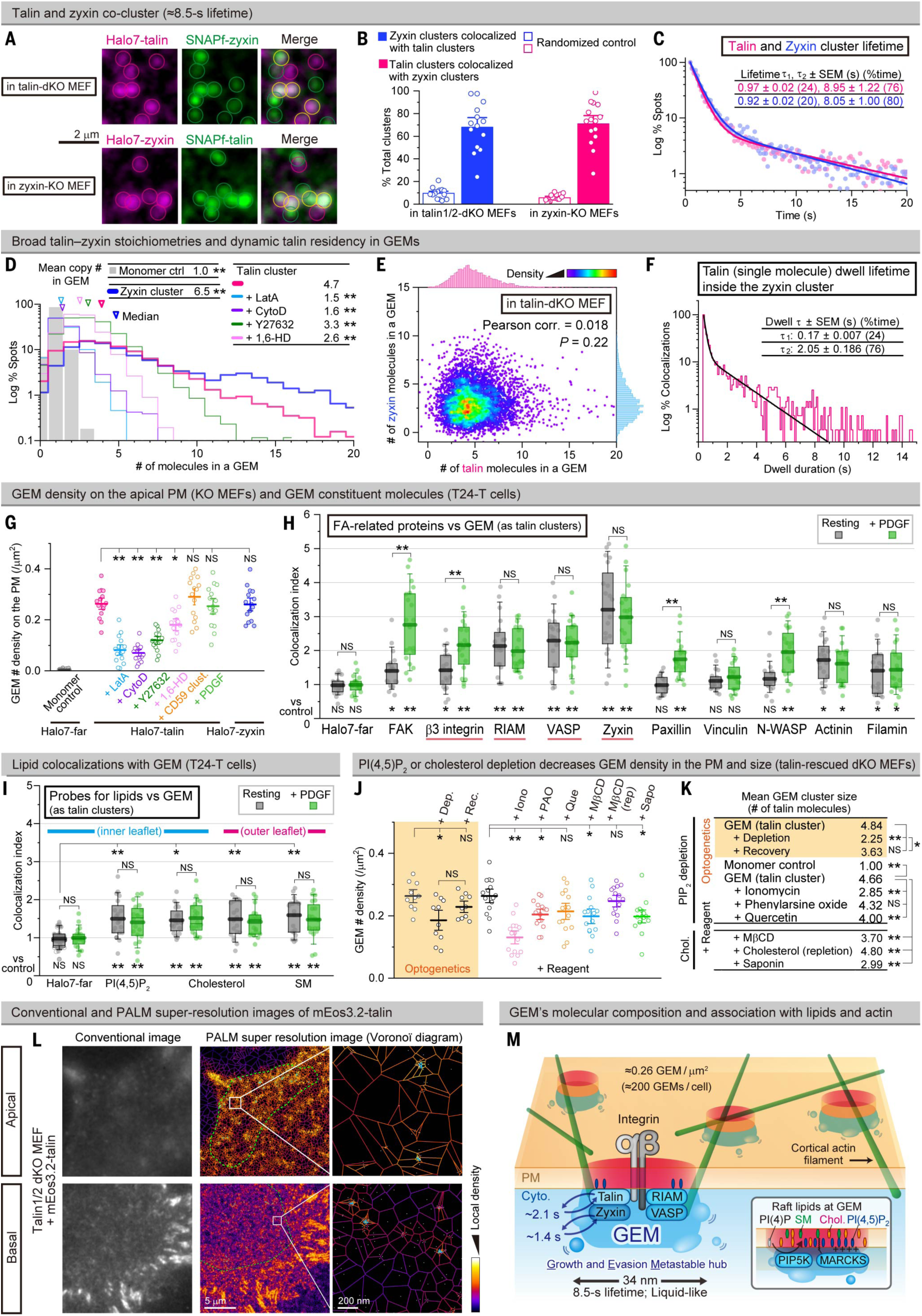
Single molecule tracking and PALM define GEM as a ≈34 nm, nanoscale, liquid-like assembly associated with PI(4,5)P_2_ and raft nanodomains. (**A and B)** Typical dual color images of talin and zyxin clusters (circles, A) and their colocalized fractions (B). (**C**) Distributions of the cluster lifetimes. (**D**) Distributions of the numbers of talin and zyxin molecules residing in GEM. (**E**) Stoichiometries of talin and zyxin in GEM. (**F**) Dwell time distributions of single talin molecules in zyxin-containing GEM. (**G**) Cluster number densities for GEMs on the apical PM (dots for individual cells; bars for mean ± SEM). (**H**) Single-molecule colocalization indexes of various FA proteins with GEM. (**I**) Single-molecule colocalization indexes of protein probes for various lipids (Halo7-tagged) with GEM (see fig. S4A for other probes). (**J and K**) GEM number densities (J) and mean talin copy numbers per GEM (K; see fig. S4, C and J for distributions) on the apical PM after the indicated treatments (talin clusters; mean ± SEM). (**L**) Conventional and PALM super-resolution images (Voronoï diagram) of GEM, visualized by mEos3.2-talin on the apical and basal PMs. The areas surrounded by the green curve (middle image) indicate regions with good focus (top; apical PM, which is tilted) or without obvious focal adhesions (bottom; basal PM) and were used for further analysis. The cyan lines in the right images indicate the smallest polygon enclosing each clustered region. (**M**) Schematic diagram showing the molecular composition of GEM and the association of GEM with PI(4,5)P_2_, raft nanodomains, and cortical actin filaments. Steel-Dwass’ test for (D (vs. talin cluster) and K), Games-Howell’s test for (G, I, and J), Two-tailed Welch’s t-test for (H and I).

A cluster-lifetime analysis revealed two lifetime components of GEMs: 22% short-lived (0.95 s) and 78% longer-lived (8.5 s) (Fig. 2C). In contrast, clusters lacking one partner, talin-only or zyxin-only clusters (hereafter “incomplete GEMs”), showed only a single short lifetime component (≈1.1 s) (fig. S3A). The longer-lived component was observed only when both talin and zyxin were present (after rescue with zyxin or talin), accounting for 78% of talin- or zyxin-containing GEMs (fig. S3A), consistent with their ≈70% colocalization. Thus talin-zyxin co-clusters are substantially more stable than incomplete GEMs.

A copy-number analysis showed broad stoichiometries for both talin and zyxin (1 to ≥20 molecules; medians 4.0 and 4.8, respectively), in contrast to the monomeric farnesylated Halo7 control (Halo7-far; median 1.0) (Fig. 2D). No correlation was observed between talin and zyxin copy numbers in individual GEMs (Fig. 2E and fig.3B).

**Fig. 3.**
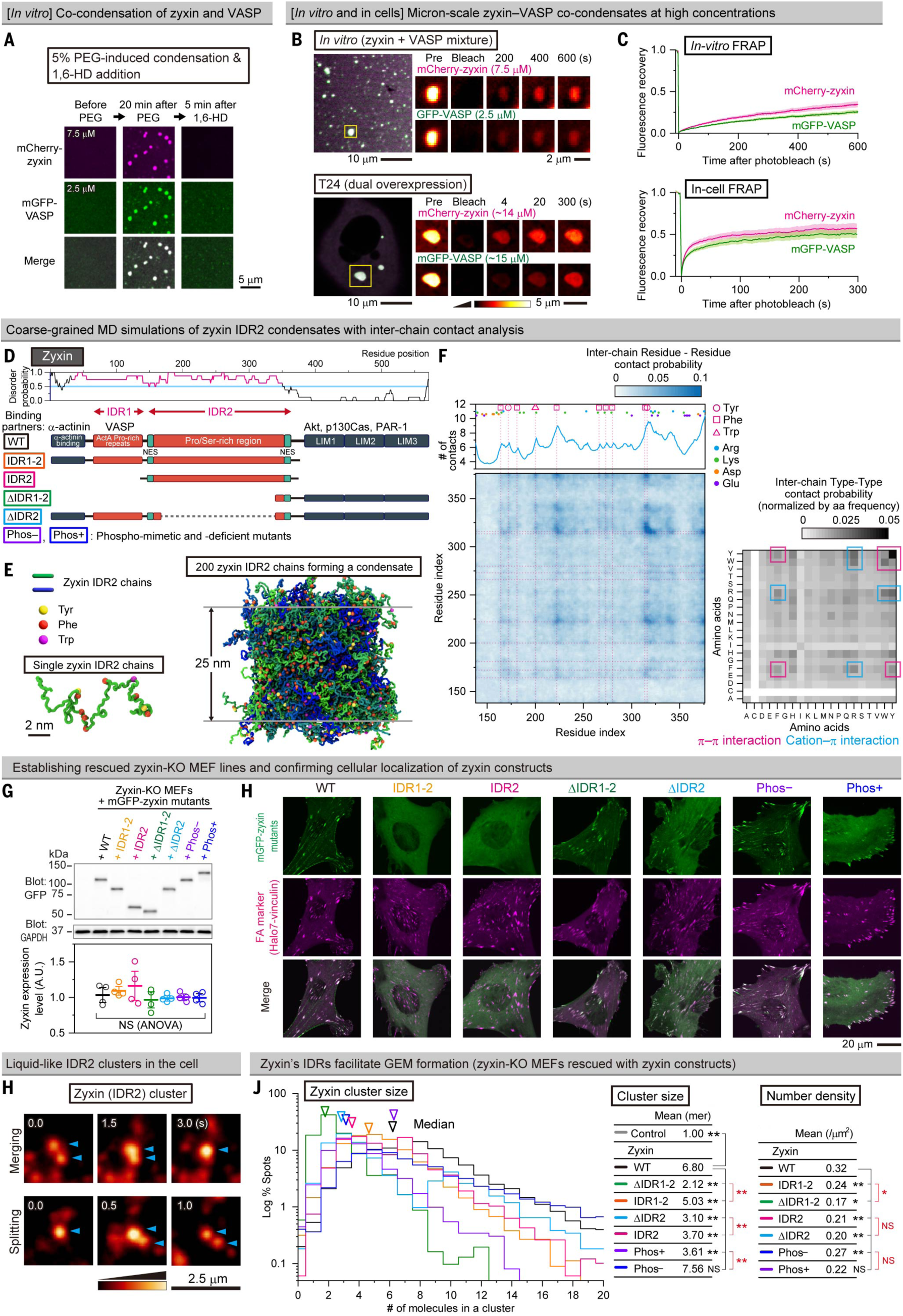
Zyxin and VASP can undergo LLPS and zyxin’s IDRs facilitate GEM formation. (**A**) Typical confocal images of *in vitro* co-condensation of purified mCherry-zyxin and mGFP-VASP in the presence of 1% PEG8K. (**B and C**) Micron-scale co-condensates of mCherry-zyxin and mGFP-VASP formed *in vitro* (top; 1% PEG8K) and in cells under overexpression conditions (bottom; T24), with FRAP indicating liquid-to-hydrogel-like behavior. (**D to F**) Coarse-grained MD simulations of zzyxin IDR2. (D) Domain structure of zyxin and their mutants employed in this study. (E) Modeling and MD simulations of a single chain and a 200-chain condensate of zyxin IDR2 at 210 K (snapshot from movie S5). (F) Inter-chain contact analysis of zyxin IDR2 condensates. (Left) Inter-chain residue-residue contact probability. (Top-left) Number of inter-chain contacts per residue. (Right) Type-type contact heatmap normalized by amino acid frequency. (**G**) Establishment of rescued zyxin-KO MEF lines expressing full-length and partial sequences of zyxin (± IDRs, mGFP-tagged) at comparable expression levels (mean ± SEM). (**H**) Typical confocal micrographs near the basal PMs of the rescued zyxin-KO MEF lines shown in G. (**H**) Typical images of nanoscale IDR2 clusters (IDR2-rescued GEM-like assemblies) undergoing merging and splitting. (**J**) Distributions of the copy numbers of zyxin (WT and mutants) per GEM cluster, determined for the rescued zyxin-KO MEF lines shown in G. One-way ANOVA for (G) and Steel-Dwass’ test for (J).

During the 8.5-s lifetime of GEMs, individual talin and zyxin molecules in GEMs rapidly exchange with their cytoplasmic pools, repeatedly entering and leaving GEMs, with dwell lifetimes in the GEM of 2.1 and 1.4 s, respectively (Fig. 2F and fig. S3C). This rapid exchange, together with highly heterogeneous stoichiometry (Fig. 2E and fig. S2B), is consistent with the notion that GEMs possess liquid-like properties.

The mean GEM density on the apical PM was ≈0.26 clusters/µm^2^ (Fig. 2G), comparable to clathrin-coated structures on the apical PM (*23*), suggesting the presence of several hundred GEMs per cell. Stimulation of CD59 or PDGFR did not alter the talin copy number per GEM or the GEM density (Fig. 2G and fig. S3D).

Partial actin depolymerization with latrunculin A and cytochalasin D, as well as myosin deactivation with the ROCK inhibitor Y27632, significantly reduced both GEM size (talin copy number per cluster) and GEM density on the apical PM (Fig. 2, D and G). These results suggest that the actin-myosin system (*24*) contributes to GEM formation and/or association with the PM.

To summarize, these data indicate that GEMs are abundant, short-lived talin–zyxin co-assemblies that exhibit heterogeneous stoichiometry and rapid molecular exchange, features consistent with metastable, liquid-like membrane hubs supported by the actin–myosin system.

### GEMs are colocalized with integrin, RIAM, VASP, α-actinin, filamin, and FAK, but not with vinculin or paxillin

We next examined the possibility that some other IAC-related proteins are incorporated into GEMs. Single-molecule colocalization analyses revealed that, in resting T24-T cells, β3 integrin, RIAM, VASP, α-actinin, filamin, and FAK colocalize with GEMs, whereas vinculin and paxillin, two major components of IACs, showed little or no detectable colocalization with GEMs (Fig. 2H and fig. S3E). Combined with the high abundance of GEMs on the apical PM (and later, also on the basal PM), these observations indicate that GEMs are distinct from classical IACs.

Upon PDGF stimulation, the single-molecule colocalization indexes of RIAM, VASP, zyxin, α-actinin and filamin with GEMs were unchanged, whereas those for integrin and FAK increased. Furthermore, paxillin and N-WASP, but not vinculin, became detectably colocalized with GEMs only after PDGF stimulation (Fig. 2H). In the following subsections, we focus on integrin, talin, RIAM, VASP, and zyxin as core GEM constituents that are shared with, yet compositionally and spatially distinct from, classical IACs.

### Essential roles of PM raft domains and PI(4,5)P_2_ in GEM formation

Because GEMs recruit the GPI-anchored receptors CD59 and Thy1, prototypical raft-associated molecules (*25–27*), we next investigated whether GEMs are associated with raft-like PM domains. Single-molecule colocalization analyses revealed that protein probes for cholesterol (D4H (*28*) for both outer and inner leaflets), sphingomyelin (OlyA-E69A (*29*) for the outer leaflet), and phosphatidylserine (the PH domain of evectin2 (*30*, *31*) for the inner leaflet) were enriched at GEMs, both before and after PDGF stimulation, to similar extents (Fig. 2I, fig. S4, A and B, and movie S4).

Perturbing raft integrity markedly affected GEMs: partial cholesterol depletion reduced both GEM number density and cluster size (talin copy number per cluster) on the apical PM, and the subsequent cholesterol repletion restored them (Fig. 2, J and K, and fig. S4C).

Because the GEM-constituent molecules talin, RIAM, and FAK can bind PI(4,5)P_2_ (fig. S3E), we examined whether GEMs and PI(4,5)P_2_ colocalize. Single-molecule colocalization analyses showed that PI(4,5)P_2_ and its biosynthetic enzyme PI5K (*32*) are enriched at GEMs both before and after PDGF stimulation (Fig. 2I, fig. S4, A and B, and movie S4). Partial PI(4,5)P_2_ depletion from the PM, achieved either optogenetically using the CIBN-CRY2 system (*33*) or chemically (*34*), reduced both the GEM number density and cluster size on the apical PM (Fig. 2, J and K, and fig. S4, D to J).

Importantly, the depletion of cholesterol or PI(4,5)P_2_ from the PM reduces not only GEM abundance on the apical PM but also the GEM cluster size (Fig. 2, J and K). These results indicate that raft domains and PI(4,5)P_2_ would contribute to GEM assembly at the PM, rather than merely tethering pre-formed GEMs to the PM, consistent with a model in which PI(4,5)P_2_-associated or enriched raft nanodomains provide a prewet membrane surface that promotes GEM formation (*35–38*).

### GEM is a nanoscale structure with a diameter of ≈34 nm

To directly determine the physical size of GEMs, we performed super-resolution photoactivated localization microscopy (PALM) imaging (*39–41*) of mEos3.2-tagged talin and zyxin on the apical PM of fixed cells (Fig. 2L and fig. S5, A to E). The PALM analysis resolved GEMs as discrete clusters containing mean copy numbers of 6.1 talin and 8.2 zyxin molecules, with a median diameter of 40.0 nm (fig. S5, F and G). These copy numbers per cluster were slightly higher than those evaluated by the live-cell single-molecule intensity analysis (Fig. 2D), probably because the PALM tessellation analysis does not detect clusters containing fewer than three molecules.

GEMs occupied the apical PM at a mean density of 0.35 clusters/µm^2^ (fig. S5H), reasonably consistent, given the different quantification methods, with the 0.26 clusters/µm^2^ measured by live-cell single-molecule imaging (Fig. 2G). A monomeric control molecule, farnesylated mEos3.2 (mEos3.2-far), did not exhibit any sign of clustering (fig. S5H).

On the *basal* PM, talin-identified and zyxin-identified GEMs with a median diameter of 28.5 nm were detected (fig. S5F) outside obvious classical/conventional IACs (see the bottom-middle panel in Fig. 2L). Taking the arithmetic mean of the median diameters measured on the apical and basal PMs, we estimate the characteristic GEM diameter to be ≈34 nm, as a representative value across the two membrane contexts (Fig. 2M).

### Zyxin and VASP show intrinsic capacities to undergo LLPS

GEMs exhibit liquid-like features, including a wide-range of molecular composition stoichiometries (Fig. 2E and fig. S3B), short lifetimes, and rapid molecular exchange (Fig. 2, C and F). These properties are frequently implicated in condensates generated through liquid-liquid phase separation (LLPS). Furthermore, zyxin, VASP, and RIAM, the main structural constituents of GEMs, contain long intrinsically disordered regions (IDRs; fig. S6) (*42*), which are frequently involved in LLPS (*43*, *44*). Various RTK cytoplasmic tails also contain IDRs (e.g., an 88-aa IDR in PDGFR). Although GEMs are nanoscale assemblies, rather than conventional micron-scale LLPS condensates, LLPS-like behavior at the nanoscale has been reported (*45–50*). Furthermore, GEM formation depends on raft domains and PI(4,5)P_2_ (Fig. 2, I to K), features consistent with the idea that LLPS can be enhanced on the membrane surface. Therefore, we asked whether an LLPS-like process underlies GEM formation, potentially aided by raft nanodomains, PI(4,5)P_2_, and the cortical actin meshwork, which could provide a permissive environment for LLPS-driven condensation of IDR-containing proteins (*35*, *37*, *38*, *51*).

We first tested the intrinsic LLPS capacities of zyxin and VASP *in vitro*. Purified mCherry-zyxin and mGFP-VASP, at concentrations of 2.5-7.5 µM, formed micron-scale condensates, alone and in combination, in the presence of 1% (w/w) polyethyleneglycol (PEG) (Fig. 3A and fig. S7A). These condensates exhibited liquid-hydrogel-like FRAP behavior (Fig. 3, B top and C top, and fig. S7, B and C) and were sensitive to 1,6-hexanediol (Fig. 3A), consistent with LLPS.

To further examine whether zyxin’s IDRs can drive condensation, we performed coarse-grained (CG) molecular dynamics simulations of a partial zyxin chain (IDR2; Fig. 3, D and E, and movie S5). The simulations revealed temperature-dependent condensate formation by IDR2 (fig S8, A to C). Contact analysis indicated that ρ-ρ and cation-ρ interactions, primarily involving Phe, Tyr, Trp, and Arg residues, play key roles in condensate formation (Fig. 3F), consistent with a model in which aromatic and positively charged residues provide dominant interaction motifs stabilizing IDR2 condensates (*52*).

We next asked whether zyxin and VASP can form condensates in cells. Under overexpression conditions in T24 cells, both zyxin and VASP formed micron-scale, liquid-like condensates (Fig. 3, B bottom and C bottom, and fig. S9). Estimated cytoplasmic concentrations were ≈13 µM (zyxin) and ≈15 µM (VASP), compared with endogenous 0.73 and 0.63 µM (3.11 and 0.25 µM in a MEF line, respectively; fig. S10). In contrast, over-expressed RIAM failed to form detectable micron-scale condensates, even at a higher intracellular concentration (≈18.4 µM, fig. S9A and fig. S10E).

These results obtained *in vitro*, in silico, and in cells under overexpression conditions demonstrate that zyxin and VASP possess intrinsic capacities to undergo LLPS, whereas RIAM does not. The LLPS capability of VASP is consistent with previous reports (*53*, *54*). These properties provide a physical basis for how zyxin and VASP could drive the formation of nanoscale, liquid-like assemblies at physiological expression levels.

### In cells, under physiological expression conditions, zyxin and VASP form nanoscale liquid-like assemblies rather than micron-scale condensates

Although zyxin at endogenous concentrations forms micron-scale condensates *in vitro* in the presence of 1% PEG, zyxin expressed at endogenous levels does not form micron-scale condensates in living cells (fig. S11, A and B). Instead, zyxin expressed at endogenous levels in cells assembles into *nanoscale* structures corresponding to GEMs, which undergo dynamic merging and splitting (fig. S11C). Furthermore, as found in talin-identified GEMs, zyxin-identified GEMs decrease in both size and number density on the apical PM upon treatment with 1,6-hexanediol (Fig. 2, D and G), exhibit rapid subsecond-to-second exchange of both recruited signaling molecules and zyxin with the cytoplasmic pools (Fig. 1, D and H, and fig. S3C; see also Fig. 2F for talin-identified GEMs), and show highly flexible stoichiometries of talin and zyxin copy numbers (fig. S3B; see also Fig. 2E).

Collectively, these behaviors suggest that GEMs possess liquid-like properties. We therefore describe the GEM formation process (by LLPS-capable proteins at endogenous concentrations) as an “LLPS-like process” defined here as a nanoscale liquid-like co-assembly process driven by multivalent, weak but specific interactions in cells, analogous in physical principle to micron-scale LLPS (*48*, *55*). Submicron-scale condensates of LLPS-capable proteins under subsaturation concentrations have been reported both *in vitro* and in cells (*45–47*, *49*), whereas nanoscale condensates as small as GEMs, with clearly defined signaling functions, remain poorly characterized.

While signal-induced, large, micron-scale, long-lived, LLPS-driven signaling assemblies have been reported in *in vitro* systems containing very high protein concentrations or in cells overexpressing relevant molecules (*56–59*), those structures differ from GEMs in spatiotemporal scales and formation regime. GEMs are substantially smaller (≈34 nm) and short-lived (≈8.5 s), form at physiological expression levels in cells, and exist constitutively in cells even before stimulation.

### Zyxin’s IDR is critical for GEM formation through LLPS-like processes

We hypothesized that zyxin IDRs, particularly IDR2, are major driving elements for GEM assembly (fig. S12A), given that talin lacks IDRs (fig. S6), RIAM does not form condensates in cells (fig. S9A and fig. S10E), and endogenous VASP is expressed at substantially lower levels than zyxin (fig. S10D).

First, the isolated zyxin IDR2 forms submicron-scale condensates *in vitro* at 3 µM in the presence of 1% PEG, similar to the full-length zyxin, whereas the ΔIDR2 construct does not (fig. S12B).

Second, in T24-T cells, zyxin IDR constructs exhibited significantly higher colocalization with GEMs than their IDR-deleted counterparts (fig. S12C). A similar trend was observed for VASP constructs (fig. S12, A and C).

Third, in five independent zyxin-KO MEF cell lines rescued with physiological levels of partial and full-length zyxin constructs (Fig. 3G), confocal imaging of basal optical sections revealed that full-length zyxin, ΔIDR1-2, and ΔIDR2 localize to IACs and diffusely in the basal PM region, whereas IDR1-2 and IDR2 showed no detectable localization at IACs (Fig. 3H). These results are consistent with single-molecule colocalization analyses showing the preferential association of IDR-containing zyxin constructs with GEMs on the PM (fig. S12C). Thus, zyxin’s IDR1-2 and IDR2 are selectively incorporated into GEMs, but not into IACs.

Fourth, in zyxin-KO MEFs, expressed IDR2 forms nanoscale, liquid-like assemblies in the GEM size range that undergo merging and splitting (Fig. 3H), similar to complete GEMs (fig. S11C), although they contain approximately half the copy number of full-length zyxin per GEM cluster (Fig. 3J). Nevertheless, IDR2 rescue produced significantly larger clusters than ΔIDR rescue (Fig. 3J). These results suggest that IDR2 might be a major driver of LLPS-like processes and liquid-like properties of GEMs.

Finally, rescue experiments using phospho-deficient and -mimetic zyxin mutants indicated that phosphorylation lowers zyxin’s ability to colocalize with or form GEMs (Fig. 3, C, H, and J, and fig. S12C).

Taken together, these findings demonstrate that zyxin’s IDR2 is necessary for GEM formation through LLPS-like processes. Moreover, in zyxin-KO cells, where other endogenous GEM components are present, zyxin’s IDR2 is sufficient to drive the condensation step that yields nanoscale, liquid-like clusters corresponding to GEMs. These results also indicate that IDR2 is a main cause of GEM liquidity. Importantly, zyxin’s IDR2 preferentially partitions into GEMs rather than IACs (Fig. 3, H and J, and fig. S12C). This selective GEM-rescue strategy using IDR constructs, particularly IDR2, was used extensively in subsequent experiments.

### GEMs recruit growth and survival signaling molecules

We next examined whether GEMs recruit downstream cytoplasmic signaling proteins implicated in growth signaling. After PDGF stimulation, the single-molecule colocalization index analysis showed recruitment of PLCγ1 to GEMs (Fig. 4A), with a dwell lifetime of ≈80 ms (Fig. 1H). FAK was detectably colocalized with GEMs even before stimulation, and its colocalization was strongly enhanced upon PDGF stimulation (Fig. 4A) with a sub-second dwell lifetime of ≈200 ms (Fig. 4, B and C, and movie S4).

**Fig. 4.**
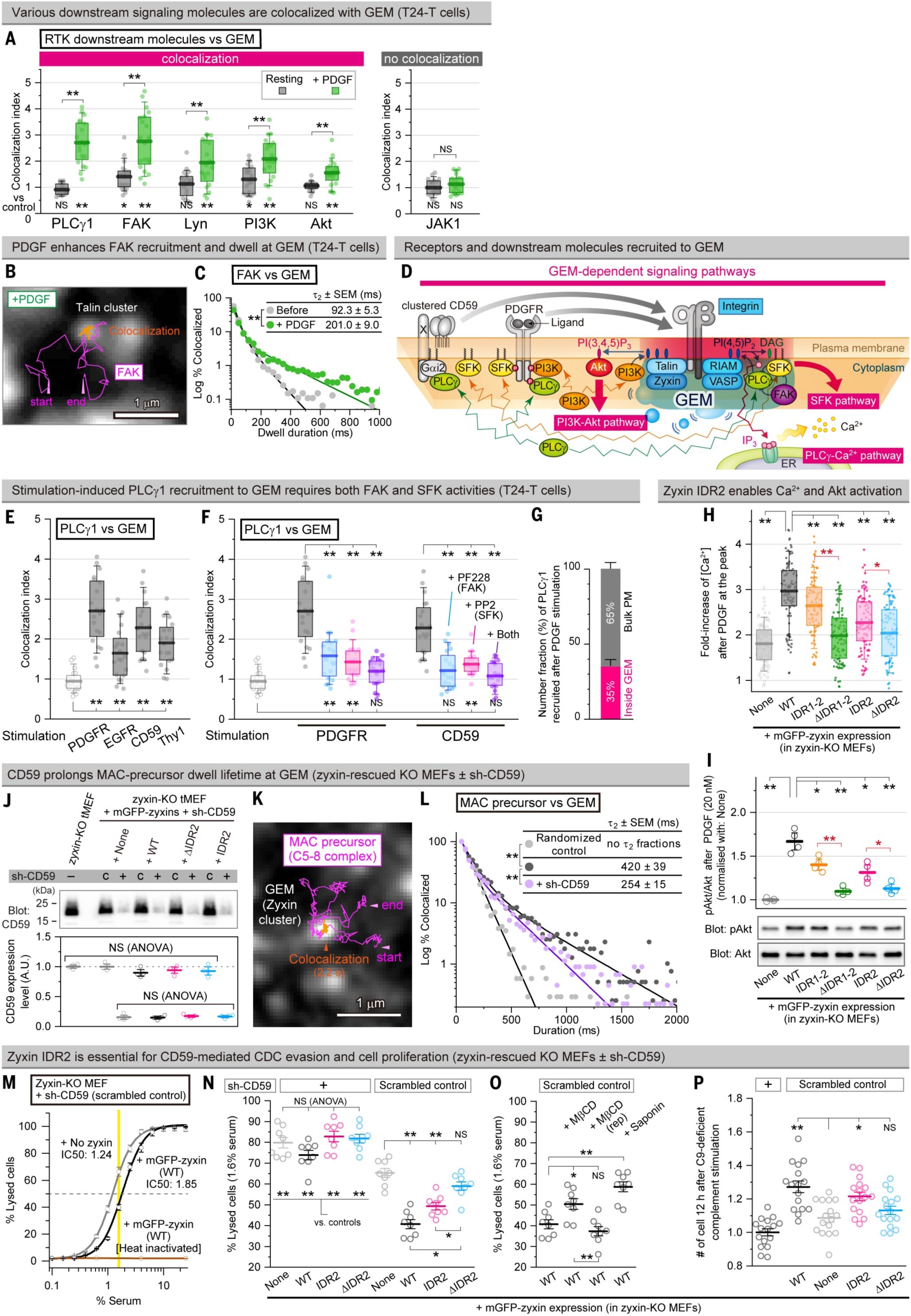
GEM functions as a signaling platform for both RTK and GPI-anchored receptor signaling pathways. (**A**) Single-molecule colocalization indexes of the PDGFR downstream signaling molecules (Halo7-tagged) with GEM. (**B and C**) Typical image showing transient colocalization of Halo7-FAK (trajectories) with GEM (white puncta) after PDGF stimulation (B, snapshot from movie S4). Distributions of the dwell durations of FAK at GEM before and after PDGF stimulation (C). (**D**) Schematic summarizing receptors and downstream signaling molecules recruited to GEM. (**E**) Stimulations of RTKs and GPI-anchored receptors induce PLCγ1 recruitment to GEM. (**F**) Inhibition of either FAK or SFK suppresses PLCγ1 recruitment to GEM after PDGFR stimulation. (**G**) Fraction of PLCγ1 recruited to GEM vs. non-GEM regions of the apical PM after PDGF stimulation (mean ± SEM). (**H and I**) IDR expression restores PDGF-induced Ca^2+^ mobilization (fold increase at the peak, **H**) and Akt activation (S473 phosphorylation; mean ± SEM; I). (**J**) Sh-CD59-mediated CD59 knockdown in zyxin-KO MEFs, reducing the CD59 expression to ≈15%. Typical western blot pattern (top) and quantification (bottom, mean ± SEM). (**K and L**) Transient recruitment of MAC precursors (trajectory; K) to GEM (white puncta) and their dwell lifetime at GEM (L). (**M to O**) CDC assay as a function of serum concentration (mean ± SEM, M). The 1.6% normal serum condition (yellow line in M) was used for the comparison in N and O (mean ± SEM) to maximize the dynamic range. (**P**) Cell growth after stimulation with C9-depleted serum (mean ± SEM). Two-tailed Welch’s t-test for (A, H), Games-Howell’s test for (E, F, H, I, and N to P), one-way ANOVA for (J and N), and Brunner-Munzel test for (L).

These results indicate that GEMs intermittently and repetitively recruit PLCγ1 and FAK molecules, one molecule after another, resulting in multiple sequential recruitment events. This intermittent, serial recruitment mirrors the subsecond-to-second dwell behavior observed for engaged receptors and core GEM constituents (Figs. 1D and 2F, and fig. S3C).

What signaling output can occur during such brief residence of a single signaling molecule at GEMs? For PLCγ1 (mean dwell ≈80 ms; Fig. 1H), the reported maximal catalytic turnover rate (≈230/s) (*60*, *61*) provides an upper-bound estimate that each recruited PLCγ1 molecule could hydrolyze ≈20 PI(4,5)P_2_ molecules enriched at GEMs (Fig. 2I), thereby generating ≈20 IP_3_ molecules within ≈80 ms. Because multiple PLCγ1 molecules are repeatedly recruited to GEMs, GEMs are expected to generate frequent IP₃ bursts. The temporal summation of these burst sequences would drive robust Ca²⁺ mobilization.

Single-molecule colocalization further revealed that Lyn (an SFK; hereafter, SFK for general discussion and Lyn for this particular kinase), PI3K, and Akt, prototypical kinases involved in cell survival and growth, were also recruited to GEMs after PDGF stimulation (Fig. 4A). PI3K is localized at GEMs even before stimulation, and its colocalization was enhanced upon PDGF stimulation (Fig. 4A). Among these kinases, Akt activation is strongly associated with cell-survival signaling (*62*). In contrast, JAK1 did not show detectable colocalization with GEMs either before or after PDGF stimulation (Fig. 4A right). Taken together, these results indicate that GEMs function as hubs that preferentially recruit multiple receptors and downstream signaling molecules involved in cell growth and survival (Fig. 4D).

PLCγ1 recruitment to GEMs was also triggered by the activation of another RTK, EGFR, and by the stimulation of another GPI-anchored receptor, Thy1 (Fig. 4E). These results suggest that GEM-associated PLCγ1 recruitment can be induced downstream of multiple RTKs and GPI-anchored receptors.

Mechanistically, PLCγ1 recruitment to GEMs after PDGFR or CD59 stimulation was markedly suppressed by the inhibition of either FAK or SFK, and was nearly abolished when both kinases were inhibited (Fig. 4F). These findings suggest that PLCγ1 is recruited to proteins that are tyrosine-phosphorylated by FAK and/or SFK at GEMs, including FAK and SFK themselves and other GEM-associated substrates.

Because PLCγ is known to be directly recruited to activated/phosphorylated RTKs, we quantified the relative contributions of GEM-dependent versus GEM-independent pathways to PLCγ1 recruitment to the apical PM after PDGF stimulation. Approximately 35% of PLCγ1 molecules arriving at the apical PM colocalized with GEMs in a FAK/SFK-dependent manner, whereas ≈65% localized outside GEMs (Fig. 4G), consistent with direct binding to activated RTKs reported previously (*1*). Thus, PLCγ1 likely reaches the PM through two parallel routes: (1) direct binding to activated RTKs outside GEMs (≈65%), and (2) binding to GEM-localized proteins tyrosine-phosphorylated by activated PDGFR, FAK, and/or SFK (≈35%).

In zyxin-KO cells, where complete GEMs are depleted, Ca^2+^ mobilization upon PDGF stimulation was markedly reduced and restored by the re-expression of full-length zyxin. Importantly, Ca^2+^ mobilization was partially rescued by the expression of IDR1-2 or IDR2, but not by the ΔIDR1-2 or ΔIDR2 construct (Fig. 4H). PDGF-induced Akt activation was similarly impaired in zyxin-KO cells and rescued by the expression of IDR1-2 or IDR2, but not by the ΔIDR constructs (Fig. 4I).

These results show that zyxin IDR-driven GEMs are required for the efficient activation of these signaling pathways. Moreover, the selective rescue by zyxin IDR constructs point to an important contribution of GEM’s liquid-like organization to signaling, including the capacity to dynamically recruit RTKs, GPI-anchored receptors, and diverse downstream signaling molecules.

### GEMs enhance CD59-mediated immune evasion and cell growth

To examine the involvement of GEMs in CD59-mediated immune evasion, we generated a zyxin-KO MEF line in which CD59 expression was reduced to ≈15% of parental levels (Fig. 4J). Upon rescue with full-length zyxin, MAC precursors were transiently and intermittently recruited to GEMs, as observed in T24-T cells, even at reduced CD59 expression. Furthermore, the presence of CD59 prolonged MAC-precursor dwell lifetimes at GEMs (Fig. 4, K and L). Accordingly, GEMs increase the local opportunity for CD59–MAC precursor binding and prolong MAC precursor residence at GEMs.

We next examined whether GEMs enhance CD59-mediated cell protection from CDC, using cell-lysis assays (Fig. 4M). Zyxin-KO MEFs rescued with full-length zyxin exhibited a ≈1.5-fold increase in the IC_50_ for non-heat-inactivated serum (Fig. 4M). Rescue with full-length zyxin or IDR2, but not with ΔIDR2, significantly enhanced the CDC resistance compared with non-rescued cells (Fig. 4N). The CDC protection was abolished in CD59-knockdown cells, regardless of zyxin rescue (Fig. 4N), confirming CD59 dependence. Importantly, cholesterol depletion, which reduces GEM number and size despite the presence of all GEM protein components, markedly diminished CD59-mediated CDC protection, and the subsequent cholesterol repletion restored it (Fig. 4O). These results link raft-domain-dependent GEM integrity (number/size) to CD59-mediated immune evasion.

Rescue with full-length zyxin or IDR2, but not with ΔIDR2, also enhanced MAC-precursor-induced, CD59-dependent cell proliferation (Fig. 4P; see materials and methods), in line with the recruitment of PLCγ1, a signaling molecule involved in cell growth, to CD59-resident GEMs (Fig. 1, G to I) and with our previous results (*4*, *5*). Thus, CD59 signaling at GEMs contributes not only to CDC resistance but also to proliferative signaling (Figs. 1L and 4D).

Because zyxin IDR1-2 and IDR2 do not localize to IACs (Fig. 3H), the functional recoveries observed here (Fig. 5, H, I, and N to P) can be attributed to the restoration of GEMs rather than IACs. Together, these data show that GEMs greatly enhance both CD59-dependent immune evasion via CDC resistance and growth signaling, probably by confining signaling molecules within a nanoscale liquid-like environment, consistent with the requirement for IDR2 in GEM formation.

**Fig. 5.**
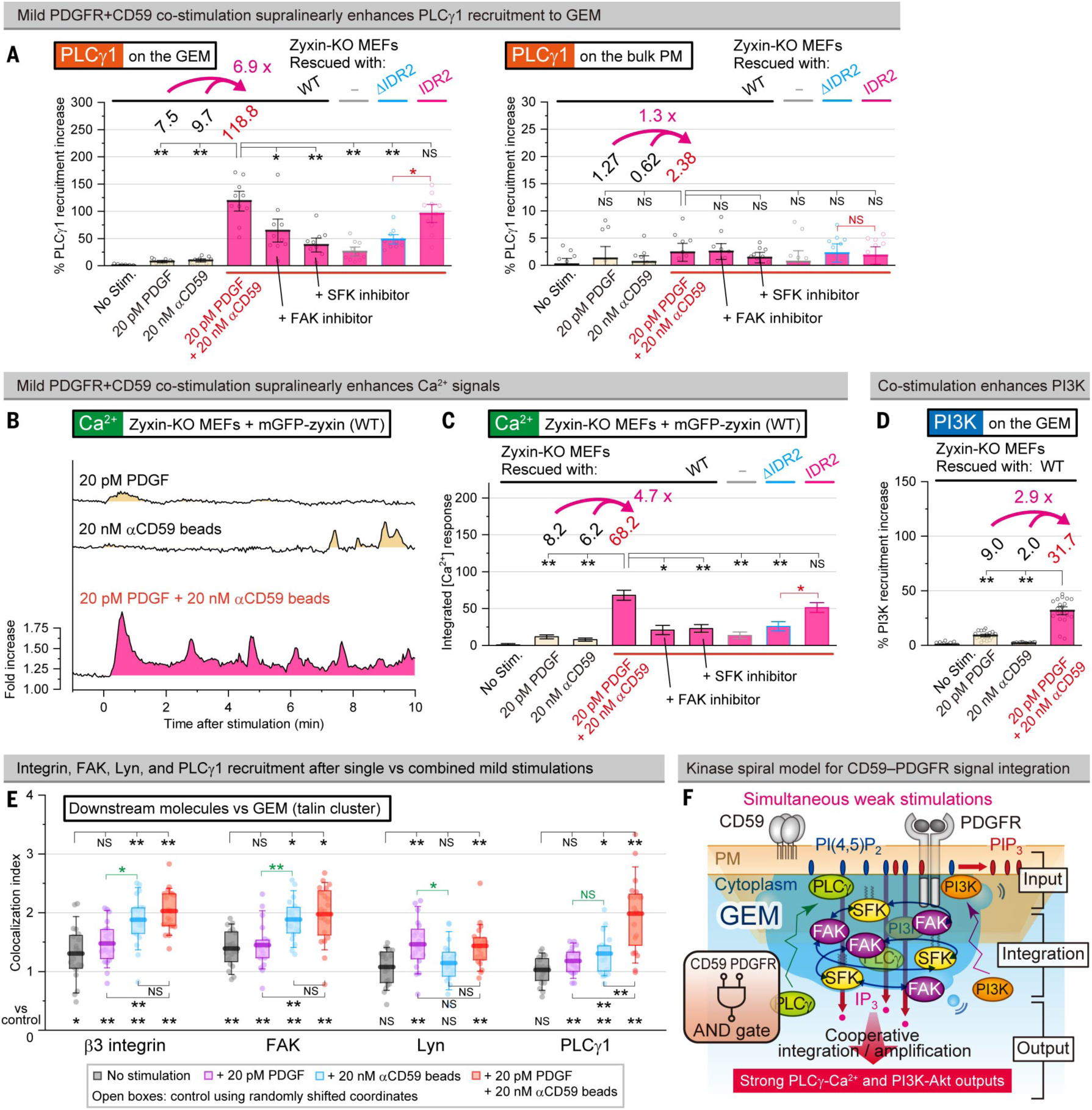
GEM cooperatively integrates CD59 and PDGFR signaling, generating supralinearly amplified outputs. (**A**) Percent increases in PLCγ1 recruitment frequency to GEM (left) and to non-GEM regions of the PM (right) after mild individual stimulations and co-stimulation of CD59 and PDGFR, in zyxin-KO MEFs rescued with full-length zyxin, IDR2, and ΔIDR2 (mean ± SEM). Mild stimulation conditions were 20 pM PDGF and 20 nM αCD59-coated beads. Twenty pM PDGF approximates physiological serum levels (*106*), and corresponds to 1/1,000 of our standard PDGF stimulation; 20 nM αCD59 beads corresponds to 1/100 of the standard stimulation. (**B**) Typical Ca^2+^ mobilization time courses in individual cells after mild individual and simultaneous stimulations. (**C**) Total Ca^2+^ signal (areas under the curve; AUC) (mean ± SEM). (**D**) Percent increases of the PI3K recruitment frequency to GEM (mean ± SEM). (**E**) Single-molecule colocalization indexes of Halo7-tagged β3 integrin, FAK, Lyn, and PLCγ1 with GEM (SNAPf-talin clusters) after mild single stimulations and co-stimulations of PDGFR and CD59. (**F**) Schematic illustrating the kinase spiral model, in which reciprocal FAK-SFK chain activation drives cooperative and supralinear amplification of the GEM outputs upon CD59+PDGFR co-stimulations. Games-Howell’s test for (A, C, D, and E) and two-tailed Welch’s *t*-test for (E).

### RTK and CD59 signals are physically integrated and cooperatively amplified at GEMs

Because CD59 can trigger both CDC resistance and cell-growth signaling, whereas PDGFR is a canonical growth factor receptor, we asked how cells respond to simultaneous CD59 and PDGFR stimulation. We applied concurrent CD59 and PDGFR stimulations to zyxin-KO MEFs rescued with full-length zyxin, IDR2, or ΔIDR2, as well as to non-rescued cells.

In cells rescued with full-length zyxin, under mild, physiologically-relevant stimulation conditions, co-stimulation increased PLCγ1 recruitment to GEMs to ≈6.9-fold above the arithmetic sum of responses to the individual stimulations (Fig. 5A, left). This supralinear increase was restricted to GEMs, with no corresponding enhancement detected outside GEMs on the PM (Fig. 5A, right). Similar co-stimulation-dependent supralinear responses were observed for Ca^2+^ mobilization (Fig. 5, B and C) and PI3K recruitment to GEMs (Fig. 5D). Thus, co-stimulation drives GEM-restricted supralinear amplification of PLCγ1/PI3K recruitment and Ca^2+^ signaling, pointing to cooperative integration of RTK- and CD59-derived inputs at GEMs.

Under mild single stimulations, CD59 preferentially induced FAK recruitment, whereas PDGF preferentially induced Lyn (SFK) recruitment (Fig. 5E; note that strong PDGF stimulation induced the recruitment of both FAK and Lyn, as shown in Fig. 4A). Supralinear amplification required both kinases: inhibiting either FAK or SFK abolished the enhanced PLCγ1 recruitment and Ca^2+^ mobilization upon co-stimulation (Fig. 5, A and C). These findings indicate a cooperative mechanism in which reciprocal FAK-SFK activation supralinearly increases phosphotyrosine-bearing docking sites in GEMs, thereby boosting the recruitment of PLCγ1 and PI3K (see Fig. 4F) (*62*, *63*).

Given their nanoscale diameter (≈34 nm) and liquid-like environment, GEMs will increase the encounter frequency between FAK and SFKs, facilitating reciprocal chain phosphorylation (activation) (*64–66*), a process referred to as a “kinase spiral”. To probe this possibility, we performed independent tests using the binder-tag system (*67*) and the optoDroplet system (*68*, *69*). The results supported a confinement-based mechanism (fig. S13). Thus, these results together position GEMs as nanoscale integrative platforms in which RTK- and CD59-derived inputs are converted into supralinearly amplified PLCγ1-IP_3_-Ca^2+^ and PI3K-PI(3,4,5)P_3_-Akt outputs that are central to cell survival and growth (Fig. 5F).

Importantly, IDR2-rescued GEMs fully restored the co-stimulation-induced cooperative supralinear amplification of PLCγ1 recruitment and Ca^2+^ signaling, whereas ΔIDR2 did not (Fig. 6, A and C), demonstrating that the IDR2-driven liquid-like GEM organization (Fig. 3H) is required for GEM’s integrative signaling function under physiological expression and stimulation conditions.

**Fig. 6.**
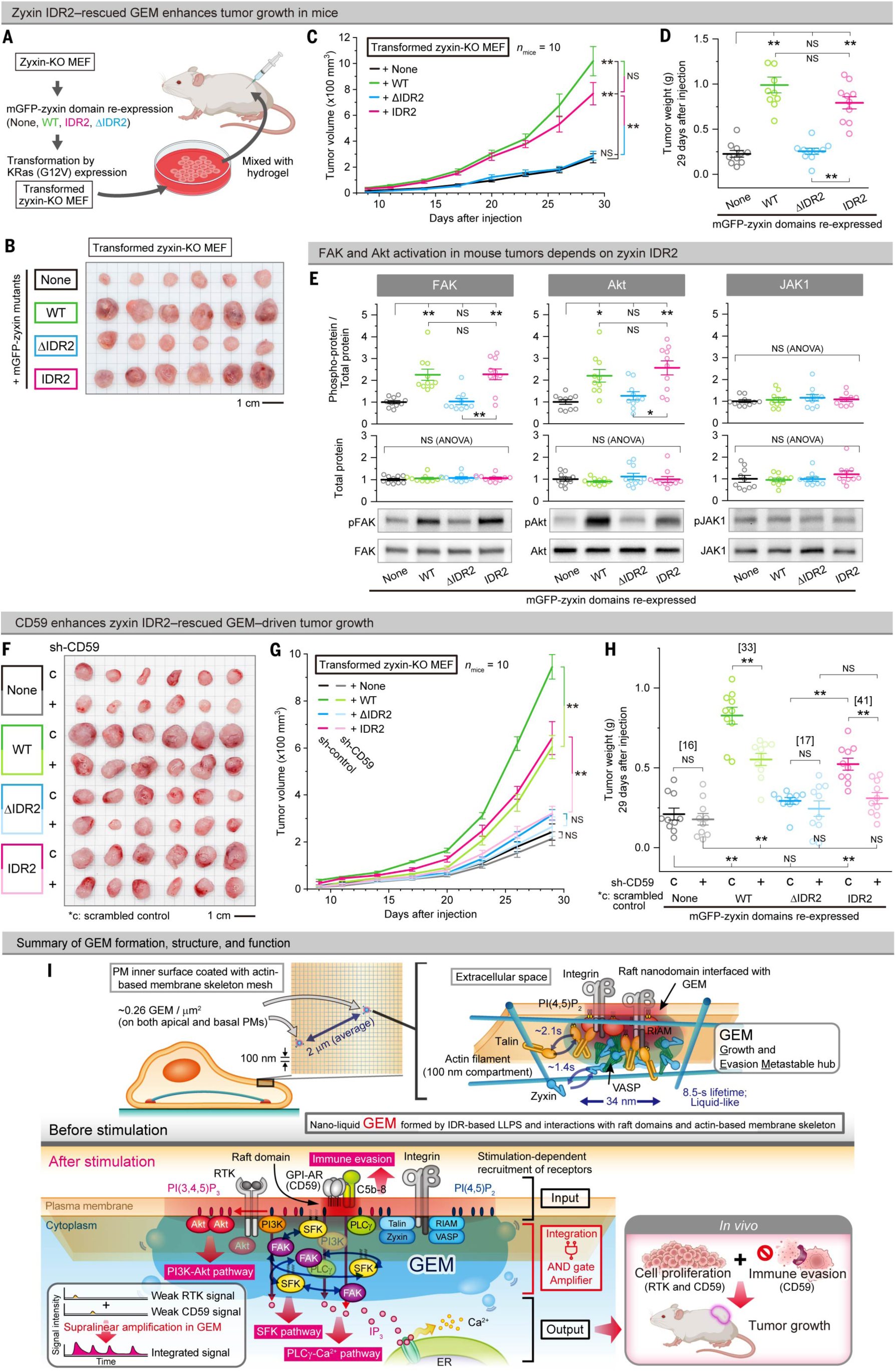
GEM promotes tumor growth in mice. (**A**) Schematic of the xenograft workflow. (**B to D**) Tumor growth of transformed zyxin-KO MEFs is enhanced by GEM restoration. (B) Images of excised tumors. (C) Tumor growth curves (mean ± SEM). (D) Weights of excised tumors at day 29 after cell injection (mean ± SEM). (**E**) Western blot analysis of FAK, Akt, and JAK1 phosphorylation (pY397, pS473, and pY1034/1035, respectively) in excised tumors (mean ± SEM). (**F to H**) CD59 dependence of GEM-supported tumor growth. (F) Images of excised tumors. (G) Tumor growth curves (mean ± SEM). (H) Weights of excised tumors. Games-Howell’s test for (C, D, E, G, and H), one-way ANOVA for (E), and two-tailed Welch’s *t*-test for (H). (**I**) Model summarizing GEM organization and function. (**Top**) GEM (Growth and Evasion Metastable hub) assemblies, mainly consisting of integrins, talin, RIAM, VASP, and zyxin (Fig. 2H), existing at a number density of ≈0.26 clusters/µm^2^ (Fig. 2G). Given that the cortical-actin-induced PM compartment size is ≈100 nm in T24 cells (*107*), approximately one GEM is expected per ≈400 compartments. GEMs are ≈34 nm nanoscale, liquid-like protein assemblies (fig. S5F) that constitutively form and disperse with a lifetime of ≈8.5 s on both the apical and basal PMs (outside the IACs) (Fig. 2C). Constituent molecules are exchanging rapidly (<2 s; Figs. 2F, 4C, and fig. S3C). Liquid-like behavior is consistent with multiple observations, including broad stoichiometries (Fig. 2E and fig. S3B), rapid exchange rates (Figs. 2F, 4C, and fig. S3C), merging and splitting (fig. S11C), and the requirement of zyxin’s IDR for formation and function (Fig. 3 and fig. S12). GEM assembly is enabled by a permissive, prewet membrane environment provided by the actin-based membrane-skeleton meshwork (*23*, *108*) (Fig. 2, D and G) and PI(4,5)P_2_-containing raft nanodomains (Fig. 2, I to K, and fig. S4), which together furnish a nanoscale surface that supports condensation and co-assembly. (**Bottom**) Upon stimulation, engaged GPI-anchored receptors and RTKs are transiently recruited to GEM, which in turn induces further recruitment of receptors’ downstream signaling molecules (Figs. 1, C to E, and 4). Nano-liquid confinement within GEM increases the encounter frequency among recruited molecules within GEM, and interactions would occur extremely efficiently. GEM serves as a hub for CD59-driven proliferative signaling (Fig. 4P), consistent with prior work (*4–6*), and for CD59-mediated immune evasion by recruiting both CD59 and MAC precursors (C5b-8, Figs. 1K and 4L). Upon co-stimulation of CD59 and RTK, FAK (CD59 pathway) and SFKs (RTK pathway) are co-recruited (Fig. 5E), promoting reciprocal chain phosphorylation (kinase spiral; Fig. 5F and fig. S13), which induces the supralinear recruitment of PLCγ and PI3K to GEM (Fig. 5, A and D). Thus, GEM functions as an AND gate (coincidence detector) for CD59 and RTK inputs (Fig. 5F). *In vivo*, GEM competence correlates with the elevated activation of FAK and Akt, critical molecules for cell survival and growth (Fig. 6E), and is required for robust tumor growth (Fig. 7, A to D and F to H).

### Tumor growth in mice is promoted by GEM, with a requirement for zyxin’s IDR2

To test whether GEMs contribute to tumor growth *in vivo*, we established a K-RasG12V-driven tumor model using zyxin-KO MEFs. The parental zyxin-KO MEF line, immortalized by serial passaging, did not form solid tumors in mice (Fig. 6A and fig. S14A). After K-RasG12V transformation, zyxin-KO MEFs formed solid tumors only slowly in mice (nude mice; BALB/cAJcl-Foxn1nu) after subcutaneous injection, whereas rescue with full-length zyxin produced significantly larger and faster-growing tumors (Fig. 6, B to D). IDR2 rescue phenocopied full-length zyxin rescue, whereas ΔIDR2 rescue did not (Fig. 6, B to D). Because IDR2 shows minimal incorporation into IACs (Fig. 3H), these tumor phenotypes are attributable to the restoration of GEMs, rather than the recovery of IACs. Meanwhile, zyxin-KO MEFs still formed tumors, albeit slowly (Fig. 6, B to D), indicating that GEMs are not the sole driver of proliferation. Instead, GEMs provide a substantial quantitative fitness advantage within a redundant pro-survival network *in vivo*.

Tumors derived from full-length zyxin- or IDR2-rescued cells exhibited elevated phosphorylations of FAK and Akt, whereas ΔIDR2-rescued tumors did not (Fig. 6E). JAK1 phosphorylation remained uniformly low across all tumors generated from all four rescued cell lines (Fig. 6E), consistent with the lack of detectable JAK1 recruitment to GEMs in cultured cells (Fig. 4A). Together, these findings indicate that GEM-linked FAK-SFK signaling and PI3K-Akt activation, defined in cultured cells (Fig. 4A), are also engaged in tumor growth *in vivo*.

To examine generality beyond MEFs, we generated zyxin-KO clones in the human cancer cell lines RKO (colon) and MDA-MB231 (breast) and re-expressed endogenous levels of full-length zyxin, IDR2, or ΔIDR2 (fig. S14B). Both the *in vitro* proliferation (fig. S14C) and tumor growth in mice (RKO; fig. S14, D to F) showed the same IDR2 dependence as in MEFs, supporting a broad requirement for zyxin’s IDR2, and thereby IDR2-dependent GEM formation, for efficient tumor growth.

Because nude mice lack T cells but retain intact innate immunity, including complement (*70*), CD59 on grafted cells can be activated by MAC precursors *in vivo*. To assess the contributions of CD59-mediated immune evasion and proliferative signaling *in vivo*, we employed the four transformed MEF rescue lines and reduced CD59 expression by ≈85%, using CD59 shRNA (Fig. 4J). An *in vivo* analysis showed that functional GEMs (full-length zyxin or IDR2 rescue) support tumor growth even with ≈85% CD59 knockdown, whereas CD59 expression further accelerates tumor progression (Fig. 6, F to H), consistent with combined contributions of CD59-dependent proliferative signaling and complement evasion.

The GEM-driven, CD59-independent growth signaling *in vivo* is likely mediated in part by RTK-linked Akt activation: *in vivo*, Akt phosphorylation is elevated in GEM-competent tumors (Fig. 6E), and in cultured cells, Akt is recruited to GEMs upon PDGF stimulation, but not upon CD59 stimulation (Fig. 4A and fig. S14G), suggesting that RTK-dependent Akt activation is amplified, at least in part, via GEMs.

As control experiments, we examined whether CD59 knockdown affects the proliferation of the four MEF lines under standard *in vitro* culture conditions, where the complement activity is eliminated by heat-inactivation. The CD59 knockdown had no detectable effect on the proliferation of any of the four MEF lines (Fig. S14H), indicating that the CD59-dependent effects become biologically relevant specifically *in vivo*, where complement is active.

In conclusion, these findings demonstrate that GEMs are major contributors to tumor growth in mice, boosting growth-factor signaling and CD59-associated *in vivo* advantages. The recurring requirement for zyxin IDR2 across tumor models connects these *in vivo* phenotypes to an IDR2-dependent, liquid-like GEM organization characterized *in silico* and in cultured cells (Fig. 3, D to F and H).

## Discussion

In this study, we demonstrate that growth signaling and CD59-mediated immune-evasion signaling, long treated as independent cellular programs, can be physically coupled at the PM by a dedicated nanoscale structure. Using super-resolution single-molecule imaging, we identify GEM (Growth and Evasion Metastable hub) as a previously unrecognized, nanoscale, (≈34 nm), metastable (8.5-s lifetime), liquid-like signaling platform on the PM that physically integrates RTK and CD59 inputs within the same nanoscale assembly and supports cooperative, supralinear amplification of survival signaling, including PLCγ1/PI3K recruitment and Ca²⁺/Akt signaling. GEM assembles constitutively at endogenous protein levels through LLPS-like interactions driven primarily by zyxin’s IDR2. In addition, GEM contributes to CD59-mediated immune evasion via CDC resistance.

A defining feature of GEM is its nanoscale liquidity. Unlike previously described LLPS-driven signaling condensates, which are typically micron-scale and often require overexpression or non-physiological concentrations *in vitro* (*56–59*). GEM forms constitutively at endogenous protein levels and remains confined to the tens-of-nanometer length scales. It is also distinct from stable micron-scale signaling structures observed in specialized contexts (e.g., neuronal and immune synapses (*71–73*), and septin-stabilized blebs (*74*)), and becomes apparent only through single-molecule imaging capable of detecting individual recruited molecules and GEM clusters composed of as few as 2–20 constituents, as observed in the present study. Importantly, GEM’s short lifetime (≈8.5 s) and the sub-second dwell lifetimes of recruited signaling molecules (0.1–0.4 s) ensure rapid and reliable signal termination (*48*).

These physical characteristics plausibly provide the basis for GEMs’ unique ability to cooperatively integrate the concurrent RTK and CD59 inputs. We propose that repeated encounters between FAK and SFKs by nanoconfinement within liquid-like GEM enhance reciprocal chain activation, a process we term a “kinase spiral” (Fig. 5F). This kinase spiral can, in turn, drive supralinear amplification of downstream PI3K-PI(3,4,5)P_3_-Akt and PLCγ-IP_3_-Ca^2+^ outputs, pathways central to cell survival and proliferation.

GEM also enhances CD59’s two key functions: immune evasion via CDC resistance by co-recruiting CD59 and MAC precursors (Figs. 1, J and K, and 4, K to O) and CD59-dependent cell-growth signaling (Fig. 4P). Collectively, these results provide a mechanistic basis for the coupling of growth signaling and CDC resistance signaling, previously considered separate processes, at the PM that promotes cell survival and tumor growth *in vivo*.

More generally, confining signaling components within a liquid nanoscale volume can dramatically increase encounter frequencies, enabling cooperative, supralinear signal amplification beyond what is achievable by classical solid scaffolds (*75*) or diffusion-limited cytoplasmic signaling (*48*). A related precedent is FcεRI phosphorylation by Lyn in raft nanodomains induced by receptor oligomerization (*76*).

GEM formation is enhanced by raft domains, PI(4,5)P_2_, and cortical actin filaments (Fig. 2, D, G, and I to K, and fig. S4). Talin and RIAM can bind to PI(4,5)P_2_ associated with raft domains, and might be activated by lipid-binding (*77*). Both RIAM and folded talin directly bind PI(4,5)P₂-rich membranes, allowing them to co-assemble at low-tension cortical sites before any force-driven talin unfolding or adhesion maturation (*78–81*). In this pre-mechanochemical regime, integrins typically reside in ligand-engaged or preactivated, but not fully tension-activated, states that support receptor-proximal signaling without triggering vinculin recruitment or IAC growth (*82*, *83*). Our results suggest that GEM selectively captures this PI(4,5)P_2_**-**dependent, low-force configuration as a biochemical pre-assembly step for GEM formation.

Importantly, this arrangement might create a prewet membrane surface that favors zyxin/VASP recruitment and condensation (*35–38*), converting the RIAM–talin precomplex into the nanoscale liquid-like scaffold characteristic of GEM. In this way, GEM leverages only the tension-independent initiation module of the adhesion machinery and repurposes it to seed a zyxin/VASP-based nano-liquid hub dedicated to signal integration, rather than mechanical anchoring. The cortical actin meshwork may further help establish a prewet surface for recruiting talin and VASP for GEM assembly.

From an evolutionary perspective, phylogenetic analyses suggest that many IAC components originated as ancient signaling scaffolds before being co-opted for force transmission (*84*, *85*), although talin might also have played a central role in the evolution of cell adhesion and force transmission (*86*). GEM might thus preserve a primordial signaling function of these scaffolds that persists in modern cells alongside their mechanical roles.

LLPS drivers for IAC formation include p130Cas, FAK, paxillin, kindlin, vinculin, talin, and integrin tails (*55*, *77*, *87*, *88*), but zyxin’s IDRs are largely excluded from mature IACs, whereas the dominant GEM driver is zyxin’s IDR2 (Fig. 3H). These findings underscore that distinct molecular grammars underlie different condensate types within the same protein network. However, given the overlap in components between GEMs and IACs, certain IAC states might also function in signaling under specific mechanical and compositional conditions, whereas GEMs on the basal PM might convert into IACs (*89*) potentially under mechanical stress (*15*, *90*, *91*).

GEM also differs fundamentally from previously described aster structures or GPI-anchored protein nanoclusters, which are typically larger (>500-nm structures) and depend on integrin force-induced activation, vinculin engagement, and/or overexpression of GPI-anchored proteins (*15*, *92*, *93*). Nor does GEM conform to classical scaffold models composed of a small number of adaptor proteins or solid-like assemblies (*75*, *94*). Instead, GEM operates as a nano-liquid reactor, in which rapid molecular exchanges and high encounter frequencies dramatically accelerate reaction kinetics.

The physiological relevance of GEM-based signal integration is supported by our *in vivo* data. Consistent with the IDR2-dependent nano-liquid organization characterized *in silico* and in cultured cells, restoration of GEM through the zyxin IDR2 rescues tumor growth in multiple mouse models and human cancer cell lines (Fig. 6 and fig, S14). GEM is required for tumor growth even without CD59, whereas CD59 further enhances this effect via immune evasion (CDC resistance) and CD59-dependent growth signaling (Fig. 6, F to H, and fig, S14G). Importantly, zyxin-KO MEFs still formed tumors, albeit slowly (Fig. 6, B to D), indicating that GEM is not an exclusive driver of proliferation in a redundant pro-survival network. Together, these observations support that GEM acts as a quantitative amplifier that confers a substantial fitness advantage *in vivo*, where complement remains active.

Many GEM components, including CD59, integrin, talin, VASP, and zyxin, are overexpressed in diverse cancers (*95–103*), suggesting that the up-regulation of a nano-liquid signaling infrastructure might be a general strategy for tumor survival under immune pressure. With the growing recognition of LLPS in cancer biology (*104*, *105*), GEM highlights a previously unappreciated dimension: liquid condensates operating at the nanoscale, rather than micron-scale, can dominate signaling outcomes *in vivo*.

Several mechanistic issues remain. These include how receptors are recruited and transiently retained at GEMs, how integrin/talin activation states are configured, and how GEMs relate to adhesion structures on the basal membrane.

In summary, this work establishes GEM as a nano-liquid signaling hub that integrates the growth and CD59-mediated immune-evasion pathway through confinement-enhanced, cooperative kinase activation. By extending LLPS concepts from large micron-scale condensates to metastable nanoscale hubs, our findings reveal a previously unappreciated layer of cellular organization that governs cell survival signaling *in vivo*. Similar structures may operate across many signaling systems, and uncovering them will require single-molecule approaches capable of resolving dynamics at the molecular scale.

## Acknowledgments

We thank the researchers who kindly provided us with the cell lines and cDNAs used in this research (details are given in the materials and methods). We also thank Drs. P. Kanchanawong, S. Liu, and M. Kham of the National University of Singapore for technical help with the mouse embryonic stem cell culture, Drs. P. Barzaghi and B. Humbel of the OIST Imaging Section (IMG) for their help with FRAP experiments, Ms. F. Koja of the OIST Instrumental Analysis Section (IAS) for supporting FACS experiments, the OIST Sequencing Section (SQC) for their help in performing DNA sequencing, the OIST Animal Resource Section (ARS) for their help in animal care, and Dr. K. Simons of Lipotype GmbH, Dr. S. Mayor of National Centre for Biological Sciences, and all members of the Kusumi laboratory for valuable discussions. Illustrations in Fig. 6, A and I were created using BioRender.com.

## Funding

Japan Society for the Promotion of Science (JSPS) Grants-in-Aid for Scientific Research Kiban A 21H04772 (AK)

JSPS Grants-in-Aid for Scientific Research (S) 16H06386 (AK)

JSPS Grants-in-Aid for Scientific Research (B) 20H02585 (TKF)

JSPS Grants-in-Aid for Scientific Research (B) 24K01310 (TKF)

JSPS Grants-in-Aid for Scientific Research (C) 17K07333 (RSK)

JSPS Grant-in-Aid for Early-Career Scientists (B) 26870292 (RSK)

JSPS Grant-in-Aid for Early-Career Scientists 21K15058 (TAT)

JSPS Grant-in-Aid for Early-Career Scientists 25K18434 (TAT)

JSPS Grants-in-Aid for Scientific Research (B) 23K27326 (HI)

JSPS Grant-in-Aid for Challenging Research (Exploratory) 18K19001 (TKF)

JSPS Grant-in-Aid for Innovative Areas 18H04671 (KGNS)

JSPS Grants-in-Aid for Transformative Research Areas (A) 21H05252 (TKF)

JSPS Grants-in-Aid for Transformative Research Areas (A) 23H04414 (RSK)

Japan Science and Technology Agency (JST) grant in the program of the Core Research for Evolutional Science and Technology (CREST) in the field of “Biodynamics” JPMJCR14W2 (AK)

JST grant in the program of CREST in the field of “Extracellular Fine Particles” JPMJCR18H2 (KGNS)

JST grant in the program of CREST in the field of “Cell Control” JPMJCR24B3 (KGNS) JST grant ACT-X JPMJAX211B (TAT)

WPI-iCeMS of Kyoto University is supported by the World Premiere Research Center Initiative (WPI) of MEXT.

Start-up funds from German Center for Neurodegenerative Diseases (DZNE) (DM)

German Research Foundation SFB 1286/B10 (DM)

German Research Foundation MI 2104 (DM)

European Research Council Grant 101078172 (DM)

Human Frontier Science Organization RGEC32/2023 (DM)

Fellowship of the Innovative Minds Program of the German Dementia Association (CH)

## Author contributions

T.A.T. performed large majorities of the single-fluorescent-molecule imaging, biochemical, and molecular biology experiments and analyses. T.A.T., T.K.F., and A.K. developed the single-molecule imaging camera system, set up single-molecule imaging stations, and developed the analysis software. T.A.T., K.G.N.S., and A.K. established single-molecule observation conditions for CD59 and PDGFR signaling cascades. C.H. and D.M. purified zyxin and VASP and, together with T.A.T., performed LLPS and FRAP experiments. B.T. generated protein probes for cholesterol and sphingomyelin. T.A.T., D.S., and H.I. designed and performed the animal experiments and analyzed the data. C.T. and Y.S. performed coarse-grained molecular dynamics simulations. T.A.T., C.H., D.S., Y.L.N., K.M.H., R.S.K., T.K.F., K.G.N.S., H.I., D.M., and A.K. participated in discussions throughout the course of this research. T.A.T. and A.K. conceived and formulated this project and evaluated and discussed the data. T.A.T. and A.K. wrote the manuscript and all authors participated in its revision.

## Data, code, and materials availability

All data are available in the main text or the supplementary materials. All cell lines, plasmids, and materials used in this study are available from the corresponding authors under a materials transfer agreement (MTA) with OIST, Japan.

## References and Notes

1. M. A. Lemmon, J. Schlessinger, Cell signaling by receptor tyrosine kinases. Cell 141, 1117–1134 (2010).

2. C. Tomuleasa, A.-B. Tigu, R. Munteanu, C.-S. Moldovan, D. Kegyes, A. Onaciu, D. Gulei, G. Ghiaur, H. Einsele, C. M. Croce, Therapeutic advances of targeting receptor tyrosine kinases in cancer. Signal Transduct. Target. Ther. 9, 201 (2024).

3. E. C. Couves, S. Gardner, T. B. Voisin, J. K. Bickel, P. J. Stansfeld, E. W. Tate, D. Bubeck, Structural basis for membrane attack complex inhibition by CD59. Nat. Commun. 14, 890 (2023).

4. K. G. Suzuki, T. K. Fujiwara, F. Sanematsu, R. Iino, M. Edidin, A. Kusumi, GPI-anchored receptor clusters transiently recruit Lyn and G alpha for temporary cluster immobilization and Lyn activation: single-molecule tracking study 1. J. Cell Biol. 177, 717–30 (2007).

5. K. G. Suzuki, T. K. Fujiwara, M. Edidin, A. Kusumi, Dynamic recruitment of phospholipase C gamma at transiently immobilized GPI-anchored receptor clusters induces IP3-Ca2+ signaling: single-molecule tracking study 2. J. Cell Biol. 177, 731–42 (2007).

6. I. Koyama-Honda, T. K. Fujiwara, R. S. Kasai, K. G. N. Suzuki, E. Kajikawa, H. Tsuboi, T. A. Tsunoyama, A. Kusumi, High-speed single-molecule imaging reveals signal transduction by induced transbilayer raft phases. J. Cell Biol. 219, e202006125 (2020).

7. J. Humeau, J. M. B.-S. Pedro, I. Vitale, L. Nuñez, C. Villalobos, G. Kroemer, L. Senovilla, Calcium signaling and cell cycle: Progression or death. Cell Calcium 70, 3–15 (2018).

8. W. Guo, F. G. Giancotti, Integrin signalling during tumour progression. Nat. Rev. Mol. Cell Biol. 5, 816–826 (2004).

9. Z. Chen, D. Oh, A. K. Dubey, M. Yao, B. Yang, J. T. Groves, M. Sheetz, EGFR family and Src family kinase interactions: mechanics matters? Curr. Opin. Cell Biol. 51, 97–102 (2018).

10. M. A. Alfonzo-Méndez, K. A. Sochacki, M.-P. Strub, J. W. Taraska, Dual clathrin and integrin signaling systems regulate growth factor receptor activation. Nat. Commun. 13, 905 (2022).

11. T. S. van Zanten, A. Cambi, M. Koopman, B. Joosten, C. G. Figdor, M. F. Garcia-Parajo, Hotspots of GPI-anchored proteins and integrin nanoclusters function as nucleation sites for cell adhesion. Proc. Natl. Acad. Sci. USA 106, 18557–18562 (2009).

12. V. F. Fiore, P. W. Strane, A. V. Bryksin, E. S. White, J. S. Hagood, T. H. Barker, Conformational coupling of integrin and Thy-1 regulates Fyn priming and fibroblast mechanotransduction. J. Cell Biol. 211, 173–190 (2015).

13. X. Cui, X. Zhang, H. Bu, N. Liu, H. Li, X. Guan, H. Yan, Y. Wang, H. Zhang, Y. Ding, M. Cheng, Shear stress-mediated changes in the expression of complement regulatory protein CD59 on human endothelial progenitor cells by ECM-integrinαVβ3-F-actin pathway in vitro. Biochem. Biophys. Res. Commun. 494, 416–421 (2017).

14. M. G. Strainic, J. Liu, F. An, E. Bailey, A. Esposito, J. Hamann, P. S. Heeger, M. E. Medof, CD55 is essential for CD103^+^ dendritic cell tolerogenic responses that protect against autoimmunity. Am. J. Pathol. 189, 1386–1401 (2019).

15. J. M. Kalappurakkal, A. A. Anilkumar, C. Patra, T. S. van Zanten, M. P. Sheetz, S. Mayor, Integrin mechano-chemical signaling generates plasma membrane nanodomains that promote cell spreading. Cell 177, 1738–1756. e23 (2019).

16. M. Z. Miao, J. S. Lee, K. M. Yamada, R. F. Loeser, Integrin signalling in joint development, homeostasis and osteoarthritis. Nat. Rev. Rheumatol. 20, 492–509 (2024).

17. R. Nishimura, P. Kanchanawong, Nanoscale mechano-adaption of integrin-based cell adhesions: New tools and techniques lead the way. Curr. Opin. Cell Biol. 94, 102509 (2025).

18. T. Chen, G. Giannone, Single molecule imaging unveils cellular architecture, dynamics and mechanobiology. Curr. Opin. Cell Biol. 88, 102369 (2024).

19. M. Theodosiou, M. Widmaier, R. T. Böttcher, E. Rognoni, M. Veelders, M. Bharadwaj, A. Lambacher, K. Austen, D. J. Müller, R. Zent, R. Fässler, Kindlin-2 cooperates with talin to activate integrins and induces cell spreading by directly binding paxillin. eLife 5, e10130 (2016).

20. P. Zhou, T. A. Tsunoyama, R. S. Kasai, K. M. Hirosawa, Z. Kalay, A. Aladag, T. K. Fujiwara, T. Yokoyama, M. Sakamoto, R. Kise, M. Yanagawa, A. Inoue, S. Pigolotti, A. Kusumi, Single-molecule methods for characterizing receptor dimers reveal metastable opioid receptor homodimers that induce functional modulation. Nat. Commun. 16, 9858 (2025).

21. M. B. Stone, S. L. Veatch, Steady-state cross-correlations for live two-colour super-resolution localization data sets. Nat. Commun. 6, 7347 (2014).

22. L. M. Hoffman, C. C. Jensen, S. Kloeker, C.-L. A. Wang, M. Yoshigi, M. C. Beckerle, Genetic ablation of zyxin causes Mena/VASP mislocalization, increased motility, and deficits in actin remodeling. J. Cell Biol. 172, 771–782 (2006).

23. T. K. Fujiwara, K. Iwasawa, Z. Kalay, T. A. Tsunoyama, Y. Watanabe, Y. M. Umemura, H. Murakoshi, K. G. N. Suzuki, Y. L. Nemoto, N. Morone, A. Kusumi, Confined diffusion of transmembrane proteins and lipids induced by the same actin meshwork lining the plasma membrane. Mol. Biol. Cell 27, 1101–1119 (2016).

24. H. Winkelmann, C. P. Richter, J. Eising, J. Piehler, R. Kurre, Correlative single-molecule and structured illumination microscopy of fast dynamics at the plasma membrane. Nat. Commun. 15, 5813 (2024).

25. P. Lajoie, J. G. Goetz, J. W. Dennis, I. R. Nabi, Lattices, rafts, and scaffolds: domain regulation of receptor signaling at the plasma membrane. J. Cell Biol. 185, 381–385 (2009).

26. K. Simons, J. L. Sampaio, Membrane organization and lipid rafts. Cold Spring Harb. Perspect. Biol. 3, a004697 (2011).

27. A. Kusumi, T. K. Fujiwara, T. A. Tsunoyama, R. S. Kasai, A. A. Liu, K. M. Hirosawa, M. Kinoshita, N. Matsumori, N. Komura, H. Ando, K. G. N. Suzuki, Defining raft domains in the plasma membrane. Traffic 21, 106–137 (2020).

28. M. Maekawa, Domain 4 (D4) of perfringolysin O to visualize cholesterol in cellular membranes—the update. Sensors 17, 504 (2017).

29. S. Endapally, D. Frias, M. Grzemska, A. Gay, D. R. Tomchick, A. Radhakrishnan, Molecular discrimination between two conformations of sphingomyelin in plasma membranes. Cell 176, 1040–1053. e17 (2019).

30. Y. Uchida, J. Hasegawa, D. Chinnapen, T. Inoue, S. Okazaki, R. Kato, S. Wakatsuki, R. Misaki, M. Koike, Y. Uchiyama, S. Iemura, T. Natsume, R. Kuwahara, T. Nakagawa, K. Nishikawa, K. Mukai, E. Miyoshi, N. Taniguchi, D. Sheff, W. I. Lencer, T. Taguchi, H. Arai, Intracellular phosphatidylserine is essential for retrograde membrane traffic through endosomes. Proc. Natl. Acad. Sci. USA 108, 15846–15851 (2011).

31. S. Lee, Y. Uchida, J. Wang, T. Matsudaira, T. Nakagawa, T. Kishimoto, K. Mukai, T. Inaba, T. Kobayashi, R. S. Molday, T. Taguchi, H. Arai, Transport through recycling endosomes requires EHD1 recruitment by a phosphatidylserine translocase. EMBO J. 34, 669–688 (2015).

32. M. Furutani, K. Tsujita, T. Itoh, T. Ijuin, T. Takenawa, Application of phosphoinositide-binding domains for the detection and quantification of specific phosphoinositides. Anal. Biochem. 355, 8–18 (2006).

33. O. Idevall-Hagren, E. J. Dickson, B. Hille, D. K. Toomre, P. D. Camilli, Optogenetic control of phosphoinositide metabolism. Proc. Natl. Acad. Sci. USA 109, E2316–E2323 (2012).

34. M. de S. Santos, R. M. Z. G. Naal, B. Baird, D. Holowka, Inhibitors of PI(4,5)P₂ synthesis reveal dynamic regulation of IgE receptor signaling by phosphoinositides in RBL mast cells. Mol. Pharmacol. 83, 793–804 (2013).

35. M. Rouches, S. L. Veatch, B. B. Machta, Surface densities prewet a near-critical membrane. Proc. Natl. Acad. Sci. USA 118, e2103401118 (2021).

36. T. A. Tsunoyama, C. Hoffmann, B. Tang, K. M. Hirosawa, Y. L. Nemoto, R. S. Kasai, T. K. Fujiwara, K. G. N. Suzuki, D. Milovanovic, A. Kusumi, iTRVZ: liquid nano-platform for signal integration on the plasma membrane. bioRxiv, 2021.12.30.474523 (2022).

37. C. Hoffmann, D. Milovanovic, Dipping contacts – a novel type of contact site at the interface between membraneless organelles and membranes. J. Cell Sci. 136, jcs261413 (2023).

38. Y. Wan, R. Hudson, J. Smith, J. D. Forman-Kay, J. A. Ditlev, Protein interactions, calcium, phosphorylation, and cholesterol modulate CFTR cluster formation on membranes. Proc. Natl. Acad. Sci. USA 122, e2424470122 (2025).

39. P. Sengupta, S. Van Engelenburg, J. Lippincott-Schwartz, Visualizing cell structure and function with point-localization superresolution imaging. Dev. Cell 23, 1092–1102 (2012).

40. F. Levet, E. Hosy, A. Kechkar, C. Butler, A. Beghin, D. Choquet, J.-B. Sibarita, SR-Tesseler: a method to segment and quantify localization-based super-resolution microscopy data. Nat. Methods 12, 1065–1071 (2015).

41. I. M. Khater, I. R. Nabi, G. Hamarneh, A review of super-resolution single-molecule localization microscopy cluster analysis and quantification methods. Patterns 1, 100038 (2020).

42. D. Piovesan, M. Necci, N. Escobedo, A. M. Monzon, A. Hatos, I. Mičetić, F. Quaglia, L. Paladin, P. Ramasamy, Z. Dosztányi, W. F. Vranken, N. E. Davey, G. Parisi, M. Fuxreiter, S. C. E. Tosatto, MobiDB: intrinsically disordered proteins in 2021. Nucleic Acids Res. 49, D361–D367 (2020).

43. K. Jaqaman, J. A. Ditlev, Biomolecular condensates in membrane receptor signaling. Curr. Opin. Cell Biol. 69, 48–54 (2021).

44. C. Hoffmann, R. Sansevrino, G. Morabito, C. Logan, R. M. Vabulas, A. Ulusoy, M. Ganzella, D. Milovanovic, Synapsin condensates recruit alpha-synuclein. J. Mol. Biol. 433, 166961 (2021).

45. S. Peng, W. Li, Y. Yao, W. Xing, P. Li, C. Chen, Phase separation at the nanoscale quantified by dcFCCS. Proc. Natl. Acad. Sci. USA 117, 27124–27131 (2020).

46. M. Kar, F. Dar, T. J. Welsh, L. T. Vogel, R. Kühnemuth, A. Majumdar, G. Krainer, T. M. Franzmann, S. Alberti, C. A. M. Seidel, T. P. J. Knowles, A. A. Hyman, R. V. Pappu, Phase-separating RNA-binding proteins form heterogeneous distributions of clusters in subsaturated solutions. Proc. Natl. Acad. Sci. USA 119, e2202222119 (2022).

47. C. Lan, J. Kim, S. Ulferts, F. Aprile-Garcia, S. Weyrauch, A. Anandamurugan, R. Grosse, R. Sawarkar, A. Reinhardt, T. Hugel, Quantitative real-time in-cell imaging reveals heterogeneous clusters of proteins prior to condensation. Nat. Commun. 14, 4831 (2023).

48. A. Kusumi, T. A. Tsunoyama, K. G. N. Suzuki, T. K. Fujiwara, A. Aladag, Transient, nano-scale, liquid-like molecular assemblies coming of age. Curr. Opin. Cell Biol. 89, 102394 (2024).

49. Y. Ge, T. Paul, M. Gordiychuk, N. Das, Y. Zhang, S. Myong, FUS nanoclusters are a distinct state within the dilute phase. Nat. Commun. 16, 9956 (2025).

50. M. K. Shinn, D. T. Tomares, V. Liu, A. Pant, Y. Qiu, A. Vitalis, Y. J. Song, Y. Ayala, K. M. Ruff, G. W. Strout, M. D. Lew, K. V. Prasanth, R. V. Pappu, Nuclear speckle proteins form intrinsic and MALAT1-dependent microphases. Cell 189, 832–852.e24 (2026).

51. T. Wiegand, J. Liu, L. Vogeley, I. LuValle-Burke, J. Geisler, A. W. Fritsch, A. A. Hyman, S. W. Grill, Actin polymerization counteracts prewetting of N-WASP on supported lipid bilayers. Proc. Natl. Acad. Sci. USA 121, e2407497121 (2024).

52. J. A. Joseph, A. Reinhardt, A. Aguirre, P. Y. Chew, K. O. Russell, J. R. Espinosa, A. Garaizar, R. Collepardo-Guevara, Physics-driven coarse-grained model for biomolecular phase separation with near-quantitative accuracy. Nat. Comput. Sci. 1, 732–743 (2021).

53. K. Graham, A. Chandrasekaran, L. Wang, A. Ladak, E. M. Lafer, P. Rangamani, J. C. Stachowiak, Liquid-like VASP condensates drive actin polymerization and dynamic bundling. Nat. Phys. 19, 574–585 (2023).

54. K. B. Narayan, H. P. James, J. Cope, S. Mondal, L. Baeyens, F. Milano, J. Zheng, M. Krause, T. Baumgart, VASP phase separation with priming proteins of fast endophilin mediated endocytosis modulates actin polymerization. bioRxiv, 2024.03.21.586200 (2024).

55. C.-P. Hsu, J. Aretz, A. Hordeichyk, R. Fässler, A. R. Bausch, Surface-induced phase separation of reconstituted nascent integrin clusters on lipid membranes. Proc. Natl. Acad. Sci. USA 120, e2301881120 (2023).

56. X. Su, J. A. Ditlev, E. Hui, W. Xing, S. Banjade, J. Okrut, D. S. King, J. Taunton, M. K. Rosen, R. D. Vale, Phase separation of signaling molecules promotes T cell receptor signal transduction. Science 352, 595–599 (2016).

57. L. B. Case, X. Zhang, J. A. Ditlev, M. K. Rosen, Stoichiometry controls activity of phase-separated clusters of actin signaling proteins. Science 363, 1093–1097 (2019).

58. W. Y. C. Huang, S. Alvarez, Y. Kondo, Y. K. Lee, J. K. Chung, H. Y. M. Lam, K. H. Biswas, J. Kuriyan, J. T. Groves, A molecular assembly phase transition and kinetic proofreading modulate Ras activation by SOS. Science 363, 1098–1103 (2019).

59. E. Dine, E. H. Reed, J. E. Toettcher, Positive feedback between the T cell kinase Zap70 and its substrate LAT acts as a clustering-dependent signaling switch. Cell Reports 35, 109280–109280 (2021).

60. S. H. Ryu, K. S. Cho, K. Y. Lee, P. G. Suh, S. G. Rhee, Purification and characterization of two immunologically distinct phosphoinositide-specific phospholipases C from bovine brain. J. Biol. Chem. 262, 12511–8 (1987).

61. Y. S. Bae, L. G. Cantley, C. S. Chen, S. R. Kim, K. S. Kwon, S. G. Rhee, Activation of phospholipase C-gamma by phosphatidylinositol 3,4,5-trisphosphate. J. Biol. Chem. 273, 4465–9 (1998).

62. A. Glaviano, A. S. C. Foo, H. Y. Lam, K. C. H. Yap, W. Jacot, R. H. Jones, H. Eng, M. G. Nair, P. Makvandi, B. Geoerger, M. H. Kulke, R. D. Baird, J. S. Prabhu, D. Carbone, C. Pecoraro, D. B. L. Teh, G. Sethi, V. Cavalieri, K. H. Lin, N. R. Javidi-Sharifi, E. Toska, M. S. Davids, J. R. Brown, P. Diana, J. Stebbing, D. A. Fruman, A. P. Kumar, PI3K/AKT/mTOR signaling transduction pathway and targeted therapies in cancer. Mol. Cancer 22, 138 (2023).

63. C.-C. Lin, K. M. Suen, P.-A. Jeffrey, L. Wieteska, J. A. Lidster, P. Bao, A. P. Curd, A. Stainthorp, C. Seiler, H. Koss, E. Miska, Z. Ahmed, S. D. Evans, C. Molina-París, J. E. Ladbury, Receptor tyrosine kinases regulate signal transduction through a liquid-liquid phase separated state. Mol. Cell 82, 1089–1106. e12 (2022).

64. I. Acebrón, R. D. Righetto, C. Schoenherr, S. Buhr, P. Redondo, J. Culley, C. F. Rodríguez, C. Daday, N. Biyani, O. Llorca, A. Byron, M. Chami, F. Gräter, J. Boskovic, M. C. Frame, H. Stahlberg, D. Lietha, Structural basis of focal adhesion kinase activation on lipid membranes. EMBO J. 39, e104743 (2020).

65. H. Wei, M. Malik, A. M. Sheikh, G. Merz, W. T. Brown, X. Li, Abnormal cell properties and down-regulated FAK-Src complex signaling in B lymphoblasts of autistic subjects. Am. J. Pathol. 179, 66–74 (2011).

66. Q. Wan, T. TruongVo, H. E. Steele, A. Ozcelikkale, B. Han, Y. Wang, J. Oh, H. Yokota, S. Na, Subcellular domain-dependent molecular hierarchy of SFK and FAK in mechanotransduction and cytokine signaling. Sci. Rep. 7, 9033 (2017).

67. B. Liu, O. J. Stone, M. Pablo, J. C. Herron, A. T. Nogueira, O. Dagliyan, J. B. Grimm, L. D. Lavis, T. C. Elston, K. M. Hahn, Biosensors based on peptide exposure show single molecule conformations in live cells. Cell 184, 5670–5685.e23 (2021).

68. Y. Shin, J. Berry, N. Pannucci, M. P. Haataja, J. E. Toettcher, C. P. Brangwynne, Spatiotemporal control of intracellular phase transitions using light-activated optoDroplets. Cell 168, 159–171.e14 (2017).

69. E. Dine, A. A. Gil, G. Uribe, C. P. Brangwynne, J. E. Toettcher, Protein phase separation provides long-term memory of transient spatial stimuli. Cell Syst. 6, 655–663.e5 (2018).

70. D. K. Hirenallur-Shanthappa, J. A. Ramírez, B. M. Iritani, Patient derived tumor xenograft models. Academic Press, 57–73 (2017).

71. M. L. Dustin, D. Depoil, New insights into the T cell synapse from single molecule techniques. Nat. Rev. Immunol. 11, 672–684 (2011).

72. T. C. Südhof, The cell biology of synapse formation. J. Cell Biol. 220, e202103052 (2021).

73. B. Jia, Z. Shen, S. Zhu, J. Huang, Z. Liao, S. Zhao, H. Li, S. Chen, Y. Xu, Y. Wang, H. Peng, G. Bai, Y. Lu, P. Tong, W. Tao, M. Zhang, Shank3 oligomerization governs material properties of the postsynaptic density condensate and synaptic plasticity. Cell 188, 6473–6491.e21 (2025).

74. A. D. Weems, E. S. Welf, M. K. Driscoll, F. Y. Zhou, H. Mazloom-Farsibaf, B.-J. Chang, V. S. Murali, G. M. Gihana, B. G. Weiss, J. Chi, D. Rajendran, K. M. Dean, R. Fiolka, G. Danuser, Blebs promote cell survival by assembling oncogenic signalling hubs. Nature 615, 517–525 (2023).

75. C. J. DiRusso, M. Dashtiahangar, T. D. Gilmore, Scaffold proteins as dynamic integrators of biological processes. J. Biol. Chem. 298, 102628 (2022).

76. N. Bag, A. Wagenknecht-Wiesner, A. Lee, S. M. Shi, D. A. Holowka, B. A. Baird, Lipid-based and protein-based interactions synergize transmembrane signaling stimulated by antigen clustering of IgE receptors. Proc. Natl. Acad. Sci. USA 118, e2026583118 (2021).

77. T. Litschel, C. F. Kelley, X. Cheng, L. Babl, N. Mizuno, L. B. Case, P. Schwille, Membrane-induced 2D phase separation of the focal adhesion protein talin. Nat. Commun. 15, 4986 (2024).

78. E. M. Lafuente, A. A. F. L. van Puijenbroek, M. Krause, C. V. Carman, G. J. Freeman, A. Berezovskaya, E. Constantine, T. A. Springer, F. B. Gertler, V. A. Boussiotis, RIAM, an Ena/VASP and profilin ligand, interacts with Rap1-GTP and mediates Rap1-induced adhesion. Dev. Cell 7, 585–595 (2004).

79. J. P. Wynne, J. Wu, W. Su, A. Mor, N. Patsoukis, V. A. Boussiotis, S. R. Hubbard, M. R. Philips, Rap1-interacting adapter molecule (RIAM) associates with the plasma membrane via a proximity detector. J. Cell Biol. 199, 317–329 (2012).

80. B. T. Goult, X.-P. Xu, A. R. Gingras, M. Swift, B. Patel, N. Bate, P. M. Kopp, I. L. Barsukov, D. R. Critchley, N. Volkmann, D. Hanein, Structural studies on full-length talin1 reveal a compact auto-inhibited dimer: implications for talin activation. J. Struct. Biol. 184, 21–32 (2013).

81. T. Orré, A. Joly, Z. Karatas, B. Kastberger, C. Cabriel, R. T. Böttcher, S. Lévêque-Fort, J.-B. Sibarita, R. Fässler, B. Wehrle-Haller, O. Rossier, G. Giannone, Molecular motion and tridimensional nanoscale localization of kindlin control integrin activation in focal adhesions. Nat. Commun. 12, 3104 (2021).

82. E. M. Morse, N. N. Brahme, D. A. Calderwood, Integrin cytoplasmic tail interactions. Biochemistry 53, 810–820 (2014).

83. J. Li, M. H. Jo, J. Yan, T. Hall, J. Lee, U. López-Sánchez, S. Yan, T. Ha, T. A. Springer, Ligand binding initiates single-molecule integrin conformational activation. Cell 187, 2990–3005.e17 (2024).

84. A. Sebé-Pedrós, A. J. Roger, F. B. Lang, N. King, I. Ruiz-Trillo, Ancient origin of the integrin-mediated adhesion and signaling machinery. Proc. Natl. Acad. Sci. USA 107, 10142–10147 (2010).

85. I. Ruiz-Trillo, K. Kin, E. Casacuberta, The origin of metazoan multicellularity: a potential microbial black swan event. Annu. Rev. Microbiol. 77, 499–516 (2023).

86. S. Rangarajan, L. Espeter, H. C. A. Drexler, A. Chrostek-Grashoff, C. Grashoff, Talin force coupling underlies eukaryotic cell-substrate adhesion. Nat. Commun. 16, 10950 (2025).

87. L. B. Case, M. D. Pasquale, L. Henry, M. K. Rosen, Synergistic phase separation of two pathways promotes integrin clustering and nascent adhesion formation. eLife 11, e72588 (2022).

88. A. Kumar, K. Tanaka, M. A. Schwartz, Focal adhesion-derived liquid-liquid phase separations regulate mRNA translation. eLife, doi: 10.7554/elife.96157.1 (2024).

89. J. Aretz, M. Aziz, N. Strohmeyer, M. Sattler, R. Fässler, Talin and kindlin use integrin tail allostery and direct binding to activate integrins. Nat. Struct. Mol. Biol. 30, 1913–1924 (2023).

90. V. Swaminathan, G. M. Alushin, C. M. Waterman, Mechanosensation: a catch bond that only hooks one way. Curr. Biol. 27, R1158–R1160 (2017).

91. S. Massou, F. N. Vicente, F. Wetzel, A. Mehidi, D. Strehle, C. Leduc, R. Voituriez, O. Rossier, P. Nassoy, G. Giannone, Cell stretching is amplified by active actin remodelling to deform and recruit proteins in mechanosensitive structures. Nat. Cell Biol. 22, 1011–1023 (2020).

92. K. Gowrishankar, S. Ghosh, S. Saha, C. Rumamol, S. Mayor, M. Rao, Active remodeling of cortical actin regulates spatiotemporal organization of cell surface molecules. Cell 149, 1353–1367 (2012).

93. M. F. Garcia-Parajo, S. Mayor, The ubiquitous nanocluster: a molecular scale organizing principle that governs cellular information flow. Curr. Opin. Cell Biol. 86, 102285 (2024).

94. C. T. Pawson, J. D. Scott, Signal integration through blending, bolstering and bifurcating of intracellular information. Nat. Struct. Mol. Biol. 17, 653–658 (2010).

95. A. Chaturvedi, L. M. Hoffman, C. C. Jensen, Y.-C. Lin, A. H. Grossmann, R. L. Randall, S. L. Lessnick, A. L. Welm, M. C. Beckerle, Molecular dissection of the mechanism by which EWS/FLI expression compromises actin cytoskeletal integrity and cell adhesion in Ewing sarcoma. Mol. Biol. Cell 25, 2695–2709 (2014).

96. B. Ma, H. Cheng, R. Gao, C. Mu, L. Chen, S. Wu, Q. Chen, Y. Zhu, Zyxin-Siah2–Lats2 axis mediates cooperation between Hippo and TGF-β signalling pathways. Nat. Commun. 7, 11123 (2016).

97. A. Kotb, M. E. Hyndman, T. R. Patel, The role of zyxin in regulation of malignancies. Heliyon 4, e00695 (2018).

98. H. Hamidi, J. Ivaska, Every step of the way: integrins in cancer progression and metastasis. Nat. Rev. Cancer 18, 533–548 (2018).

99. J. Zhou, Y. Zeng, L. Cui, X. Chen, S. Stauffer, Z. Wang, F. Yu, S. M. Lele, G. A. Talmon, A. R. Black, Y. Chen, J. Dong, Zyxin promotes colon cancer tumorigenesis in a mitotic phosphorylation-dependent manner and through CDK8-mediated YAP activation. Proc. Natl. Acad. Sci. USA 115, 201800621 (2018).

100. X. Xiang, Y. Wang, H. Zhang, J. Piao, S. Muthusamy, L. Wang, Y. Deng, W. Zhang, R. Kuang, D. D. Billadeau, S. Huang, J. Lai, R. Urrutia, N. Kang, Vasodilator-stimulated phosphoprotein promotes liver metastasis of gastrointestinal cancer by activating a β1-integrin-FAK-YAP1/TAZ signaling pathway. npj Precis. Oncol. 2, 2 (2018).

101. Y. Tian, L. Xu, Y. He, X. Xu, K. Li, Y. Ma, Y. Gao, D. Wei, L. Wei, Knockdown of RAC1 and VASP gene expression inhibits breast cancer cell migration. Oncol. Lett. 16, 2151–2160 (2018).

102. Y. Rezaie, F. Fattahi, B. Mashinchi, K. K. Hesari, S. Montazeri, E. Kalantari, Z. Madjd, L. S. Zanjani, High expression of Talin-1 is associated with tumor progression and recurrence in melanoma skin cancer patients. BMC Cancer 23, 302 (2023).

103. H. Shadman, S. Gomrok, C. Litle, Q. Cheng, Y. Jiang, X. Huang, J. D. Ziebarth, Y. Wang, A machine learning-based investigation of integrin expression patterns in cancer and metastasis. Sci. Rep. 15, 5270 (2025).

104. D. Cai, Z. Liu, J. Lippincott-Schwartz, Biomolecular condensates and their links to cancer progression. Trends Biochem. Sci. 46, 535–549 (2021).

105. T. P. López-Palacios, J. L. Andersen, Kinase regulation by liquid–liquid phase separation. Trends Cell Biol. 33, 649–666 (2023).

106. E. Rossi, B. Casali, G. Regolisti, S. Davoli, F. Perazzoli, A. Negro, C. Sani, B. Tumiati, D. Nicoli, Increased plasma levels of platelet-derived growth factor (PDGF-BB + PDGF-AB) in patients with never-treated mild essential hypertension. Am. J. Hypertens. 11, 1239–1243 (1998).

107. K. Murase, T. Fujiwara, Y. Umemura, K. Suzuki, R. Iino, H. Yamashita, M. Saito, H. Murakoshi, K. Ritchie, A. Kusumi, Ultrafine membrane compartments for molecular diffusion as revealed by single molecule techniques. Biophys. J. 86, 4075–4093 (2004).

108. N. Morone, T. Fujiwara, K. Murase, R. S. Kasai, H. Ike, S. Yuasa, J. Usukura, A. Kusumi, Three-dimensional reconstruction of the membrane skeleton at the plasma membrane interface by electron tomography. J. Cell Biol. 174, 851–862 (2006).

109. K. Naruse, X. Sai, N. Yokoyama, M. Sokabe, Uni-axial cyclic stretch induces c-src activation and translocation in human endothelial cells via SA channel activation. FEBS Lett. 441, 111–115 (1998).

110. M. Ihara, H. Tomimoto, H. Kitayama, Y. Morioka, I. Akiguchi, H. Shibasaki, M. Noda, M. Kinoshita, Association of the cytoskeletal GTP-binding protein Sept4/H5 with cytoplasmic inclusions found in Parkinson’s disease and other synucleinopathies. J. Biol. Chem. 278, 24095–24102 (2003).

111. M. Hooper, K. Hardy, A. Handyside, S. Hunter, M. Monk, HPRT-deficient (Lesch–Nyhan) mouse embryos derived from germline colonization by cultured cells. Nature 326, 292–295 (1987).

112. S. Xia, Y. B. Lim, Z. Zhang, Y. Wang, S. Zhang, C. T. Lim, E. K. F. Yim, P. Kanchanawong, Nanoscale architecture of the cortical actin cytoskeleton in embryonic stem cells. Cell Rep. 28, 1251–1267.e7 (2019).

113. S. A. McKinney, C. S. Murphy, K. L. Hazelwood, M. W. Davidson, L. L. Looger, A bright and photostable photoconvertible fluorescent protein. Nat. Methods 6, 131–133 (2009).

114. T. K. Fujiwara, S. Takeuchi, Z. Kalay, Y. Nagai, T. A. Tsunoyama, T. Kalkbrenner, K. Iwasawa, K. P. Ritchie, K. G. N. Suzuki, A. Kusumi, Development of ultrafast camera-based single fluorescent-molecule imaging for cell biology. J. Cell Biol. 222, e202110160 (2023).

115. T. K. Fujiwara, T. A. Tsunoyama, S. Takeuchi, Z. Kalay, Y. Nagai, T. Kalkbrenner, Y. L. Nemoto, L. H. Chen, A. C. E. Shibata, K. Iwasawa, K. P. Ritchie, K. G. N. Suzuki, A. Kusumi, Ultrafast single-molecule imaging reveals focal adhesion nano-architecture and molecular dynamics. J. Cell Biol. 222, e202110162 (2023).

116. D. S. Bindels, L. Haarbosch, L. van Weeren, M. Postma, K. E. Wiese, M. Mastop, S. Aumonier, G. Gotthard, A. Royant, M. A. Hink, T. W. J. Gadella, mScarlet: a bright monomeric red fluorescent protein for cellular imaging. Nat. Methods 14, 53–56 (2017).

117. M. Maekawa, G. D. Fairn, Complementary probes reveal that phosphatidylserine is required for the proper transbilayer distribution of cholesterol. J. Cell Sci. 128, 1422–1433 (2015).

118. B. B. Johnson, P. C. Moe, D. Wang, K. Rossi, B. L. Trigatti, A. P. Heuck, Modifications in perfringolysin O domain 4 alter the cholesterol concentration threshold required for binding. Biochemistry 51, 3373–3382 (2012).

119. T. A. Tsunoyama, Y. Watanabe, J. Goto, K. Naito, R. S. Kasai, K. G. N. Suzuki, T. K. Fujiwara, A. Kusumi, Super-long single-molecule tracking reveals dynamic-anchorage-induced integrin function. Nat. Chem. Biol. 14, 497–506 (2018).

120. Y. L. Nemoto, R. J. Morris, H. Hijikata, T. A. Tsunoyama, A. C. E. Shibata, R. S. Kasai, A. Kusumi, T. K. Fujiwara, Dynamic Meso-Scale Anchorage of GPI-Anchored Receptors in the Plasma Membrane: Prion Protein vs. Thy1. Cell Biochem Biophys 75, 399–412 (2017).

121. D. Ilić, E. A. C. Almeida, D. D. Schlaepfer, P. Dazin, S. Aizawa, C. H. Damsky, Extracellular matrix survival signals transduced by focal adhesion kinase suppress p53-mediated apoptosis. J. Cell Biol. 143, 547–560 (1998).

122. H.-S. Lee, P. Anekal, C. J. Lim, C.-C. Liu, M. H. Ginsberg, Two modes of integrin activation form a binary molecular switch in adhesion maturation. Mol. Biol. Cell 24, 1354–1362 (2013).

123. D. Tsuruta, M. Gonzales, S. B. Hopkinson, C. Otey, S. Khuon, R. D. Goldman, J. C. R. Jones, Microfilament-dependent movement of the β3 integrin subunit within focal contacts of endothelial cells. FASEB J. 16, 866–868 (2002).

124. J. Okrut, S. Prakash, Q. Wu, M. J. S. Kelly, J. Taunton, Allosteric N-WASP activation by an inter-SH3 domain linker in Nck. Proc. Natl Acad. Sci. USA 112, E6436–E6445 (2015).

125. Y. Sun, N. Thapa, A. C. Hedman, R. A. Anderson, Phosphatidylinositol 4,5-bisphosphate: targeted production and signaling. Bioessays 35, 513–522 (2013).

126. L. E. Diamond, K. R. McCurry, E. R. Oldham, M. Tone, H. Waldmann, J. L. Platt, J. S. Logan, Human CD59 expressed in transgenic mouse hearts inhibits the activation of complement. Transpl. Immunol. 3, 305–312 (1995).

127. C. M. Johannessen, J. S. Boehm, S. Y. Kim, S. R. Thomas, L. Wardwell, L. A. Johnson, C. M. Emery, N. Stransky, A. P. Cogdill, J. Barretina, G. Caponigro, H. Hieronymus, R. R. Murray, K. Salehi-Ashtiani, D. E. Hill, M. Vidal, J. J. Zhao, X. Yang, O. Alkan, S. Kim, J. L. Harris, C. J. Wilson, V. E. Myer, P. M. Finan, D. E. Root, T. M. Roberts, T. Golub, K. T. Flaherty, R. Dummer, B. L. Weber, W. R. Sellers, R. Schlegel, J. A. Wargo, W. C. Hahn, L. A. Garraway, COT drives resistance to RAF inhibition through MAP kinase pathway reactivation. Nature 468, 968–972 (2010).

128. N. P. Jones, J. Peak, S. Brader, S. A. Eccles, M. Katan, PLCγ1 is essential for early events in integrin signalling required for cell motility. J. Cell Sci. 118, 2695–2706 (2005).

129. S. Ishida, T. Matsu-ura, K. Fukami, T. Michikawa, K. Mikoshiba, Phospholipase C-β1 and β4 contribute to non-genetic cell-to-cell variability in histamine-induced calcium signals in HeLa cells. PLoS One 9, e86410 (2014).

130. F. P. Lindberg, H. D. Gresham, E. Schwarz, E. J. Brown, Molecular cloning of integrin-associated protein: an immunoglobulin family member with multiple membrane-spanning domains implicated in alpha v beta 3-dependent ligand binding. J. Cell Biol. 123, 485–496 (1993).

131. K. G. Suzuki, R. S. Kasai, K. M. Hirosawa, Y. L. Nemoto, M. Ishibashi, Y. Miwa, T. K. Fujiwara, A. Kusumi, Transient GPI-anchored protein homodimers are units for raft organization and function. Nat. Chem. Biol. 8, 774–83 (2012).

132. A. Nakamura, C. Oki, K. Kato, S. Fujinuma, G. Maryu, K. Kuwata, T. Yoshii, M. Matsuda, K. Aoki, S. Tsukiji, Engineering orthogonal, plasma membrane-specific SLIPT systems for multiplexed chemical control of signaling pathways in living single cells. ACS Chem. Biol. 15, 1004–1015 (2020).

133. H. Sun, R. Taneja, Analysis of transformation and tumorigenicity using mouse embryonic fibroblast cells. Methods Mol. Biol. 383, 303–310 (2007).

134. C. L. Harris, S. M. Hanna, M. Mizuno, D. S. Holt, K. J. Marchbank, B. P. Morgan, Characterization of the mouse analogues of CD59 using novel monoclonal antibodies: tissue distribution and functional comparison. Immunology 109, 117–126 (2003).

135. I. Štefanová, I. Hilgert, H. Krištofová, R. Brown, M. G. Low, V. Hořejši, Characterization of a broadly expressed human leucocyte surface antigen MEM-43 anchored in membrane through phosphatidylinositol. Mol. Immunol. 26, 153–161 (1989).

136. N. Madore, K. L. Smith, C. H. Graham, A. Jen, K. Brady, S. Hall, R. Morris, Functionally different GPI proteins are organized in different domains on the neuronal surface. EMBO J. 18, 6917–6926 (1999).

137. D. W. Mason, A. F. Williams, The kinetics of antibody binding to membrane antigens in solution and at the cell surface. Biochem. J. 187, 1–20 (1980).

138. J. H. Hanke, J. P. Gardner, R. L. Dow, P. S. Changelian, W. H. Brissette, E. J. Weringer, B. A. Pollok, P. A. Connelly, Discovery of a novel, potent, and Src family-selective tyrosine kinase inhibitor. J. Biol. Chem. 271, 695–701 (1996).

139. J. K. Slack-Davis, K. H. Martin, R. W. Tilghman, M. Iwanicki, E. J. Ung, C. Autry, M. J. Luzzio, B. Cooper, J. C. Kath, W. G. Roberts, J. T. Parsons, Cellular characterization of a novel focal adhesion kinase inhibitor. J. Biol. Chem. 282, 14845–14852 (2007).

140. A. Kosenko, N. Hoshi, A change in configuration of the calmodulin-KCNQ channel complex underlies Ca^2+^-dependent modulation of KCNQ channel activity. PLoS One 8, e82290 (2013).

141. K. A. Johnson, M. R. Budicini, N. Bhattarai, T. Sharma, S. Urata, B. S. Gerstman, P. P. Chapagain, S. Li, R. V. Stahelin, PI(4,5)P2 binding sites in the Ebola virus matrix protein VP40 modulate assembly and budding. J. Lipid Res. 65, 100512 (2024).

142. S. V. Ulianov, A. K. Velichko, M. D. Magnitov, A. V. Luzhin, A. K. Golov, N. Ovsyannikova, I. I. Kireev, A. S. Gavrikov, A. S. Mishin, A. K. Garaev, A. V. Tyakht, A. A. Gavrilov, O. L. Kantidze, S. V. Razin, Suppression of liquid–liquid phase separation by 1,6-hexanediol partially compromises the 3D genome organization in living cells. Nucleic Acids Res. 49, 10524–10541 (2021).

143. P. Zhou, R. S. Kasai, W. Fujita, T. A. Tsunoyama, H. Neyama, H. Ueda, T. Yokoyama, M. Sakamoto, S. Pigolotti, T. K. Fujiwara, A. Kusumi, Single-molecule characterization of opioid receptor heterodimers reveals soluble µ-δ dimer blocker peptide alleviates morphine tolerance. Nat. Commun. 16, 9859 (2025).

144. T. Fujiwara, K. Ritchie, H. Murakoshi, K. Jacobson, A. Kusumi, Phospholipids undergo hop diffusion in compartmentalized cell membrane. J. Cell Biol. 157, 1071–1082 (2002).

145. I. Koyama-Honda, K. Ritchie, T. Fujiwara, R. Iino, H. Murakoshi, R. S. Kasai, A. Kusumi, Fluorescence imaging for monitoring the colocalization of two single molecules in living cells. Biophys. J. 88, 2126–2136 (2005).

146. A. C. Shibata, T. K. Fujiwara, L. Chen, K. G. Suzuki, Y. Ishikawa, Y. L. Nemoto, Y. Miwa, Z. Kalay, R. Chadda, K. Naruse, A. Kusumi, Archipelago architecture of the focal adhesion: membrane molecules freely enter and exit from the focal adhesion zone. Cytoskeleton 69, 380–92 (2012).

147. S. J. Sahl, M. Leutenegger, M. Hilbert, S. W. Hell, C. Eggeling, Fast molecular tracking maps nanoscale dynamics of plasma membrane lipids. Proc. Natl Acad. Sci. USA 107, 6829–6834 (2010).

148. P. Zhou, T. A. Tsunoyama, R. S. Kasai, K. M. Hirosawa, Z. Kalay, A. Aladag, T. Fujiwara, S. Pigolotti, A. Kusumi, Single-molecule detection of transient dimerization of opioid receptors 1: Homodimers’ effect on signaling and internalization. bioRxiv, 2024.07.25.605080 (2024).

149. S. L. Veatch, B. B. Machta, S. A. Shelby, E. N. Chiang, D. A. Holowka, B. A. Baird, Correlation functions quantify super-resolution images and estimate apparent clustering due to over-counting. PLoS One 7, e31457 (2012).

150. K. Xu, G. Zhong, X. Zhuang, Actin, spectrin, and associated proteins form a periodic cytoskeletal structure in axons. Science 339, 452–456 (2013).

151. K. A. K. Tanaka, K. G. N. Suzuki, Y. M. Shirai, S. T. Shibutani, M. S. H. Miyahara, H. Tsuboi, M. Yahara, A. Yoshimura, S. Mayor, T. K. Fujiwara, A. Kusumi, Membrane molecules mobile even after chemical fixation. Nat Methods 7, 865–866 (2010).

152. S.-H. Lee, J. Y. Shin, A. Lee, C. Bustamante, Counting single photoactivatable fluorescent molecules by photoactivated localization microscopy (PALM). Proc. Natl Acad. Sci. USA 109, 17436–17441 (2012).

153. E. D. Zitter, D. Thédié, V. Mönkemöller, S. Hugelier, J. Beaudouin, V. Adam, M. Byrdin, L. V. Meervelt, P. Dedecker, D. Bourgeois, Mechanistic investigation of mEos4b reveals a strategy to reduce track interruptions in sptPALM. Nat. Methods 16, 707–710 (2019).

154. M. Ovesný, P. Křížek, J. Borkovec, Z. Švindrych, G. M. Hagen, ThunderSTORM: a comprehensive ImageJ plug-in for PALM and STORM data analysis and super-resolution imaging. Bioinformatics 30, 2389–2390 (2014).

155. S. Berg, D. Kutra, T. Kroeger, C. N. Straehle, B. X. Kausler, C. Haubold, M. Schiegg, J. Ales, T. Beier, M. Rudy, K. Eren, J. I. Cervantes, B. Xu, F. Beuttenmueller, A. Wolny, C. Zhang, U. Koethe, F. A. Hamprecht, A. Kreshuk, ilastik: interactive machine learning for (bio)image analysis. Nat. Methods 16, 1226–1232 (2019).

156. M. Dundr, T. Misteli, Measuring dynamics of nuclear proteins by photobleaching. Curr. Protoc. Cell Biol. 18, 13.5.1–13.5.18 (2003).

157. N. Otsu, A threshold selection method from gray-level histograms. IEEE Trans. Syst. Man Cybern. 9, 62–66 (1979).

158. N. J. Anthis, G. M. Clore, Sequence-specific determination of protein and peptide concentrations by absorbance at 205 nm: sequence-specific protein concentration at 205 nm. Protein Sci. 22, 851–858 (2013).

159. M. M. Tomayko, C. P. Reynolds, Determination of subcutaneous tumor size in athymic (nude) mice. Cancer Chemother. Pharmacol. 24, 148–154 (1989).

160. C. Tan, J. Jung, C. Kobayashi, D. U. L. Torre, S. Takada, Y. Sugita, Implementation of residue-level coarse-grained models in GENESIS for large-scale molecular dynamics simulations. PLoS Comput. Biol. 18, e1009578 (2022).

161. J. Jung, K. Yagi, C. Tan, H. Oshima, T. Mori, I. Yu, Y. Matsunaga, C. Kobayashi, S. Ito, D. U. L. Torre, Y. Sugita, GENESIS 2.1: High-performance molecular dynamics software for enhanced sampling and free-energy calculations for atomistic, coarse-grained, and quantum mechanics/molecular mechanics models. J. Phys. Chem. B 128, 6028–6048 (2024).

162. J. Jung, C. Tan, Y. Sugita, GENESIS CGDYN: large-scale coarse-grained MD simulation with dynamic load balancing for heterogeneous biomolecular systems. Nat. Commun. 15, 3370 (2024).

163. J. Schindelin, I. Arganda-Carreras, E. Frise, V. Kaynig, M. Longair, T. Pietzsch, S. Preibisch, C. Rueden, S. Saalfeld, B. Schmid, J.-Y. Tinevez, D. J. White, V. Hartenstein, K. Eliceiri, P. Tomancak, A. Cardona, Fiji: an open-source platform for biological-image analysis. Nat. Methods 9, 676–682 (2012).

164. U. Praekelt, P. M. Kopp, K. Rehm, S. Linder, N. Bate, B. Patel, E. Debrand, A. M. Manso, R. S. Ross, F. Conti, M.-Z. Zhang, R. C. Harris, R. Zent, D. R. Critchley, S. J. Monkley, New isoform-specific monoclonal antibodies reveal different sub-cellular localisations for talin1 and talin2. Eur. J. Cell Biol. 91, 180–191 (2012).

165. G. D. Paolo, L. Pellegrini, K. Letinic, G. Cestra, R. Zoncu, S. Voronov, S. Chang, J. Guo, M. R. Wenk, P. D. Camilli, Recruitment and regulation of phosphatidylinositol phosphate kinase type 1γ by the FERM domain of talin. Nature 420, 85–89 (2002).

166. I. de Curtis, Biomolecular condensates at the front: cell migration meets phase separation. Trends Cell Biol. 31, 145–148 (2021).

167. C.-C. Lin, K. M. Suen, J. Lidster, J. E. Ladbury, The emerging role of receptor tyrosine kinase phase separation in cancer. Trends Cell Biol. 34, 371–379 (2024).

168. K. Graham, A. Chandrasekaran, L. Wang, N. Yang, E. M. Lafer, P. Rangamani, J. C. Stachowiak, Liquid-like condensates mediate competition between actin branching and bundling. Proc. Natl. Acad. Sci. USA 121, e2309152121 (2024).

169. Y. Wang, C. Zhang, W. Yang, S. Shao, X. Xu, Y. Sun, P. Li, L. Liang, C. Wu, LIMD1 phase separation contributes to cellular mechanics and durotaxis by regulating focal adhesion dynamics in response to force. Dev. Cell 56, 1313–1325.e7 (2021).

170. P. Liang, Y. Wu, S. Zheng, J. Zhang, S. Yang, J. Wang, S. Ma, M. Zhang, Z. Gu, Q. Liu, W. Jiang, Q. Xing, B. Wang, Paxillin phase separation promotes focal adhesion assembly and integrin signaling. J. Cell Biol. 223, e202209027 (2024).

171. J. Zhu, Q. Zhou, Y. Xia, L. Lin, J. Li, M. Peng, R. Zhang, M. Zhang, GIT/PIX condensates are modular and ideal for distinct compartmentalized cell signaling. Mol. Cell 79, 782–796.e6 (2020).

172. T. Litschel, C. F. Kelley, X. Cheng, L. Babl, N. Mizuno, L. B. Case, P. Schwille, Membrane-induced 2D phase separation of the focal adhesion protein talin. Nat. Commun. 15, 4986 (2024).

173. C. M. Lim, A. G. Díaz, M. Fuxreiter, F. W. Pun, A. Zhavoronkov, M. Vendruscolo, Multiomic prediction of therapeutic targets for human diseases associated with protein phase separation. Proc. Natl. Acad. Sci. USA 120, e2300215120 (2023).

174. A. Tulpule, J. Guan, D. S. Neel, H. R. Allegakoen, Y. P. Lin, D. Brown, Y.-T. Chou, A. Heslin, N. Chatterjee, S. Perati, S. Menon, T. A. Nguyen, J. Debnath, A. D. Ramirez, X. Shi, B. Yang, S. Feng, S. Makhija, B. Huang, T. G. Bivona, Kinase-mediated RAS signaling via membraneless cytoplasmic protein granules. Cell 184, 2649–2664. e18 (2021).

175. K. Zhou, Q. Chen, J. Chen, D. Liang, W. Feng, M. Liu, Q. Wang, R. Wang, Q. Ouyang, C. Quan, S. Chen, Spatiotemporal regulation of insulin signaling by liquid–liquid phase separation. Cell Discov. 8, 64 (2022).

176. X. K. Gao, X. S. Rao, X. X. Cong, Z. K. Sheng, Y. T. Sun, S. B. Xu, J. F. Wang, Y. H. Liang, L. R. Lu, H. Ouyang, H. Ge, J. Guo, H. Wu, Q. M. Sun, H. Wu, Z. Bao, L. L. Zheng, Y. T. Zhou, Phase separation of insulin receptor substrate 1 drives the formation of insulin/IGF-1 signalosomes. Cell Discov. 8, 60 (2022).

177. J. Sampson, M. W. Richards, J. Choi, A. M. Fry, R. Bayliss, Phase-separated foci of EML4-ALK facilitate signalling and depend upon an active kinase conformation. EMBO Rep. 22, EMBR202153693 (2021).

178. L. Zeng, I. Palaia, A. Šarić, X. Su, PLCγ1 promotes phase separation of T cell signaling components. J. Cell Biol. 220, e202009154 (2021).

179. N. Ma, F. Wu, J. Liu, Z. Wu, L. Wang, B. Li, Y. Liu, X. Dong, J. Hu, X. Fang, H. Zhang, D. Ai, J. Zhou, X. Wang, Kindlin-2 phase separation in response to flow controls vascular stability. Circ. Res. 135, 1141–1160 (2024).

180. K. Sala, A. Corbetta, C. Minici, D. Tonoli, D. H. Murray, E. Cammarota, L. Ribolla, M. Ramella, R. Fesce, D. Mazza, M. Degano, I. de Curtis, The ERC1 scaffold protein implicated in cell motility drives the assembly of a liquid phase. Sci. Rep. 9, 13530 (2019).

181. K. Sala, A. Raimondi, D. Tonoli, C. Tacchetti, I. de Curtis, Identification of a membrane-less compartment regulating invadosome function and motility. Sci. Rep. 8, 1164 (2018).

182. K. Guo, J. Zhang, P. Huang, Y. Xu, W. Pan, K. Li, L. Chen, L. Luo, W. Yu, S. Chen, S. He, Z. Wei, C. Yu, KANK1 shapes focal adhesions by orchestrating protein binding, mechanical force sensing, and phase separation. Cell Rep. 42, 113321 (2023).

183. D. A. Calderwood, I. D. Campbell, D. R. Critchley, Talins and kindlins: partners in integrin-mediated adhesion. Nat. Rev. Mol. Cell Biol. 14, 503–517 (2013).

184. J. Yang, L. Zhu, H. Zhang, J. Hirbawi, K. Fukuda, P. Dwivedi, J. Liu, T. Byzova, E. F. Plow, J. Wu, J. Qin, Conformational activation of talin by RIAM triggers integrin-mediated cell adhesion. Nat. Commun. 5, 5880 (2014).

185. M. Krause, E. W. Dent, J. E. Bear, J. J. Loureiro, F. B. Gertler, ENA/VASP proteins: regulators of the actin cytoskeleton and cell migration. Cell Dev. Biol. 19, 541–564 (2003).

186. A. M. López-Colomé, I. Lee-Rivera, R. Benavides-Hidalgo, E. López, Paxillin: a crossroad in pathological cell migration. J. Hematol. Oncol. 10, 50 (2017).

187. T. Takenawa, S. Suetsugu, The WASP–WAVE protein network: connecting the membrane to the cytoskeleton. Nat. Rev. Mol. Cell Biol. 8, 37–48 (2007).

188. F. J. Sulzmaier, C. Jean, D. D. Schlaepfer, FAK in cancer: mechanistic findings and clinical applications. Nat. Rev. Cancer 14, 598–610 (2014).

189. M. E. Amri, U. Fitzgerald, G. Schlosser, MARCKS and MARCKS-like proteins in development and regeneration. J. Biomed. Sci. 25, 43 (2018).

190. J. D.- Guercio, L. Kurzawa, J. Mueller, G. Dimchev, M. Schaks, M. Nemethova, T. Pokrant, S. Brühmann, J. Linkner, L. Blanchoin, M. Sixt, K. Rottner, J. Faix, Loss of Ena/VASP interferes with lamellipodium architecture, motility and integrin-dependent adhesion. eLife 9, e55351 (2020).

