## Supplementary materials for "A nano-liquid hub integrating growth and immune-evasion signaling to promote cell survival"

**The PDF file includes:**

Materials and Methods
Figs. S1 to S14
References

**Other Supplementary Materials for this manuscript include the following:**

Movies S1 to S5
Data S1 to S2

### Materials and Methods

Cell culture

T24 cells (a kind gift from Masahiro Sokabe, Nagoya University) (*109*), HeLa cells (RIKEN Cell Bank, RCB0007), and CHO-K1 cells (Dainippon Pharma Co., Ltd., 03-402) were cultured in Ham's nutrient mixture F12 (Sigma-Aldrich, N6658) supplemented with 10% (vol/vol) fetal bovine serum (FBS; Gibco, 10270-106), 100 U/ml penicillin, and 100 μg/ml streptomycin (Nacalai Tesque, 26253-84). The talin1 and 2 double knockout mouse embryonic fibroblasts (talin-dKO MEFs) and talin2-KO MEFs (kind gifts from Reinhard Fässler, Max Planck Institute of Biochemistry) (*19*), the zyxin-KO MEF line (a kind gift from Mary C. Beckerle, University of Utah) (*22*), control MEF lines 1-3 (kind gifts from Noboru Mizushima, University of Tokyo), control MEF line 4 (a kind gift from Ryusuke Kuwahara, OIST), NIH3T3 cells (a kind gift from Makoto Kinoshita, Nagoya University) (*110*), and MDA-MB231 cells (American Type Culture Collection, ATCC, CRM-HTB-26) were cultured in Dulbecco's modified Eagle's minimum essential medium (DMEM; Sigma-Aldrich, D0822) supplemented with 10% (vol/vol) FBS, 100 U/ml penicillin, and 100 μg/ml streptomycin. RKO cells (ATCC, CRL-2577) were cultured in Eagle's minimum essential medium (MEM; Gibco, 11095080) supplemented with 10% (vol/vol) FBS, 100 U/ml penicillin, and 100 μg/ml streptomycin. Mouse embryonic stem cells (mESCs; clone E14tg2a; RIKEN Cell Bank, AES0135) (*111*) were cultured on gelatin-coated dishes in DMEM supplemented with 15% (vol/vol) embryonic stem-cell qualified FBS (Gibco, 16141079), 1x MEM non-essential amino acids solution (Gibco, 11140050), 1 mM sodium pyruvate (Gibco, 11360070), 55 μM 2-mercaptoethanol (Gibco, 21985023), 100 U/ml penicillin, 100 μg/ml streptomycin, and $10^3$ U/ml mouse leukemia inhibitory factor (LIF; Millipore, ESG1106), without feeder cells (*112*). The identities of the T24 and HeLa cell lines were authenticated by PowerPlex16 STR (contracted to Promega). All cells were confirmed to be mycoplasma-free using a Mycoplasma PCR Detection Kit (Beyotime Biotechnology, C0301S) and VenorGeM Mycoplasma Detection Kit (Minerva Biolabs, MNV11-1100).

cDNA construction

Plasmids were constructed using standard molecular biology techniques and verified by Sanger sequencing. All plasmids used in this study are listed, and the complete sequence information in GenBank format is available in Data S1. The cDNAs encoding SNAPf (N9172S), Halo7 (G9651), mGFP (Z2370N), and mCherry (Z2522N) were obtained from New England Biolabs, Promega, and Clontech, respectively. The cDNA encoding mEos3.2 was generated by introducing three point mutations, I102N, H158E, and Y189A, into the mEos2 plasmid, a kind gift from Loren L. Looger (Janelia Research Campus; Addgene, 20341) (*113*–*115*). The cDNAs encoding deletion mutants of RIAM, VASP, and zyxin were generated by PCR using designed primers. To construct cDNAs encoding phosphomimetic and phosphodeficient mutants of zyxin and VASP, all possible phosphorylation sites were identified by PhosphoSitePlus (https://www.phosphosite.org, 34 sites for zyxin and 27 sites for VASP). All phosphorylatable serine, threonine, and tyrosine residues were replaced with glutamic acid for phosphomimetic mutations, and with alanine, alanine, and phenylalanine, respectively, for phosphodeficient mutations. The mutated cDNAs were custom synthesized by Gene Universal. The cDNAs encoding mScarlet-I (*116*), domain 4 from perfringolysin O with the D434S mutation (D4H) (*117*, *118*), and ostreolysin A with the E69A mutation (OlyA [E69A]) (*29*) were custom synthesized by Sangon Biotech (Shanghai, PRC).

The cDNAs encoding human talin1 (ORK01622), human zyxin (ORS08391), human VASP (FHC28647), human vinculin (FXC01835), human PIK3CA (FXC01204), human MARCKS (FXC03149), human Akt (FXC02215), and human JAK1 (FXC01306) were purchased from Kazusa DNA Research Institute. The cDNAs encoding human α-actinin (Aact-138) and human filamin A (FILA-106) were purchased from Allele Biotechnology. The cDNAs encoding human paxillin isoform alpha (*119*) and rat Thy1 (*120*) were cloned from the WI38 cell line and the rat brain, respectively.

The following cDNAs were kind gifts: mouse FAK from Caroline H. Damsky (University of California, San Francisco) (*121*); human RIAM from Chinten J. Lim (University of California, San Diego; Addgene, 80028) (*122*); human β3 integrin from Jonathan C. Jones (Northwestern University) (*123*); rat N-WASP from Jack Taunton (University of California, San Francisco) (*124*); mouse PIP5KIγ from Richard A. Anderson (University of Wisconsin) (*125*); human CD59 from Masahide Tone (University of Oxford) (*126*); human PDGFR-β from William Hahn (Broad Institute of MIT) and David Root (Harvard University; Addgene, 23893) (*127*); rat PLCγ1 from Matilda Katan (Cancer Research UK Centre) (*128*); mouse PLCβ1 from Katsuhiko Mikoshiba (ShanghaiTech University) (*129*); and human CD47 from Eric C. Brown (Genentech) (*130*); probes for PI(4)P, PI(3,4)$P_2$, PI(4,5)$P_2$, and PI(3,4,5)$P_3$ from Toshiki Itoh (Kobe University) (*32*); a probe for phosphatidylserine from Tomohiko Taguchi (Tohoku University) (*30*, *31*); CIBN and CRY2-5ptase from Pietro de Camilli (Yale University; Addgene, 79574, 66836, and 66837) (*33*); membrane-optoDroplet from Jared E. Toettcher (Princeton University; Addgene, 111509) (*69*); tag-FAK-B and binder from Klaus M. Hahn (University of North Carolina; Addgene, 179142) (*67*).

A five-amino acid linker sequence (Ser-Gly-Gly-Gly-Gly) was inserted between the target protein and mGFP, mCherry, and mScarlet-I, and a 15-amino acid linker ($(\text{Ser-Gly-Gly-Gly-Gly})_3$) was inserted between the target protein and SNAPf, Halo7, and mEos3.2.

The sequences encoding these proteins were subcloned into the following vectors: pEGFP and pmCherry (Clontech, Z2370N and Z2522N), pcDNA3.1(+)-Hygro and pcDNA3.1(+)-Zeo (Invitrogen, V87020 and V86020), pET26B (Novagen, 69862), pOStet15T3 (an episomal Epstein–Barr virus-based vector carrying tetracycline-regulated expression units, including the transactivator (rtTA2-M2) and the tetO sequence [Tet-on vector]) from Yoshihiro Miwa (University of Tsukuba) (*131*); and pPBpuro (a PiggyBac-based custom vector containing the terminal repeat sequence and a strong composite promoter consisting of chicken β-actin promoter, CAG promoter with an artificial intron, and CMV immediate early promoter) and pCMV-mPBase from Michiyuki Matsuda (Kyoto University) (*132*);

Cell transfection

Cells were transfected with plasmid vectors by electroporation, using a Lonza 4D-Nucleofector according to the manufacturer's instructions (Lonza, V4XC-2012 and V4XC-1012). The following solutions and programs were used: for T24 and NIH3T3 cells, SF solution and program EN138; for HeLa cells, SE and CN114; for CHO-K1 cells, SF and DT133; for MDA-MB231 cells, SE and EW113; for RKO cells, SE and DN100; for talin-dKO MEFs, SE and DS150; for other MEFs, SF and CZ167; and for mESCs, SF and CG104.

For the generation of stable cell lines expressing selected proteins, cells were transfected with an expression plasmid. When using the pPBpuro expression vector (PiggyBac-based vector), pCMV-mPBase was co-transfected. Two days after transfection, selection was initiated using appropriate antibiotics. G418 was used at 800 µg/mL for T24 cells, 1,000 µg/mL for MEFs, 5,000 µg/mL for MDA-MB231 cells, and 2,000 µg/mL for RKO cells. Puromycin was used at 0.5 µg/mL for T24, RKO, and MDA-MB231 cells, and at 1 µg/mL for MEFs. Hygromycin B was used at either 500 or 1,000 µg/mL for MEFs, and zeocin was used at 500 µg/mL for MEFs. After one week of selection, cell clones were established either by limiting dilution using 96-well plates or by fluorescence-activated cell sorting (FACS) using an Aria II (BD Biosciences) to isolate populations with appropriate expression levels. Expression levels of the introduced constructs were examined by western blotting and immunofluorescence microscopy. The established stable cell lines were subsequently maintained in culture medium containing the corresponding selection antibiotics.

For the expression of CRY2-fused proteins, transfected talin1/2-dKO MEFs were first incubated in complete growth medium at 37ºC for 2 h for post-transfection recovery and then cultured at 33ºC for 2 days to promote protein folding, followed by an incubation at 37ºC for 3 h immediately prior to microscope observations.

To generate malignantly transformed MEF lines for xenograft studies, zyxin-KO MEFs (2 x $10^4$ cells) were transfected with a lentivirus vector encoding the human K-Ras G12V mutant (5 x $10^5$ inclusion-forming unit; GenTarget, LVP1139-RB). Two days after infection, cells were selected with 5 µg/ml blasticidin and continuously cultured under selection for one month to ensure complete transformation (*133*); Because RFP is fused to the blasticidin resistance gene, K-Ras (G12V) expression levels could be monitored by RFP fluorescence. To normalize Kras (G12V) expression, we selected the clones exhibiting comparable RFP fluorescence intensities.

<u>Cell culture for microscope observations</u>

Cells were plated on glass-bottom dishes (12-mm diameter glass window; No.1S; Iwaki, 3971-035) pre-coated with fibronectin (FN; Sigma-Aldrich, F1141). For coating, 100 µL of FN (10 µg/mL in phosphate buffered saline; PBS) was placed onto the glass surface and incubated at room temperature for 30 min. PBS contained 137 mM NaCl, 8.1 mM $NaHPO_4$, 2.7 mM KCl, and 1.5 mM $KH_2PO_4$ (pH 7.4). After seeding, cells were cultured in the complete growth medium for 24–48 h before microscopy.

<u>Lentiviral production and infection for shRNA gene knockdown</u>

Mice possess two homologous CD59 genes, CD59a and CD59b. Since CD59a is broadly expressed in most tissues, whereas CD59b is restricted to germ cells (*134*), gene knockdown was performed against CD59a for transformed MEFs in this study.

The oligonucleotide pair targeting mouse CD59a mRNA (5'-CCGGCGGTGGTTTCTTCATGCAATACTCGAGTATTGCATGAAGAAACCACCGTTTTTG-3' and 5'-AATTCAAAAACGGTGGTTTCTTCATGCAATACTCGAGTATTGCATGAAGAAACCACCG-3') and its scrambled control pair (5'-CCGGGATGACTAGTCGTTATGCTCTCTCGAGAGAGCATAACGACTAGTCATCTTTTTG-3' and 5'-AATTCAAAAAGATGACTAGTCGTTATGCTCTCTCGAGAGAGCATAACGACTAGTCATC-3') were cloned into the pLKO1-hygro vector (Addgene, 24150; deposited by Robert A. Weinberg, MIT) at the AgeI – EcoRI site. The shRNA constructs were co-transfected with packaging plasmids (Lentiviral High Titer Packing Mix; Takara, 6194) into Lenti-X 293T cells (Takara, 632180), using TransIT-293 (Mirus, MIR2700). At 24 h after transfection, the medium was replaced with fresh complete growth medium (DMEM + 10%FBS). At 48 h after transfection, the lentiviral supernatant was harvested and filtered through a 0.45-µm Millex filter (Millipore, SLFH05010). The lentiviral supernatant was concentrated 50x in complete growth medium, using a Lenti-X Concentrator (Takara, 631232), and the concentrated lentiviral solution was stored at -80ºC until use.

One day before infection, zyxin KO transformed MEFs were plated in 12-well plastic plates at 2.5 x $10^4$ cells per well in 1 mL medium. The concentrated lentiviral solution (4 µL) was added to each well, and the cells were incubated for 24 h. The medium was then replaced with fresh complete growth medium, and the cells were further incubated for 48 h. After selection in medium containing 1 mg/mL hygromycin for one week, FACS was used to isolate a cell population with low CD59a expression. For sorting, a mouse anti-CD59a antibody (clone mCD59.3; BioLegend, 143104) was used at 10 µg/ml. The CD59a expression levels were quantified by western blotting.

CRISPR/Cas9-mediated knockout of zyxin in MDA-MB231 and RKO cells

The synthetic guide RNA (sgRNA) targeting ZYX (Invitrogen, CRISPR865844_SGM; targeting sequence: 5' TACGCCCCGCAGAAGAAGTT 3') and Cas9 protein (Invitrogen, A36498), 38 pmol each, were mixed in 20 µL of SE solution (Lonza) and incubated for 30 min at RT. Approximately 2 x $10^5$ cells were resuspended in this solution, and the sgRNA/Cas9 complex was delivered by electroporation using a Lonza 4D-Nucleofector (program EW113 for MDA-MB231 cells and program DN100 for RKO cells). Two days after electroporation, clones were isolated by limiting dilution in 96-well plates. Clones were validated by western blotting and amplicon sequencing of the target genomic locus, using the primers 5' CCGCTCCTTGTATGGTCTGG 3' (forward) and 5' CAAATTCCCGAAACGACTGC 3' (reverse) with a minimum of 24 sequencing reads per clone.

Cell stimulation

CD59 and Thy1 were activated by polystyrene beads coated with the respective monoclonal antibodies. Mouse anti-CD59 and mouse anti-Thy1 monoclonal antibodies were purified from the culture media of hybridoma MEM43/5 cells (a kind gift from Vaclav Horejsi, Institute of Molecular Genetics, Czech Academy of Sciences) (*135*) and hybridoma OX7 cells (European Collection of Cell

Cultures, 84112008) (136, 137), respectively. The carboxylate-modified yellow-green or dark-red fluorescent 40-nm polystyrene beads (Molecular Probes, F8795 and F8789; final concentration, 0.4 μM) were functionalized by an incubation at RT for 10 min in activation solution, containing 2 mM 1-ethyl-3-(3-dimethylaminopropyl)carbodiimide (EDC; Pierce, 22980), 4 mM sulfo-N-hydroxysuccinimide (sulfo-NHS; Pierce, 24525), and 150 mM NaCl, buffered with 0.1 M 2-(N-morpholino)ethanesulfonic acid (MES; pH 5.5). A 9 μL aliquot of antibody solution (1 mg/ml in PBS) was mixed with 3 μL of activated beads, 3 μL of borate-buffered saline (150 mM NaCl buffered with 0.5 M borate, pH 8.0), and 1 μL of 0.4 M NaOH. After an incubation at RT for 1 h, the coupling reaction was stopped by adding 7.5 μL of 300 mM glycine and 1 μL of 5% polyethylene glycol (mean molecular weight 20,000, PEG20K; Sigma-Aldrich, P5413) in PBS, followed by further incubation at RT for 5 min. The mixture was ultracentrifuged at 50,000 rpm for 15 min. The antibody-coated beads were resuspended in 1 mL PBS containing 0.1% PEG20K, ultracentrifuged again, and finally resuspended in Hanks' balanced salt solution (HBSS; Nissui, 05905) buffered with 2 mM TES (Dojindo, 348-08371) at pH 7.4 (observation medium; T-HBSS) containing 2% bovine serum albumin (BSA, Sigma-Aldrich, A3059) and 0.1% PEG20K. Immediately before use, the bead suspension was sonicated for 5 min and centrifuged at 15,000 rpm for 10 min. Cells were first incubated in T-HBSS containing 0.5% BSA and 0.1% PEG20K, after which the bead suspension was added at a final concentration of 2 μM or 20 nM. All observations were completed within 10 min after bead addition to minimize the effects of internalization.

For cell stimulation experiments using epidermal growth factor (EGF) and platelet-derived growth factor (PDGF), cells were serum-starved in FBS-free medium for 3-12 h. Prior to cell stimulation and microscopy, the culture medium was replaced with T-HBSS, and cells were stimulated with 20 nM human recombinant EGF (Wako, 053-07871), and 20 nM or 20 pM human recombinant PDGF-BB (Pepro Tech, 100-14B). All microscopy observations were completed within 5 min after stimulation. When simultaneous stimulations with anti-CD59 beads and PDGF were performed, the bead buffer (T-HBSS containing BSA and PEG20K) was used, and all observations were completed within 10 min after stimulation.

Cell treatments with various reagents

Kinase activities were inhibited by incubating cells in T-HBSS containing 10 μM PP2 (Src family kinase inhibitor; Selleckchem, S7008) (138) or 10 μM PF573228 (focal adhesion kinase inhibitor; Cayman Chemical, 14924) (139) at 37°C for 10 min.

Partial actin depolymerization was performed by incubating cells in T-HBSS containing 100 nM latrunculin A (Wako, 125-04363), 500 nM cytochalasin D (Cayman Chemical, 11330), or 10 μM Y27632 (ChemScene, CS-0878) at 37°C for 10 min. Partial depletion of cholesterol was achieved by incubating cells in 4 mM methyl-β-cyclodextrin (MβCD; Sigma-Aldrich, 332615) in T-HBSS at 37°C for 30 min or in 60 μg/ml saponin (Nacalai Tesque, 30502-42) in T-HBSS on ice for 15 min (131). Replenishment of cholesterol was performed by incubating cholesterol-depleted cells in 10 mM MβCD-cholesterol complex (1:1 mol/mol; Sigma-Aldrich, C3045) in T-HBSS at 37°C for 30 min (131). Partial depletion of $PI(4,5)P_2$ from the PM was performed by incubating cells with either

10 µM ionomycin (Wako, 093-04531) in T-HBSS at 37°C for 5 min (*140*), 1 µM phenylarsine oxide (Tokyo Chemical Industry, P0140) at 37°C for 3 h, or 50 µM quercetin (Sigma-Aldrich, Q4951) at 37°C for 3 h (*34*, *141*).

To inhibit interactions mediated by intrinsically disordered regions (IDRs), cells were first mildly permeabilized with 1% Tween 20 in T-HBSS at RT for 10 min, and then treated with T-HBSS containing 5% 1,6-hexanediol (Wako, 081-00435) on the microscope stage during imaging (*142*). All imaging experiments were completed within 10 min. For *in vitro* experiments, 1,6-hexanediol was added to a final concentration of 10%.

Western blotting

For cultured cell samples, ice-cold radio immunoprecipitation assay (RIPA) buffer was used (350 µL for 60-mm plastic dish). The RIPA buffer consisted of 150 mM NaCl, 5 mM ethylenediaminetetraacetic acid (EDTA; Dojindo, 345-01865), 1% NP-40 (Wako, 141-08321), 1% sodium deoxycholate (Nacalai Tesque, 08805-72), 0.1% sodium dodecyl sulfate (SDS; Wako, 191-07145), and 1x protease inhibitor cocktail (Nacalai Tesque, 25955-24), buffered with 50 mM tris(hydroxymethyl)aminomethane (Tris)-HCl (pH 7.4). For the Akt phosphorylation assay, 1x phosphatase inhibitor cocktail (Nacalai Tesque, 07574-61) was included. The RIPA buffer was added to cells at ≈80% confluency in a 60-mm plastic dish, and cells were kept on ice for 5 min. Lysates were collected using a cell scraper and centrifuged at 15,000 rpm for 20 min.

For tumor tissue samples, homogenization was performed by two cycles of freezing and thawing, pestle grinding, and sonication (50% duty for 5 min on ice) in RIPA buffer. Insoluble fractions were removed by centrifugation at 15,000 rpm for 1 h.

The supernatant (300 µL) was mixed with 100 µL of 4x Laemmli sample buffer (8% SDS, 0.12% bromophenol blue, 40% glycerol, and 400 mM dithiothreitol (DTT), buffered with 200 mM Tris-HCl, pH 6.8) and boiled for 5 min. For CD59 detection, samples were prepared under non-reducing conditions (without DTT and boiling), because the anti-CD59 antibody only binds to non-reduced proteins.

Samples (2-10 µL; 1-2 µg of total protein per lane) were loaded onto precast 4-20% gradient polyacrylamide gels (Bio-Rad, 4561096) and electrophoresed at a constant voltage of 200 V for 25 min (CD59) or 35 min (others) using a Mini Protean Tetra Cell (Bio-Rad, 1658000JA), in Tris-glycine running buffer (25 mM Tris-HCl, 190 mM glycine, and 0.1% SDS). Direct addition of 4x Laemmli buffer to the specimens yielded comparable results but increased the background; therefore, the specimens were first solubilized with RIPA buffer.

Proteins were transferred to polyvinylidene fluoride (PVDF) membranes (0.45 µm pore size) by semi-dry transfer using a Trans-Blot Turbo (Bio-Rad, mixed MW protocol) with the PVDF transfer pack (Bio-Rad, 1704156) for zyxin, VASP, Akt, FAK, and JAK1. For CD59, proteins were transferred to PVDF membranes (0.2 µm pore size; FluoroTransW; Pall, BSP0161) using the transfer buffer (25 mM Tris-HCl, 190 mM glycine, and 20% methanol) at RT at a constant voltage of 50 V for 10 min (Trans-Blot Turbo). For talin, wet transfer to the PVDF membrane (Immobilon-P;

Millipore, IPVH00010) was performed using the same transfer buffer at 4ºC at a constant voltage of 100 V for 1 h (Mini Protean Tetra Cell).

Membranes were blocked with Blocking One (Nacalai Tesque, 03953-95) for 30 min at RT and washed with Tris-buffered saline (TBS; 137 mM NaCl and 2.7 mM KCl buffered with 25 mM Tris-HCl, pH 7.4) containing 0.1% Tween 20 (TBS-T). The blocked membranes were incubated with primary antibodies at RT for 60 min: mouse anti-talin1 (clone 93E12; Abcam, ab104913; 0.5 μg/mL), rabbit anti-zyxin (B71; Millipore, ABC1463; 1:4,000), rabbit anti-GFP (ProteinTech, 50430-2-AP; 0.5 μg/mL), rabbit anti-VASP (clone 9A2; Cell Signaling Technology, 3132; 21 ng/mL), rabbit anti-Akt (Cell Signaling Technology, 9272; 17 ng/mL), rabbit anti-phospho-Akt (pS473; clone D9E; Cell Signaling Technology, 4060; 48 ng/mL), rabbit anti-FAK (Cell Signaling Technology, 3285; 31 ng/mL), rabbit anti-phospho-FAK (pY397; clone D20B1; Cell Signaling Technology, 8556; 8 ng/mL), rabbit anti-JAK1 (Cell Signaling Technology, 3332; 1:1,000), rabbit anti-phospho-JAK1 (pY1034/1035; Cell Signaling Technology, 3331; 82 ng/mL), mouse anti-CD59a (clone mCD59.3, 1 μg/mL) and mouse anti-GAPDH (ProteinTech, 60004-1-Ig; 10-500 ng/mL). Membranes were then incubated with secondary antibodies at RT for 30 min: horseradish peroxidase (HRP)-conjugated goat anti-mouse immunoglobulin G (IgG; Invitrogen, G21040; 0.5 μg/mL) and HRP-conjugated goat anti-rabbit IgG (Invitrogen, G21234; 0.5 μg/mL).

Membranes were incubated with Western BLoT Hyper HRP Substrate (Takara, T7103B) for chemiluminescence detection and imaged using an LAS-3000 (Fuji Film, high sensitivity mode) or iBright FL1500 (Thermo Fischer Scientific; 2x2 binning, 2.0x zoom). Quantification was performed using ImageJ, with signal intensities normalized to GAPDH. For tumor tissues, where GAPDH intensities varied, normalization was based on total protein amounts measured from Coomassie Brilliant Blue R-250 (CBB R-250)-stained gels. For CD59 detection under non-reducing conditions, GAPDH could not be used (the anti-GAPDH antibody binds only to reduced proteins); therefore, normalization was performed using CBB R-250–stained gels.

For phosphorylation analysis, membranes were first probed and imaged for phosphorylated proteins, and then stripped and reprobed with antibodies against total protein. For Akt and JAK1, stripping was performed in buffer consisting of 2% SDS, 100 mM 2-mercaptoethanol, and 62.5 mM Tris-HCl (pH 6.7) at 50°C for 20 min. For FAK, stripping was performed with WB Stripping Solution (Nacalai Tesque, 05364-55) at RT for 30 min.

Cell attachment assay

Cells at ≈80% confluency in 100-mm plastic dishes were detached with freshly prepared trypsin solution (0.25% trypsin and 1 mM EDTA in PBS) at RT for 3 min. After brief centrifugation (1,000 rpm for 3 min), cells were resuspended in the complete growth medium, and cell numbers were determined using a hemocytometer. The cell concentration was adjusted to 3 x $10^4$ cells/mL, and cells were incubated at 37ºC, 5% $CO_2$ for 30 min to allow recovery after detachment. Subsequently, 1 mL of the cell suspension was dispensed into each FN-coated well of a 12-well plastic plate, and the plate was incubated at 37ºC, 5% $CO_2$ for 5 min. Wells were gently washed twice with T-HBSS, and the attached cells were imaged using a Zeiss Primovert Photo inverted phase-contrast microscope. The number of attached cells was quantified using ImageJ.

Immunofluorescence labeling

Cultured cells on FN-coated glass-base dishes were fixed with cold methanol (-20ºC) for 5 min for talin staining or with 4% paraformaldehyde at RT for 1 h for zyxin staining. Fixed cells were treated sequentially at RT with 100 mM glycine for 15 min, 0.1% Triton X-100 for 5 min, and 1% BSA for 1 h. All solutions were prepared in PBS containing 0.1% Tween 20 (PBS-T). Cells were immunostained by sequential 1-h incubations with primary and secondary antibodies at RT, with three PBS-T washes after each incubation.

The antibodies used were: mouse anti-talin1 (clone 97H6; Bio-Rad, MCA4770; 1 μg/mL), rabbit anti-zyxin (B71, 1:600), Rhodamine Red-X-conjugated donkey anti-mouse IgG (Jackson ImmunoResearch, 715-295-150; 1 μg/mL), and Rhodamine Red-X-conjugated donkey anti-rabbit IgG (Jackson ImmunoResearch, 711-295-152; 1 μg/mL).

Fluorescence labeling of molecules

To fluorescently label the SNAPf and Halo7 tags conjugated to target proteins expressed in cells, the cells were incubated at 37ºC in complete growth medium containing membrane-permeable fluorescent ligands. For SNAPf, cells were incubated for 60 min with 50 nM SNAP-tetramethylrhodamine (TMR) star (New England BioLabs, S9105S). For Halo7, cells were incubated for 30 min with 20 nM TMR-Halo linker (Promega, G8251), 20 nM SaraFluor650T Halo linker (SF650; Goryo Chemical, A308-02), or 100 nM JF646 Halo linker (Promega, GA1120). Labeling efficiencies were estimated to be 66 ± 1.8% for SNAPf, 88 ± 2.2% for Halo7, and 76 ± 2.3% for mGFP, using the monomer reference proteins mGFP-CD47-SNAPf and mGFP-CD47-Halo7 (SNAPf and Halo7 tags located in the cytoplasm). Using the 15-amino acid linker ($(\text{Ser-Gly-Gly-Gly-Gly})_3$) employed here, the labeling efficiencies were generally independent of the host target proteins (*20*, *143*). Unless otherwise noted, the Halo7 and SNAPf tags were labeled with SaraFluor650 Halo-ligand and SNAP-TMR star, respectively.

Single fluorescent-molecule imaging and analysis

All single-molecule observations were performed at 37°C, using microscopes enclosed in home-built environmental chambers. Fluorescently labeled molecules in cells were excited by total internal reflection (TIR) illumination for those on the basal PM or by oblique-angle illumination for those on the apical PM, using an objective-lens-type TIR illumination system constructed on Olympus IX83 and Nikon TiE inverted microscopes equipped with 100x/1.49 NA objective lenses (UApoN100XTIRF and ApoTIRF100XC, respectively). Multi-color fluorescence images were separated into the three detection arms of the microscope by a dichroic mirror (Chroma, ZT561rdc-UF3 and ZT640rdc-UF3). Each detection arm was equipped with band-pass filters (Chroma, ET525/50m, ET600/50m, or ET700/75m), and images were recorded using a two-stage microchannel plate intensifier (Hamamatsu Photonics, C9016-02MLP24) lens-coupled to a scientific complementary metal oxide semiconductor (sCMOS) camera (ORCA-Flash ver4.0 V2 plus; Hamamatsu Photonics, C11440-22CU). The final magnifications were 133x (50 nm/pixel) and 100x (65 nm/pixel). All instruments in the microscope station were controlled by in-house LabVIEW-based software.

For single molecule observations with oblique illumination, the incident excitation laser powers at the sample plane were 0.29 $\mu W/\mu m^2$ for 488 nm (mGFP; Coherent, OBIS 488-100 LS), 0.21 $\mu W/\mu m^2$ for 561 nm (TMR; Coherent, OBIS 561-100 LS), and 0.37 $\mu W/\mu m^2$ for 642 nm (SF650 and JF646; Omicron, LuuXPlus 640-140). For TIR illumination, very shallow angles and twice the incident laser powers were used. Under these conditions, the photobleaching lifetimes for fluorophores on the apical PM with oblique illumination (and the basal PM with TIR illumination) were: mGFP, 0.61 ± 0.02 s (0.74 ± 0.02 s); SNAPf-TMR, 3.65 ± 0.15 s (3.68 ± 0.18 s); Halo7-TMR, 8.87 ± 0.47 s (9.50 ± 0.48 s); Halo7-SaraFluor650, 12.69 ± 0.57 s (14.38 ± 0.53 s); and Halo7-JF646, 11.12 ± 0.62 s (12.01 ± 0.54 s). For GEM cluster observations, the laser powers were reduced by a factor of three compared to those used for single-molecule observations.

Individual fluorescent spots were identified and tracked using an in-house computer program, as described previously (*23*, *114*, *115*, *144*). Multi-color image superimposition was performed as described previously (*145*).

For visualization of GEM, fluorescently labeled talin or zyxin was imaged at single-molecule sensitivity (with laser power reduced by a factor of three compared to standard single-molecule observations) at 30 frames/s. Images were rolling-averaged over 30 frames and background-subtracted using a rolling-ball algorithm with a 30-pixel (1.5-µm) radius. Talin and zyxin clusters were identified using an in-house computer program originally developed for detecting single-molecule Gaussian spots (*23*, *114*, *115*, *144*). Copy number per cluster (GEM size) was determined by comparing cluster spot intensities with those of a monomer control molecule, the tag protein fused to a C-terminal K-Ras CAAX motif (farnesylated tag proteins).

Transient-immobilization events, termed Temporary Arrest of LateraL diffusion (TALL(*119*, *146*)), were detected in single-molecule trajectories using a previously described algorithm (*147*), as modified in subsequent studies (*119*, *146*). The detection circle radius was set to 200 nm, and the residency-time threshold was set to 5 frames (0.167 s). Cluster lifetimes and molecular dwell lifetimes at GEM were not corrected for photobleaching, because the photobleaching lifetimes of the probes were much longer than the observed cluster lifetimes and molecular dwell lifetimes under the employed imaging conditions.

#### Colocalization duration analysis

Colocalization of two fluorescent molecules was defined as an event in which two molecules are localized within 200 nm of each other, as described previously (*145*, *148*). The duration of each colocalization event was measured, and the distribution of colocalization durations was obtained from all detected events. We found that the distributions were best fitted by the sum of two exponentials, based on both the Akaike and Bayesian information criteria. The shorter decay component represents the incidental colocalization lifetime$\tau$ ($\tau_1$; colocalization without molecular binding), whereas the longer decay component represents the molecular binding duration ($\tau_2$) (*6*, *148*).

Colocalization index analysis

To quantitatively measure the colocalization of single molecules with GEM (talin and zyxin clusters), whose apparent sizes were close to the diffraction limit and therefore indistinguishable from single-molecule spots, we employed a "colocalization index" based on a pair cross-correlation analysis of two-color single-molecule movies (e.g., green and red) (*21*, *148*, *149*), as illustrated in fig. S1H. Briefly, (1) a region of interest (ROI) was selected, and the distances between all pairs of green and red fluorescent spots in the ROI were measured in each video frame; this procedure was repeated for all frames. (2) The distribution of pair counts, $R(r)$, was plotted as a function of the distance, $r$. The normalization factor, $N(r)$, was computed using the fast Fourier transform and the inverse fast Fourier transform (*21*, *149*). (3) The cross-correlation function was calculated as $C(r) = R(r)/N(r)$. (4) The colocalization index was defined as the ratio of the $C(r)$ between 0 and 50 nm to the $C(r)$ between 450 and 500 nm. In the absence of significant colocalization, the index equals 1, whereas increasing colocalization yields higher values. Control data without correlations between green and red spots were generated by shifting each track in the image by 1 μm in random directions.

PALM observation and analysis

Cells expressing the proteins fused to mEos3.2 were chemically fixed prior to PALM observations. Briefly, cells were initially fixed with 0.3% (vol/vol) glutaraldehyde in cytoskeleton buffer (150 mM NaCl, 5 mM EGTA, 5 mM glucose and 5 mM $MgCl_2$, buffered with 10 mM MES, pH 6.1) containing 0.25% (v/v) Triton X-100 for 1 min, followed by post-fixation with 2% (vol/vol) glutaraldehyde in the same solution for 15 min (*150*, *151*). Cells were then incubated with freshly prepared 0.1% (wt/vol) sodium borohydride for 7 min, to quench unreacted aldehydes. After replacing the cytoskeleton buffer with T-HBSS, the fixed cells were imaged at 37°C using a home-built PALM station (*114*, *115*), built on an Olympus IX83 microscope.

Samples were illuminated in the TIR or oblique-angle mode, using a 405-nm activation laser (Cobolt, 06-MLD 405-100; increased from 1.5 to 78 nW/μm$^2$ at the sample plane following the Fermi photoactivation scheme [$t_F = 162$, $T = 16.2$ s]) (*152*), a 561-nm excitation laser (MPB Communications, Fiber 560-20000; 12.4 μW/μm$^2$), and an additional weak 488-nm laser to suppress the long-lived dark state (*153*) (Coherent, Sapphire 488-300 CW; 0.164 μW/μm$^2$). PALM data were acquired at 60 Hz for 14,560 frames with an image size of 512 x 512 pixels, using a high-speed camera system (Photron, mini AX200-II) as described previously (*114*, *115*). The final magnification was 250x (80 nm/pixel).

Spot detection was performed using the ThunderSTORM plugin in ImageJ (*154*) with Wavelet filtering (B-spline order = 3; scale = 2.0) and the local maximum method (peak intensity threshold = 30-50; connectivity = 8-neighborhood). Post-processing included removing duplicates (distance threshold = uncertainty) and drift correction (cross-correlation with 5 bins). Processed datasets were imported into the SR-Tesseler (*40*) for cluster analysis and image reconstruction. ROIs were selected in apical PM regions with good focus, and for the basal PM analyses, ROIs were selected from the regions without obvious IACs. Blinking and multi-ON frame detections were corrected using the detection cleaner function (photon/background size = 0.3; fixed maximal dark time = 200; intensity-to-photon ratio = 0.039; background value = 0.3) (*153*). Cluster contour detection was then performed using a Voronoï polygon density factor of 3.5 applied across the entire ROI.

Optogenetics

For the partial depletion of $PI(4,5)P_2$, CRY2 fused to mScarlet and 5-phosphatase-OCRL (mScarlet-CRY2-5PT) and CIBN fused to farnesylated mGFP (CIBN-mGFP-far; farnesylation for the localization on the PM cytoplasmic surface) were expressed in talin-rescued dKO MEFs (used to determine the copy number of talin per GEM; Fig. 2D). In the dark, CRY2 does not bind to CIBN. Upon blue light irradiation (488-nm laser epi-illumination of the entire cell; train of 200-ms pulses at 0.5 Hz for 30 s; 0.64 nW/$\mu m^2$; at the sample plane), light-activated CRY2 bound to CIBN on the PM, recruiting mScarletI-CRY2-5PT to the PM, and inducing $PI(4,5)P_2$ hydrolysis (fig. S4D). The $PI(4,5)P_2$ levels in the PM were monitored using a fluorescent probe consisting of two tandem PH domains of PLCδ fused to SNAPf (SNAPf-2xPH). After the blue light was turned off, CRY2 was slowly deactivated, leading to the dissociation of mScarlet-CRY2-5PT from the PM (fig. S4D). The phosphatase-dead D523G mutant of 5PT fused to mScarlet-CRY2-5PT served as a control.

To quantify $PI(4,5)P_2$ levels at the PM, cell boundaries were determined from the SNAPf-2xPH images using ilastik (*155*). The outer and inner contours of the cell were then set at ± 1 µm from the cell boundary to define the PM region. Mean pixel intensities in the PM region ($F_{PM}$) and in the cytoplasm ($F_{cyto}$) were measured (fig. S4F). $PI(4,5)P_2$ levels decreased rapidly after blue light irradiation, and returned to baseline within 10 min after the 30-s irradiation was terminated. Single-molecule observations of talin clusters (GEMs) were performed before, immediately after, and 15 min after the blue light irradiation (Before, After, and Recovery in fig. S4H).

For artificial droplet formation (*68*, *69*), cDNAs of optoFAK and optoLyn were generated by combining the sequences encoding the Src myristoylation signal peptide, the N-terminal IDR of FUS, fluorescent tag proteins (mGFP, Halo7, or SNAPf), CRY2, and either FAK or Lyn (with mutations in the N-terminal myristoyl/palmitoyl modification signals), and expressed in the cell (fig. S13A). In the dark, CRY2 does not form clusters. Upon blue light irradiation, light-activated CRY2 formed clusters via homotypic interactions. The blue light intensity and pulse train were identical to those used for $PI(4,5)P_2$ depletion. Irradiation started 2 min before PDGF stimulation and continued during recording.

To detect FAK activation within droplets, the binder-tag system was used (*67*). The SsrA peptide is inserted into the central region of FAK, where it becomes exposed in the active open conformation, allowing specific binding by the SNAPf-tagged SspB binder peptide (fig. S13F).

Confocal microscopy and FRAP

For live-cell confocal fluorescence imaging at 37ºC, we used a home-built spinning-disk confocal microscope based on an Olympus IX83 enclosed in a home-built environmental chamber, equipped with a 60x or 100x objective (Olympus, PLXApo60XO or UApoN100XTIR), a confocal unit (Yokogawa CSU-W1), and an sCMOS camera (Hamamatsu Photonics, ORCA-Flash 4.0 V2 plus).

For in-cell FRAP, a Nikon A1R confocal microscope equipped with a stage-top incubator (Tokai-Hit, STXG-TIZWX-SET) was used at 37ºC with a recording rate of 0.5 Hz. The photobleached ROI was circular (≈1 μm diameter), and photobleaching was performed by illuminating the area with a 488- or 561-nm laser at 30% of the full laser power for 1 s.

For *in vitro* FRAP, purified proteins were diluted with observation buffer (150 mM NaCl and 0.5 mM tris(2-carboxyethyl)phosphine) [TCEP], buffered with 25 mM Tris–HCl, pH 7.4). After the addition of a final concentration of 1% PEG8K (Sigma-Aldrich, P5413), the solution was placed on the glass-bottom slide (Ibidi, 81507) and observed at 37ºC and 1 Hz using a Nikon spinning-disk confocal microscope, consisting of a Nikon TiE inverted microscope with a PlanApo 60x/1.40 objective lens (Nikon), a confocal unit (Yokogawa, CSU-X), an Andor Revolution SD System, an EM-CCD camera (Andor, iXon 888-U3ultra), and a stage top incubator (OkoLab). The bleached ROI was circular (≈1.5 μm diameter), and photobleaching was performed with a 488-nm laser (3.9 mW) for 0.5 s. Fluorescence recovery curves were background-corrected and normalized using ImageJ and analyzed as described previously (*156*).

Cell volume measurement

Cells were transfected with a plasmid encoding SNAPf alone (without fusion proteins) and plated on FN-coated glass-bottom dishes. After 48 h, the expressed SNAPf proteins were labeled with SNAP-TMR Star, and the medium was replaced with 1:1,000 diluted NucSpot Live 650 (Biotium, 40082) in T-HBSS for nuclear staining. Without washing, whole-cell 3D images were acquired by Z-stack scanning (250-nm step size) using the home-built Olympus spinning-disk confocal microscope.

Images were binarized in ImageJ, using Otsu's algorithm (*157*). Whole-cell and nuclear volumes were determined from the binarized images of SNAPf and NucSpot Live 650, respectively, by counting the total voxels. Cytoplasmic volume was calculated as the whole-cell volume minus the nucleus volume.

Measurements of intracellular $Ca^{2+}$ concentrations

Cells were incubated with 20 μM CalBryte 590 (AAT Bioquest, 20700) in T-HBSS at 37ºC for 30 min, followed by three washes with T-HBSS. Cells were imaged at 37ºC using a home-built objective-lens-type TIRF microscope (Nikon TiE-based) equipped with a Plan Fluor 20x/0.50 objective lens (Nikon), with epi-illumination using a 561-nm laser (0.22 nW/$\mu m^2$ at the sample plane). Recordings were performed at 1 or 2 Hz. All observations were completed within 1 h after labeling.

Purification of protein probes for cholesterol and sphingomyelin

For the production of cholesterol and sphingomyelin probes, BL21 (DE3) *E. coli* competent cells (New England BioLabs, C2527) were transformed by heat shock with expression plasmids encoding Halo7-tag fused to D4H (*117*, *118*) and Halo7-tag fused to OlyA (E69A) (*29*). Transformed cells were cultured at 37ºC in Luria-Bertani (LB) medium containing 50 µg/mL kanamycin with shaking until the optical density at 600 nm reached 0.4-0.6, and protein expression was induced with 0.6 mM isopropyl-β-D-thiogalactopyranoside (IPTG; Amresco, MFCD00063273) at 24ºC for 12 h. Cells were harvested by centrifugation at 7,000 rpm for 3 min, resuspended in lysis buffer (1% Triton X-100 and 300 mM NaCl, buffered with 20 mM Tris-HCl, pH 8.0, and supplemented with protease inhibitor cocktail; Nacalai Tesque), and lysed by agitation on ice for 20 min. Lysates were centrifuged at 12,000 rpm for 10 min, and the supernatants were filtered through a 0.45-µm Millex filter. Proteins were purified by Ni-NTA agarose chromatography (1 mL bed volume; Sangon Biotech, C600332-0001). Bound proteins were eluted sequentially with 10, 20, and 30 mM imidazole (10, 10, and 5 mL, respectively) in 150 mM NaCl buffered with 20 mM Tris-HCl (pH 8.0), followed by a final elution with 100 mM imidazole in 250 mM NaCl, 20 mM Tris-HCl, pH 8.0 (10 ml).

Eluted fractions were desalted and concentrated with an Amicon Ultra concentrator (10 kDa cutoff; Millipore, UFC501008) into 150 mM NaCl buffered with 20 mM 4-(2-hydroxyethyl)-1-piperazine ethanesulfonic acid (HEPES; pH 7.4). Protein concentrations were determined by the absorbance at 280 nm, using extinction coefficients of 1.02 and 0.977 x $10^5$ (M/cm) for Halo7-D4H and Halo7-OlyA(E69A), respectively (calculated from amino acid composition data) (*158*).

For fluorescence labeling, purified proteins were incubated with 1.5 molar equivalents of SeTau647-Halo ligand (*119*) at 37ºC for 3 h. Unreacted ligand was removed by Ni-NTA affinity column chromatography and reconcentration. Purities were 71% (Halo7-D4H) and 90% (Halo7-OlyA[E69A]), as determined by SDS-PAGE with Coomassie Brilliant Blue R-250 (CBBR-250; Fluka, 27816) staining. Aliquots (30 µL) of the labeled protein solutions were snap-frozen in liquid nitrogen and stored at -80ºC.

Purification of zyxin and VASP

For the preparation of zyxin (WT and mutants) and VASP, proteins were expressed in Expi293F cells (Thermo Fisher Scientific, A14635) for three days as fusion constructs containing a 6xHis tag, maltose binding protein (MBP), Tobacco Etch Virus (TEV) protease digestion sequence, and mCherry-zyxin or EGFP-VASP. Cells were harvested and lysed by freeze-thaw cycles in buffer A (200 mM NaCl, 1 mM EDTA, and 0.5 mM TCEP, buffered with 25 mM Tris-HCl, pH 7.4, and supplemented with protease inhibitor cocktail; Roche, 11873580001). Lysates were centrifuged at 21,000 x g at 4ºC for 1 h, and the proteins were purified in two steps. First, lysates were loaded onto an MBP-binding dextrin Sepharose column (MPBTrap; GE Healthcare, 28-9187-78) twice at 0.5 mL/min and eluted with 10 mM maltose in buffer A at RT. Eluates were digested with His-tagged TEV protease (1:10 enzyme:protein ratio by weight) at 16ºC for 18 h. Second, digests were processed by Ni-NTA chromatography (HisPur™ Ni-NTA Resin; Thermo Scientific, 88222) to separate mCherry-zyxin or EGFP-VASP from 6xHis-MBP and His-tagged TEV protease. For this step, NaCl and imidazole were added to the digested solutions to final concentrations of 400 and 25 mM, respectively, and the solution was passed twice over a pre-equilibrated Ni-NTA Sepharose slurry (5 mL bed volume) in a Poly-Prep Chromatography Column (Bio-Rad, 731-1550), equilibrated with a buffer solution (300 mM NaCl, 25 mM imidazole, and 0.5 mM TCEP, buffered with 25 mM Tris-HCl, pH 7.4). The flow-through fractions were concentrated using a Pierce Protein Concentrator (30 kDa MWCO, PES-membrane; Thermo Scientific, 88502) to ≈150 µL and dialyzed into buffer B (150 mM NaCl and 0.5 mM TCEP, buffered with 25 mM Tris-HCl, pH 7.4) at 4ºC overnight using micro-dialyzer units (10 kDa MWCO; Slide-A-Lyzer MINI; Thermo Fisher Scientific, 69576).

Protein concentrations were determined by the absorbance at 280 nm, using extinction coefficients of 6.11, 6.15, 5.30, 5.18, and 4.84 x $10^4$ (M/cm) for GFP-VASP, mCherry-zyxin (WT), mCherry-zyxin (ΔDR2), mCherry-zyxin (ΔIDR1-2, ), and mCherry-zyxin (IDR2), respectively (calculated from amino acid composition) (*158*). Purities were 84, 72, 69, 89, and 71%, respectively, as determined by SDS-PAGE stained with InstantBlue Coomassie Protein Stain (Abcam, ab119211). Aliquots (10 µL) were snap-frozen in liquid nitrogen and stored at -80ºC.

*In vitro* phase separation assay

Purified zyxin and VASP were diluted in buffer B to the desired concentrations. After the addition of 1% PEG8K, mixtures were transferred to glass-bottom slides (Ibidi, 81507) for imaging. All reactions and imaging were performed at 37ºC.

*In vitro* cell growth assay

For growth assays, 1 x$10^4$ MEFs or 5 x$10^4$ transformed MEFs, RKO, and MDA-MB231 cells were seeded per well in six-well plates in triplicate. Cells were counted every 2 days using an automated cell counter (Invitrogen, Countess), and proliferation curves were plotted based on cell numbers per well.

Complement-dependent cytotoxicity assay

Transformed zyxin-KO MEFs were plated on a polystyrene-bottom 96-well plate (Corning, 3603) at 2.5 x $10^3$ cells/well and cultured overnight. The culture medium was replaced with 50 µL T-HBSS containing 20 µg/mL rabbit anti-CHO polyclonal antibodies (ProteinTech, 27803-1-AP), 1 µM Blue DCS1 (AAT Bioquest, 17548; membrane-impermeable nuclear staining dye for dead cells), and 1:20 diluted NucRed Live 647 (Invitrogen, R37106; membrane-permeable nuclear staining dye for total cells). After an incubation at 37ºC for 30 min, the cells were washed and imaged by epi-fluorescence microscopy (4x objective lens). The culture medium was then replaced with 50 µL T-HBSS containing 1 µM Blue DCS1, 1:20 diluted NucRed Live 647, and varying concentrations of standardized human serum (Sigma-Aldrich, C9473). After an incubation at 37ºC for 2 h, the cells were washed and re-imaged. Floating dead cells were removed during washing. Acquired images were binarized by ilastik, and the NucRed Live 647-positive and Blue DCS1-negative cells were counted as live cells. Total and live cell numbers were calculated from the first and second images, respectively. Negative controls used heat-inactivated standard serum (56ºC for 30 min).

Cell growth assay after complement stimulation

Transformed zyxin-KO MEFs were plated on a polystyrene-bottom 384-well plate (Corning, 4514) at 5 x $10^2$ cells/well and cultured for 24 h. The culture medium was replaced with complete growth medium containing 20 µg/mL rabbit anti-CHO polyclonal antibodies and 10% C9-depleted human serum (Calbiochem, 234409), followed by an incubation for 12 h. Since C9-depleted serum cannot form a complete membrane attack complex (MAC), this system specifically assesses CD59 activation by the C5-8 complex (MAC precursor). After the incubation, the cells were fixed with 4% PFA, and nuclei were stained with Blue DCS1. Cell numbers were counted from fluorescence images.

Single-molecule imaging of MAC precursor

For fluorescence labeling of complement C8, human complement C8 (Sigma-Aldrich, C3535) was labeled with CF640R-succinimidyl ester (Biotium, 92108) according to the manufacturer's instructions. The labeled C8 protein was purified with an Amicon Ultra spin filter (10 kDa cutoff; Millipore). The dye/protein ratio was calculated to be 3.4.

For single-molecule imaging of MAC precursor, transformed zyxin-KO MEFs expressing mGFP-zyxin, cultured on glass-bottom dishes, were incubated in T-HBSS containing 20 µg/mL rabbit anti-CHO polyclonal antibodies for 10 min at 37ºC. Cells were washed and further incubated with T-HBSS containing 10% C8-depleted human serum (Sigma-Aldrich, C1538), 1% BSA, and 100 nM C8-CF640R for 10 min at 37ºC. After washing, the cells were imaged, and the imaging was completed within 10 min.

Mice

Female BALB/cAJcl-*Foxn1$^{nu}$* (nude) mice (6–8 weeks old) were purchased from CLEA Japan, Inc. (BALB/cAJcl-*nu/nu*). All mice were maintained under specific pathogen-free conditions. All animal procedures were approved by the Animal Care and Use Committee at Okinawa Institute of Science and Technology Graduate University (OIST; approval no. 2022-362-2).

Tumor xenograft assays

Cells were trypsinized and resuspended in DMEM supplemented with 30% FBS, and then mixed with VitroGel (TheWell Bioscience, VHM01) at a 1:2 ratio (vol/vol). A total of 100 µL of the cell-gel mixture, containing either 5 x $10^6$ transformed MEFs or 2 x $10^6$ RKO cells, was injected subcutaneously into the right flank of 6- to 8-week-old female nude mice (BALB/cAJcl-*nu/nu*). Tumor growth was monitored every ≈3 days starting ≈7 days after implantation. Tumor volume ($V$) was estimated using the formula, $V = 0.5$ x length x width$^2$,(*159*) where length and width represent the longest and shortest dimensions, respectively, measured with calipers. Measurements were performed by multiple investigators blinded to group identity. At ≈30 days after implantation, mice were euthanized in a $CO_2$ chamber, and the tumors were excised, weighed, and photographed. Small tumor pieces (≈2 x 2 x 2 mm) were collected for downstream analysis, including western blotting. Tumor implantation experiments were independently performed twice, using 6 and 4 mice per group in the two replicates.

Coarse-grained models for Zyxin (IDR2)

We modeled the zyxin IDR2 using a residue-resolution coarse-grained (CG) representation, where each amino acid is represented by a single interaction site. Simulations were performed using the Mpipi model (*52*). The total potential energy is written as:

$$V_{\mathrm{MPIPI}} = \sum_{(i,j)\in\mathrm{bonds}} E_{\mathrm{bond}}(b_{ij}) + \sum_{(i,j)\in\mathrm{nonbonded\ particles}} E_{\mathrm{WF}}(r_{ij}) + \sum_{(i,j)\in\mathrm{charged\ particles}} E_{\mathrm{ele}}(r_{ij}) \ .$$

Here, $E_{bond}(b_{ij})$ constrains chain connectivity through a harmonic bonded interaction with bond length $(b_{ij})$. Nonbonded contacts between residue pairs are treated using the Wang-Frenkel potential,

$$E_{\mathrm{WF}}(r_{ij}) = \varepsilon_{ij}\ \alpha_{ij} \left[\left(\frac{\sigma_{ij}}{r_{ij}}\right)^{2\mu_{ij}} - 1\right]\left[\left(\frac{R_{ij}}{r_{ij}}\right)^{2\mu_{ij}} - 1\right]^{2\nu_{ij}} \ ,$$

where $\varepsilon_{ij}$, $\sigma_{ij}$, $R_{ij}(= 3\sigma_{ij})$, $\mu_{ij}$, $\nu_{ij}$, and $\alpha_{ij}$ depend on the types of residues $i$ and $j$.

Electrostatic interactions between charged residues are modeled using a Debye-Hückel formulation:

$$E_{\mathrm{ele}}(r_{ij}) = \frac{q_i q_j}{4\pi\epsilon_0\epsilon_r(T,C)} \frac{\exp\left(-r_{ij}/\lambda_D\right)}{r_{ij}} \ ,$$

where $q_i$ and $q_j$ are particle charges, $\lambda_D$ is the Debye screening length, and $\epsilon_r(T,C)$ denotes the dielectric constant of the solution as a function of temperature $(T)$ and ionic strength $(C)$. The temperature and salt dependencies follow:

$$\epsilon_r(T,C) = e(T)a(C) \ ,$$

where $e(T) = 249.4 - 0.788\,T + 7.20 \times 10^{-4}T^2$, and $a(C) = 1 - 0.2551C + 5.151 \times 10^{-2}C^2 - 6.889 \times 10^{-3}C^3$.

Finally, the Debye length is computed as:

$$\lambda_D = \sqrt{\frac{\epsilon_0\epsilon_r(T,C)k_B T}{2N_A e_c^2 I}} \ ,$$

where $N_A$ is Avogadro's number, $e_c$ the elementary charge, and $I$ the ionic strength.

Molecular dynamics simulations of Zyxin IDR2

Zyxin IDR2 (Fig. 3D) was used as the model system in molecular dynamics simulations. CG topologies and initial coordinates were generated using GENESIS-cg-tool (*160*). To characterize the temperature-dependent behavior, we first simulated a single chain over a temperature range of 210-440 K in 10 K increments, with each trajectory run for $10^7$ MD steps. These simulations were used to identify the approximate temperature window associated with phase transitions (fig. S8, A and B). Structures sampled from these single-chain trajectories were subsequently used as templates for constructing multi-chain systems.

To directly probe the phase behavior, we built condensate-scale systems composed of 200 zyxin IDR2 chains. Initial configurations were generated by placing structures sampled from the single-chain simulations onto a 2 x 4 x 25 spatial grid. The simulation cell was then gradually reduced in size using the box-shrink protocol in GENESIS ATDYN until a compact size of 250 x 250 x 400 Å$^3$ was achieved. After equilibration, the periodic boundary conditions were elongated along the z-axis to 250 x 250 x 2000 Å$^3$, and these structures served as starting points for production simulations. Each condensate system was simulated for $10^7$ steps at 210, 230, 250, 270, and 290K.

All simulations employed the GENESIS MD software: ATDYN (v2.1.5) (*161*) for single-chain simulations and multi-chain box-shrink procedures, and CGDYN (v2.1.0-beta) (*162*) for condensate production runs. A time step of 10 fs was used. Nonbonded electrostatics were calculated with a 35-Å cutoff, and the time integration used a Langevin thermostat with a friction coefficient of 0.01 ps$^{-1}$.

Molecular dynamics simulation data analysis

To identify the dominant interactions stabilizing zyxin IDR2 condensates, we performed contact analysis. For each snapshot, a smoothed contact score was computed for every pair of residues with indices $i$ and $j$ across different chains:

$$n(i,j) = \frac{1}{n_{chain}} \sum_{\text{chain } I} \sum_{\text{chain } J \neq I} \sum_{\text{res}_i \in I} \sum_{\text{res}_j \in J} n_c(\text{res}_i, \text{res}_j) \quad ,$$

where the pairwise term $n_c$ was defined using a sigmoidal switching function:

$$n_c = \frac{1}{1 + e^{(r_{ij} - r_0)/r_w}} \quad .$$

Here, $r_{ij}$ is the distance between residues res$_i$ and res$_i$, with $r_0 = 10$ Å as the characteristic length and $r_w = 1$ Å controlling the smoothing width. Time-averaged values of $n(i,j)$ were used to construct the two-dimensional contact probability map (Fig. 3F, left). The total number of contacts made by the $i$-th residue was then obtained by summing over all partners:

$$N(i) = \sum_j n(i,j) \quad .$$

To quantify interaction preferences between residue types, we calculated type-type normalized contact scores (Fig. 3F, right):

$$p(a,b) = \frac{\sum_{i \mid \text{type}_i = a} \sum_{j \mid \text{type}_j = b} n(i,j)}{o_a \, o_b} \quad ,$$

where $o_a$ and $o_b$ denote the total occurrences of residue types $a$ and $b$ in zyxin IDR2, respectively.

Quantification and statistical analysis

Superimposition of dual-color image sequences and single-molecule tracking were performed using in-house C++-based computer programs, as previously described (*144*, *145*). Conventional TIRF images were analyzed using Fiji (imageJ) (*163*), MatLab 2019a (MathWorks), and ilastik (*155*).

Curve fitting was performed using Origin 2017 (OriginLab). Statistical analyses were performed using OriginPro 2017, Rstudio, and Microsoft Excel. Statistical significance was defined as $P<0.05$. Other image and data analyses were performed using imageJ (Fiji) and MatLab 2019a. Figures and videos were edited using Photoshop CC 2021 (Adobe) and Illustrator CC 2021 (Adobe).

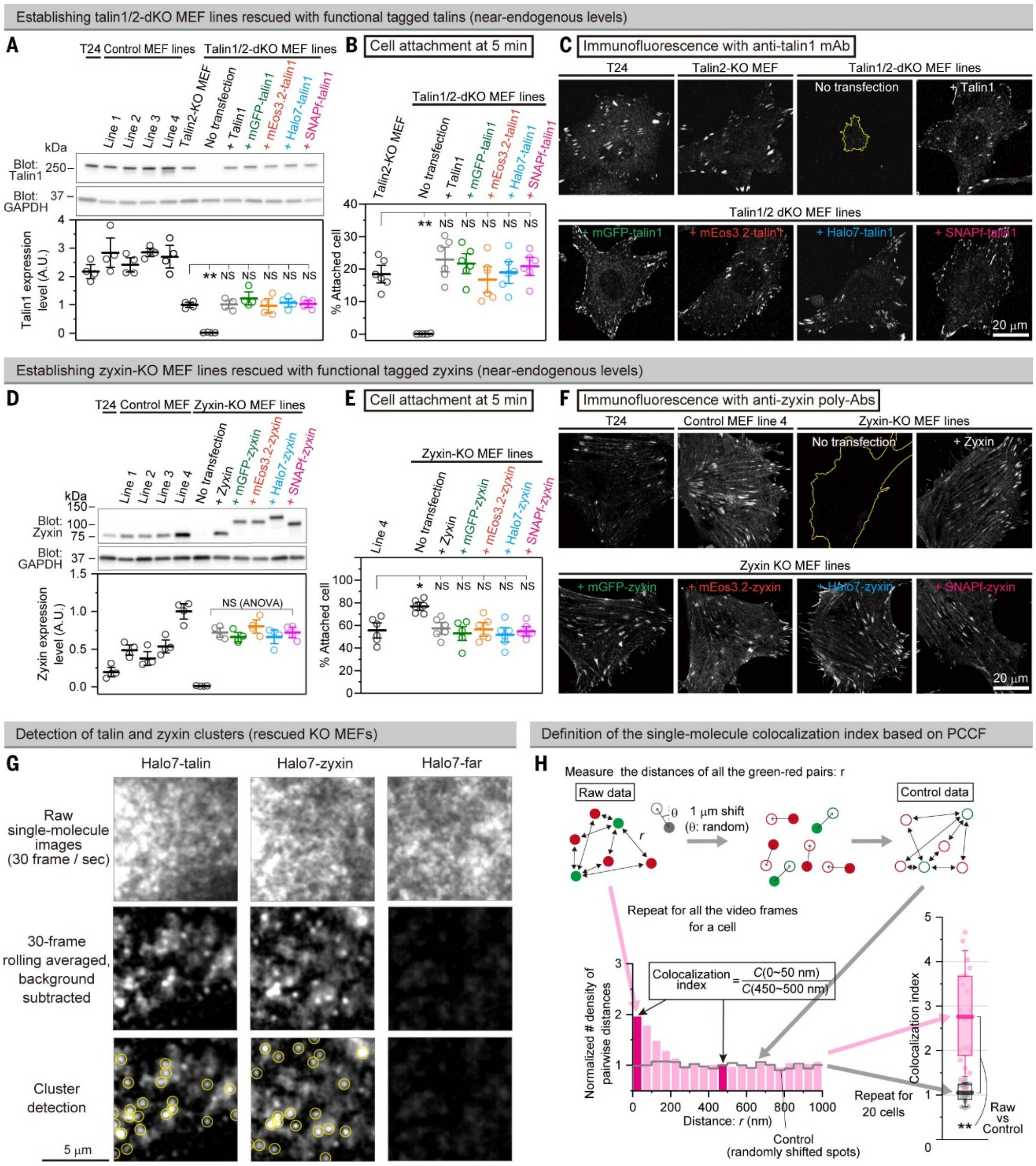


**Fig. S1. Establishment of talin- and zyxin-rescued MEF lines, detection of talin/zyxin clusters, and calculation of single-molecule colocalization index.**

(**A to C**) Establishing talin1/2-dKO MEF lines rescued with various tagged talin1 constructs expressed at levels comparable to endogenous talin1 in talin2-KO MEFs (parental cell line for generating talin1/2-dKO MEFs).

(A) Western blot quantification of talin1/tagged talin1 expression (mean ± SEM, $n$ = 4 replicates). Since talin1 is the main isoform in most tissues (*164*), we employed talin1 rather than talin2. Unless otherwise specified, "talin" refers to talin1 throughout. Because the talin-dKO MEF lines were generated from talin2-KO MEFs (as the parental cell line) (*19*), the parental talin2-KO line was used as a control; control MEF lines 1-4 are independent immortalized MEF controls.

(B) Tagged-talin constructs are functional in fibronectin-dependent attachment (5 min after plating; mean ± SEM, $n$ = 6 dishes), in agreement with the previous report (*19*).

(C) Tagged-talin constructs incorporate into IACs. Confocal immunofluorescence microscopy of rescued dKO MEFs using an anti-talin1 monoclonal antibody (confocal section near the basal PM). The contour of a non-rescued cell is shown in yellow.

(**D to F**) Establishing zyxin-KO MEF lines rescued with various tagged zyxin constructs, expressed at levels comparable to the mean endogenous zyxin level in four control MEF lines. The same validation workflow as in (A to C) was applied.

(D) Because the zyxin-KO MEF line was derived from a zyxin-KO mouse (*22*), no parental cell line is available; control MEF lines 1-4 were therefore used as references for expression matching.

(E) Tagged-zyxin constructs are functional, as shown by the *lower* ability of the zyxin-KO MEFs rescued with tagged zyxin constructs to attach to fibronectin-coated plastic dishes (mean ± SEM, $n$ = 6 dishes), in agreement with the previous report (*22*).

(F) Tagged-zyxin constructs incorporate into IACs (basal confocal section). The contour of a non-rescued cell is shown in yellow (these cells spread more than rescued MEFs).

(**G**) Snapshot from movie S1. The panel for the cluster detection of Halo7-talin is the same as Fig. 1A. Single-molecule visualization of fluorescent talin and zyxin clusters located on the apical PM (left and middle). Clusters were detected after 30-frame rolling averaging (1 s) and background subtraction using an in-house Gaussian-spot detection routine (yellow circles; bottom). The result of Halo7-far [Farnesylated Halo7] is shown as a monomeric control (right column).

(**H**) Method for evaluating the colocalization index based on the pair cross-correlation function (PCCF) (*20*, *21*, *149*), For details, see materials and methods.

Games-Howell's test for (A, B, and E) and one-way ANOVA for (D).

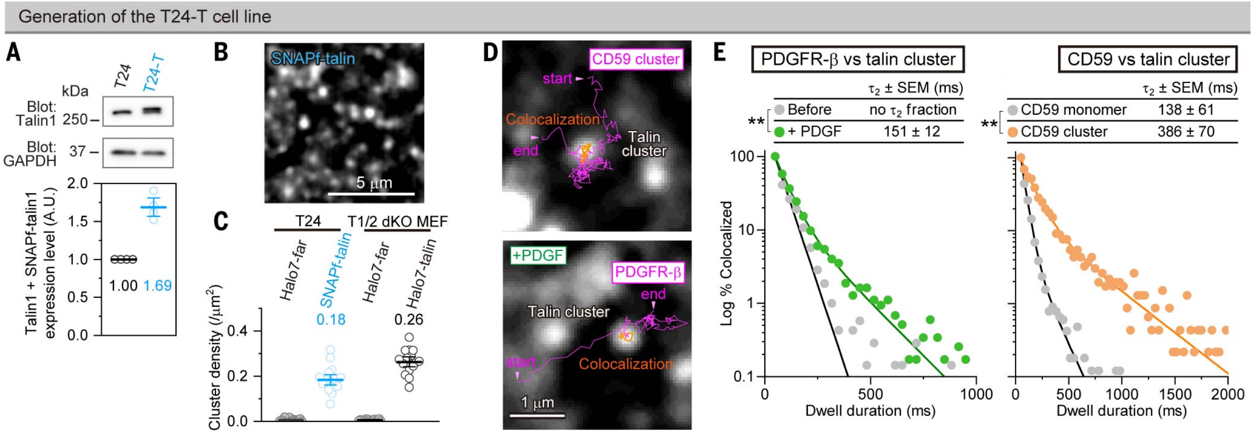


**Fig. S2. Establishment of the T24-T cell line.**

Establishment of the T24-T cell line (T24 stably expressing SNAPf-talin; "T" denotes talin), used for testing the recruitment of FA proteins and signaling molecules to talin/zyxin clusters. T24 cells were selected because single-molecule imaging on the apical PM is facilitated by their flatter apical surfaces. The Halo7 tag was used to label signaling molecules, whereas the SNAPf tag was used for talin. Labeling efficiencies were ≈88% (Halo7) and ≈66% (SNAPf) (see materials and methods).

(**A**) Western blot quantification of total talin1 (endogenous + SNAPf-talin1; bands not resolved and thus quantified together) in T24-T cells (mean ± SEM, n = 4 replicates). Total talin1 is ≈1.7× higher than in parental T24 cells.

(**B**) Typical image of SNAPf-talin located on the apical PM of a T24-T cell.

(**C**) The number density of talin clusters found on the apical PM of T24-T cells (containing SNAPf-talin) is 0.18/μm$^2$, slightly lower than in talin-dKO MEFs rescued with Halo7-talin (0.26/μm$^2$, $n$ = 15 cells).

(**D and E**) In T24-T cells, as found in MEFs, engaged CD59 and PDGFR (trajectories in **L**) are transiently recruited to talin clusters (white puncta in D), confirming the consistency between the two cell types in terms of receptor recruitment (Fig. 1, C and 1D). Their dwell lifetimes are 0.39 ± 0.070 and 0.15 ± 0.012 s, respectively (E), consistent with the findings in MEFs (0.32 ± 0.061 and 0.22 ± 0.035 s, respectively; Fig. 1D). $n$ = 20 cells, $n$ = 3,209, 2,471, 1,485, and 3,261 events for CD59 monomer, CD59 cluster, without PDGF, and PDGF addition, respectively.

Brunner-Munzel's test for (E).

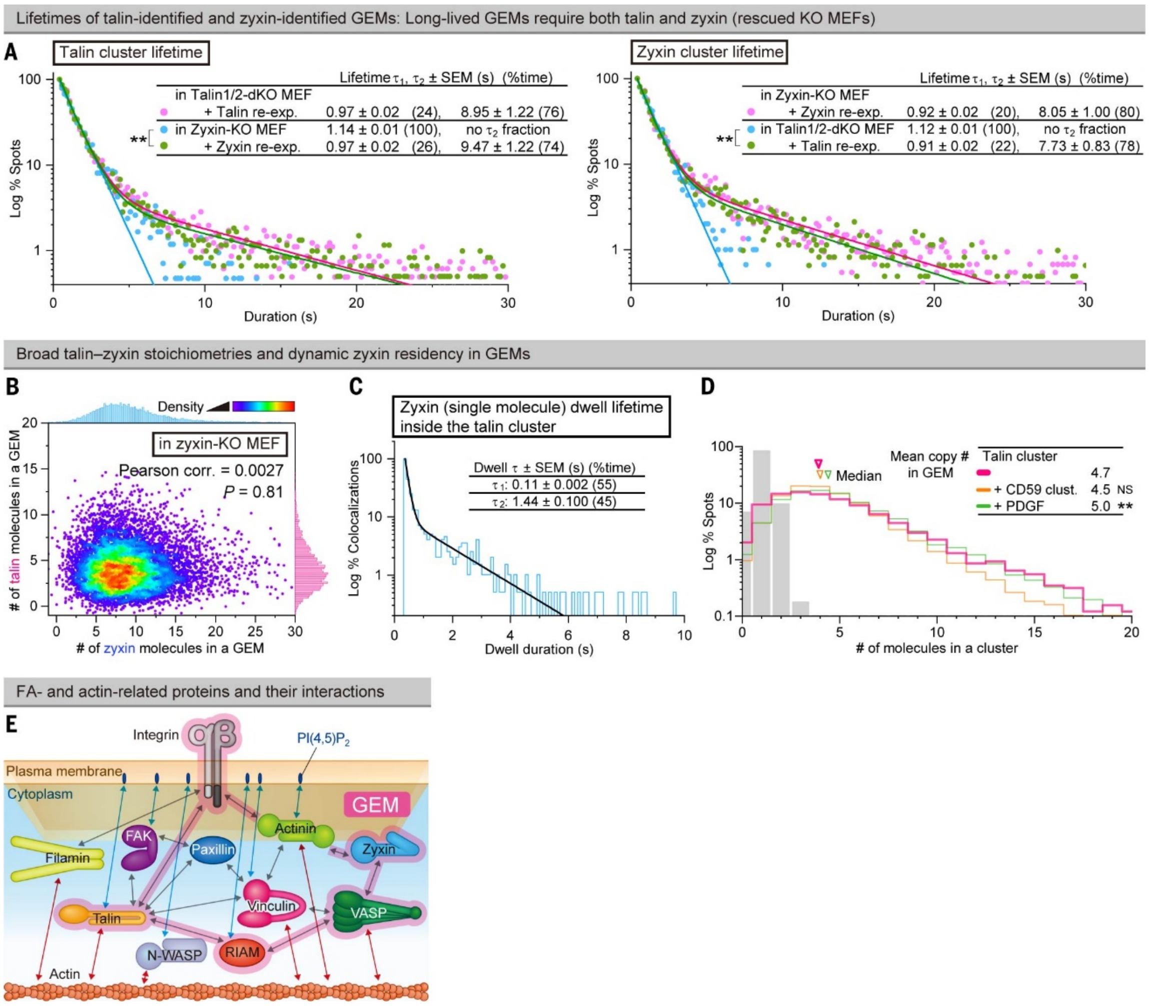


**Fig. S3. GEM lifetimes and molecular stoichiometry, composition, and dynamics.**

(**A**) Two-component lifetime analysis distinguishes talin–zyxin co-clusters from incomplete clusters. The plots show lifetime distributions of Halo7-talin clusters in zyxin-KO MEFs with or without zyxin re-expression at endogenous levels (left) and Halo7-zyxin clusters in talin-dKO MEFs with or without talin re-expression at endogenous levels (right). Distributions were fitted with a sum of two exponentials (providing decay times; SEM = fitting error, $n$ = 10 cells, $n$ = 2,148 ~ 7,273 clusters). In the absence of the partner protein (talin or zyxin), the long-lived component ($\tau_2$) is largely lost, leaving a short-lived component ($\tau_1 \approx 1$ s). In contrast, talin-zyxin co-clusters exhibit a prominent long-lived component ($\tau_2 \approx 7.7$ - 9.5 s).

(**B**) Zyxin-counterpart of Fig. 2E. Broad and uncorrelated talin/zyxin copy numbers in individual GEMs, indicating flexible stoichiometries (zyxin-KO MEFs; $n$ = 15 cells, $n$ = 7,729 clusters). Fig. 2E shows the corresponding analysis performed in rescued talin-dKO MEFs.

(**C**) Zyxin-counterpart of Fig. 2F. The dwell lifetime of zyxin in GEMs is 1.4 s ($n$ = 10 cells, $n$ = 581 events), which was obtained from single-molecule dwell time distributions. For talin, the corresponding dwell lifetime is 2.1 s (Fig. 2F).

(**D**) Distributions of talin copy numbers per GEM after CD59 and PDGF stimulation. The data before stimulation is shown in Fig. 2D.

(**E**) Schematic overview of reported interactions among IAC-associated proteins and their links to actin filaments, actin binding proteins, and $PI(4,5)P_2$.
Brunner-Munzel's test for (A) and Steel-Dwass' test for (D; vs. talin cluster).

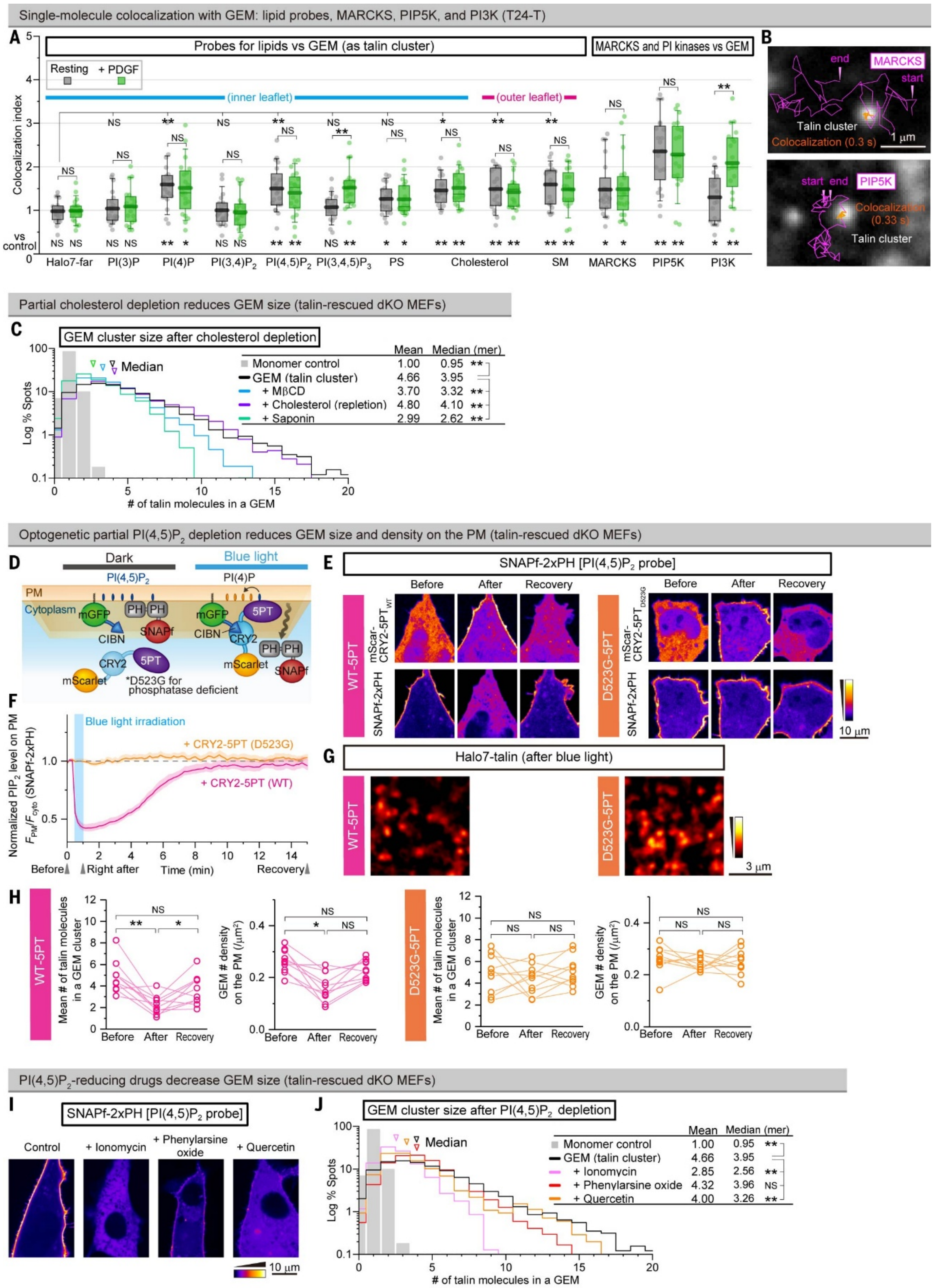


**Fig. S4. PI(4,5)P₂ and raft domains are colocalized with GEM.**

(**A**) Single-molecule colocalization indexes of protein probes for various lipids, MARCKS, and PI kinases (Halo7-tagged) with GEMs (PI3K data are the same as those shown in Fig. 5A; $n$ = 20 cells). Shorter version of this panel is shown in Fig. 2I.

$PI(4,5)P_2$ is enriched at GEMs relative to the bulk PM, consistent with the enrichment of MARCKS, a $PI(4,5)P_2$-binding protein. PIP5K, which produces $PI(4,5)P_2$ from PI(4)P and binds talin,(*165*) and PI(4) are also recruited to GEMs. Their colocalization indexes are not detectably altered by PDGF stimulation. Because PLCγ can hydrolyze $PI(4,5)P_2$ to generate PI(4)P, these localizations are consistent with local $PI(4,5)P_2$ turnover at GEMs.

$PI(3,4,5)P_3$ was detected at GEMs only after PDGF stimulation, and PI3K showed increased colocalization with GEMs only after PDGF stimulation, consistent with PDGF-dependent $PI(3,4,5)P_3$ production from $PI(4,5)P_2$ at GEMs. We did not detect PI(3)P or $PI(3,4)P_2$ at GEMs under these conditions.

(**B**) Typical trajectories of PIP5K and MARCKS diffusing on the PM cytoplasmic surface, showing transient colocalization with GEMs (talin clusters, white puncta) (snapshot from movie S4).

(**C**) Partial cholesterol depletion reduces the GEM cluster size (talin copy number per cluster). Distributions of the number of Halo7-talin molecules in a GEM cluster ($n$ = 15 cells, $n$ = 3,159 ~ 8,281 clusters), before and after cholesterol depletion (basic data for Fig. 2K).

(**D to H**) Optogenetic partial depletion of $PI(4,5)P_2$ reduces GEM size (talin copy number per cluster) and GEM # density on the PM.

(D) Schema for optogenetic partial depletion of $PI(4,5)P_2$ from the PM using the CIBN-CRY2 system, where phosphoinositide 5-phosphatase (5PT) is linked to CRY2 (*33*).

(E) Representative confocal micrographs of mScar-CRY2-5PT and SiR:SNAPf-2xPH ($PI(4,5)P_2$ probe), recorded before, 1 min after, and 15 min after the blue light irradiation (Before, After, and Recovery, respectively). Both mScar-CRY2 conjugates (WT-5PT and catalytically inactive D523G-5PT) were recruited to the PM after irradiation, whereas a decrease in the $PI(4,5)P_2$ probe signal was observed only in cells expressing (mScar-CRY2-)WT-5PT.

(F) Time courses of the $PI(4,5)P_2$ level in the PM (mean ± SEM, $n$ = 10 cells).

(G) Typical images of GEMs (Halo7-talin clusters) showing that GEM number density in the apical PM and GEM cluster size decrease immediately after irradiation in cells expressing (mScar-CRY2-)5PT, with D523G-5PT as the control.

(H) Mean talin copy number per GEM (cluster size) and mean GEM number density on the apical PM ($n$ = 10 cells). The GEM cluster size and number density decrease after light-induced $PI(4,5)P_2$ reduction and recover after illumination stops.

(**I and J**) Drugs that reduce $PI(4,5)P_2$ levels decrease GEM cluster size (talin copy number).

(I) Representative confocal micrographs of the $PI(4,5)P_2$ probe, SNAPf-2xPH, showing reduced $PI(4,5)P_2$ levels in the PM after drug treatments (*34*). The $Ca^{2+}$ ionophore ionomycin induces $PI(4,5)P_2$ hydrolysis by activating the $Ca^{2+}$-dependent PLC. Phenylarsine oxide and quercetin inhibit phosphatidylinositol 4-kinase, reducing PI(4)P production, the main substrate for $PI(4,5)P_2$ generation.

(J) Distributions of the number of Halo7-talin molecules in a GEM cluster ($n$ = 15 cells, $n$ = 3,159 ~ 8,281 clusters) before and after drug-induced partial depletion of $PI(4,5)P_2$ (basic data for Fig. 2K).

Games-Howell's test for (A and H), two-tailed Welch's t-test for (A), and Steel-Dwass' test for (C and J).

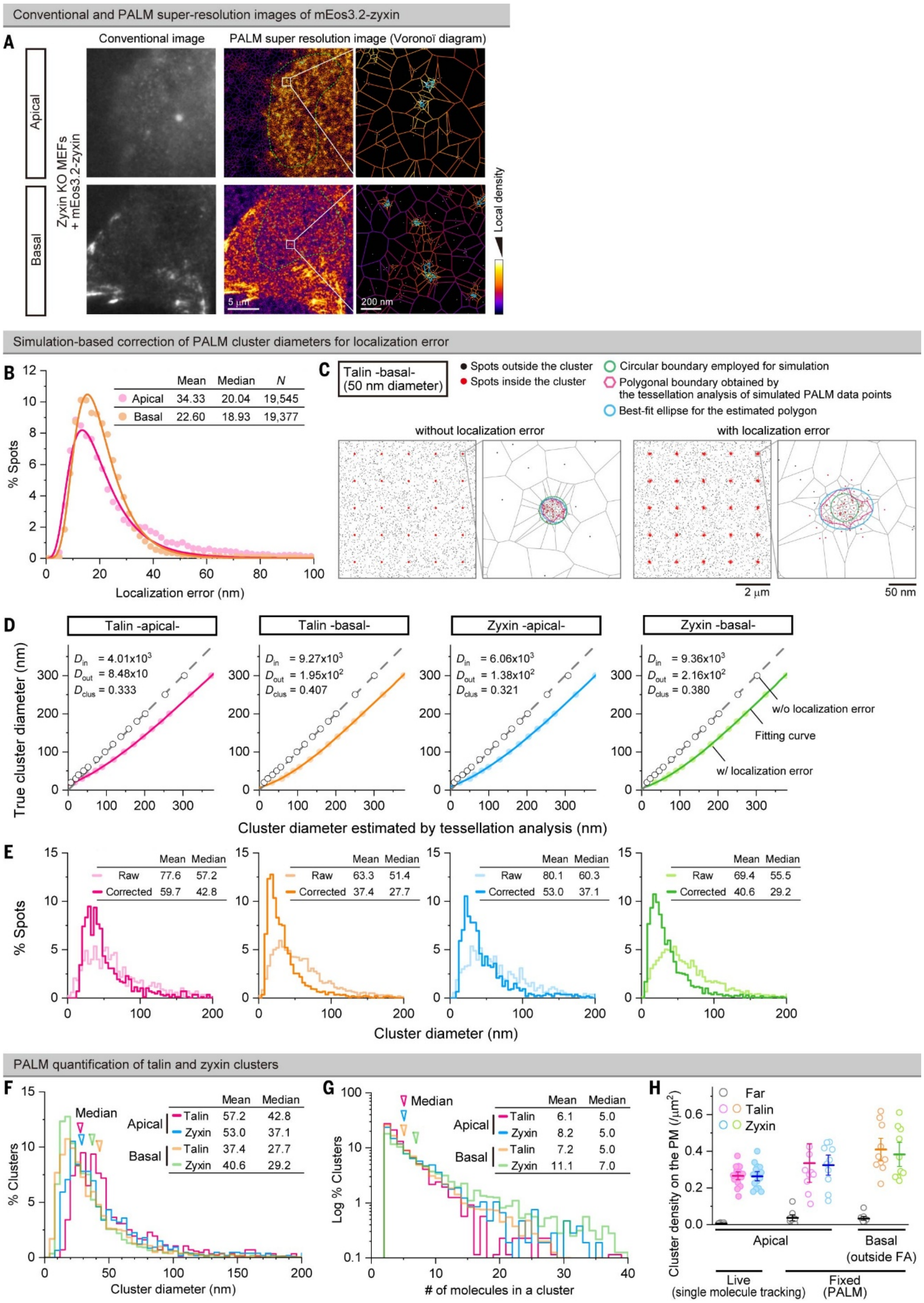


**Fig. S5. PALM-based detection and analysis of talin and zyxin clusters.**
(**A**) Conventional and PALM super-resolution images (Voronoï diagram) of GEM, visualized by mEos3.2-zyxin on the apical and basal PMs. The areas surrounded by the green curve (middle image) indicate regions with good focus (top; apical PM, which is tilted) or without obvious focal

adhesions (bottom; basal PM) and were used for further analysis. The cyan lines in the right images indicate the smallest polygon enclosing each clustered region.

(**B to E**) Evaluation of the true cluster diameter from the PALM data by correcting for localization errors using simulations (*115*).

(B) Distributions of the localization errors of mEos3.2 on the apical and basal PMs.

(C) Representative simulation workflow and typical results: Cluster centers were regularly placed at the average density observed by PALM experiments ($D_{\mathrm{clus}}$, clusters/μm$^2$). Circular boundaries with various diameters (10 – 300 nm) were drawn around the cluster centers, and localized-molecule coordinates (dots) were generated at experimentally matched densities inside and outside the boundary ($D_{\mathrm{in}}$ and $D_{\mathrm{out}}$ spots/μm$^2$, respectively, determined from experiments; C, two left panels). Localization errors were then applied to each coordinate by sampling from the distributions in (B) (C, two right panels).

(D) Calibration curves relating the cluster diameter estimated by the tessellation analysis (x axis) to the true cluster diameter (y axis). Simulated PALM datasets were processed identically to experimental data, and the resulting estimated diameters were plotted against the true input diameters. Each curve was fitted with a cubic polynomial and used for calibration.

(E) Distributions of raw (uncorrected) and calibrated (localization-error–corrected) cluster diameters.

(**F and G**) Distributions of the calibrated GEM diameters (F) and the number of molecules per GEM (G).

(**H**) Number densities of GEMs on the apical and basal PMs (mean ± SEM).

| | Uniprot ID | Total (a.a.) | IDR (a.a.) | % IDR | LLPS Ref# |
|---|---|---|---|---|---|
| **Receptors** | | | | | |
| PDGFR-β | P09619 | 1,106 | 88 | 8.0 | 167 |
| EGFR | P00533 | 1,210 | 41 | 3.4 | 105 |
| CD59 | P13987 | 128 | 0 | 0.0 | - |
| Thy1 | P04216 | 161 | 0 | 0.0 | - |
| P2Y1 | P47900 | 373 | 0 | 0.0 | - |
| **GEM** | | | | | |
| β3 integrin | P05106 | 788 | 0 | 0.0 | - |
| Talin1 | Q9Y490 | 2,540 | 0 | 0.0 | 55,77 |
| RIAM | Q7Z5R6 | 666 | 253 | 38.0 | - |
| VASP | P50552 | 380 | 233 | 61.3 | 53,54,168 |
| Zyxin | Q15942 | 572 | 322 | 56.3 | 169,170 |
| **FA related** | | | | | |
| Paxillin | P49023 | 591 | 269 | 45.5 | 55,87,170,171 |
| N-WASP | O00401 | 505 | 219 | 43.4 | 57,87 |
| FAK | Q05397 | 1,052 | 162 | 15.4 | 55,87,170 |
| Filamin-A | P21333 | 2,647 | 95 | 3.6 | - |
| Vinculin | P18206 | 1,134 | 31 | 2.7 | 172 |
| α-Actinin | P12814 | 892 | 0 | 0.0 | - |
| **PtdIns related** | | | | | |
| MARCKS | P29966 | 332 | 332 | 100.0 | 173 |
| PIP5KIγ | O60331 | 668 | 126 | 18.9 | - |
| PIK3CA | P42336 | 1,068 | 0 | 0.0 | 174-177 |
| **RTK downstream** | | | | | |
| Raf1 | P04049 | 648 | 115 | 17.7 | - |
| Lyn | P07948 | 512 | 62 | 12.1 | - |
| Akt | P31749 | 480 | 57 | 11.9 | 175 |
| PLCγ1 | P19174 | 1,290 | 43 | 3.3 | 57,63,174,178 |
| JAK1 | P23458 | 1,154 | 0 | 0.0 | - |
| K-Ras | P01116 | 189 | 0 | 0.0 | 174 |
| **Other FA related proteins** (not examined in this report) | | | | | |
| p130Cas | P56945 | 870 | 220 | 25.3 | 87,88 |
| Nck1 | P16333 | 377 | 0 | 0.0 | 57,87 |
| Kindlin2 | Q96AC1 | 680 | 25 | 3.7 | 55,87,179 |
| β1 integrin | P05556 | 798 | 33 | 4.1 | 55,87,172,179 |
| ILK | Q13418 | 452 | 0 | 0.0 | 179 |
| GIT1 | Q9Y2X7 | 761 | 126 | 17.0 | 170,171,180 |
| β-pix | Q14155 | 803 | 129 | 16.1 | 171 |
| LIMD1 | Q9UGP4 | 676 | 261 | 38.6 | 169 |
| Liprin-α1 | Q13136 | 1,202 | 321 | 26.7 | 180,181 |
| ERC1 | Q8IUD2 | 1,116 | 97 | 8.7 | 180,181 |
| LL5α | Q86UU1 | 1,377 | 487 | 35.4 | 180,181 |
| KANK1 | Q14678 | 1,352 | 1,063 | 78.6 | 182 |

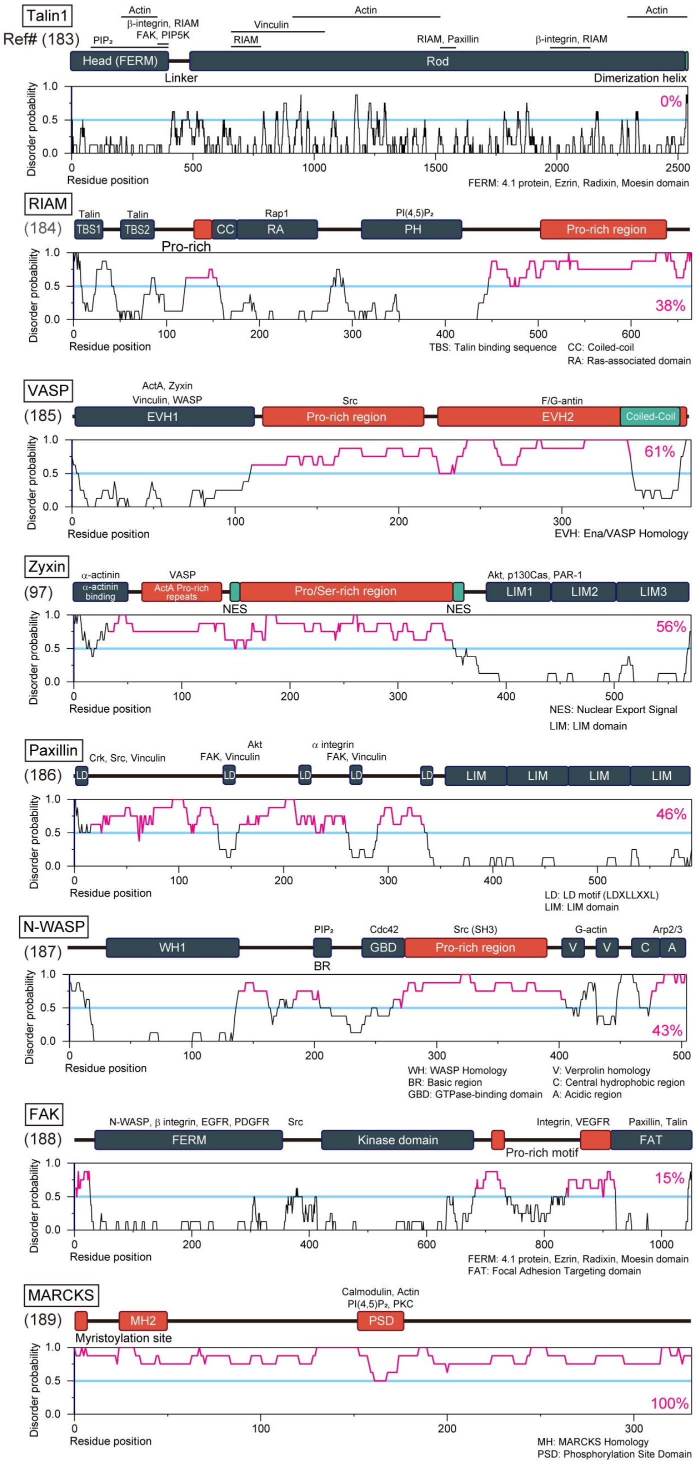


**Fig. S6. IDRs of the examined proteins.**

(**Left**) Summary of the IDRs of proteins (human) employed in this research, predicted by MobiDB 3.0 (*42*, *166*). The proteins with magenta shading contain at least one consecutive amino-acid stretch longer than 150 amino acids with the disorder probability greater than 0.5.

(**Right**) Domain structures of the proteins employed in this study. Domains shown in orange indicate those with higher disorder probabilities. The proteins that can bind to particular domains are indicated on the top of the domains.

To generate phosphodeficient (Phos–) and phosphomimetic (Phos+) mutants of zyxin and VASP, all serines, threonines, and tyrosines were mutated. For Phos-, they were mutated to alanines, alanines, and phenylalanines ,respectively, while, for Phos+, all of them were mutated to glutamic acid (see data S1 for detailed sequences). In zyxin, 28 out of these 34 sites are located within the IDR, whereas in VASP, 21 out of 27 such sites are located within IDR.

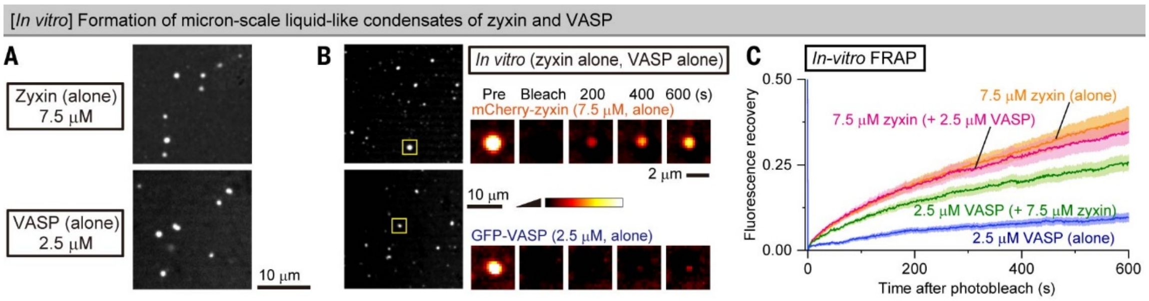


**Fig. S7. *In vitro* formation of micron-scale liquid-like condensates of zyxin alone and VASP alone.**

(**A**) Zyxin (mCherry-zyxin) alone and VASP (mGFP-VASP) alone form liquid-like micron-scale condensates *in vitro* upon the addition of PEG8K (final 1%).

(**B and C**) FRAP data were obtained 10 min after PEG8K addition (mean ± SEM, *n* = 10 condensates). Condensates of zyxin-VASP mixture exhibited intermediate FRAP recovery between VASP-only and zyxin-only condensates.

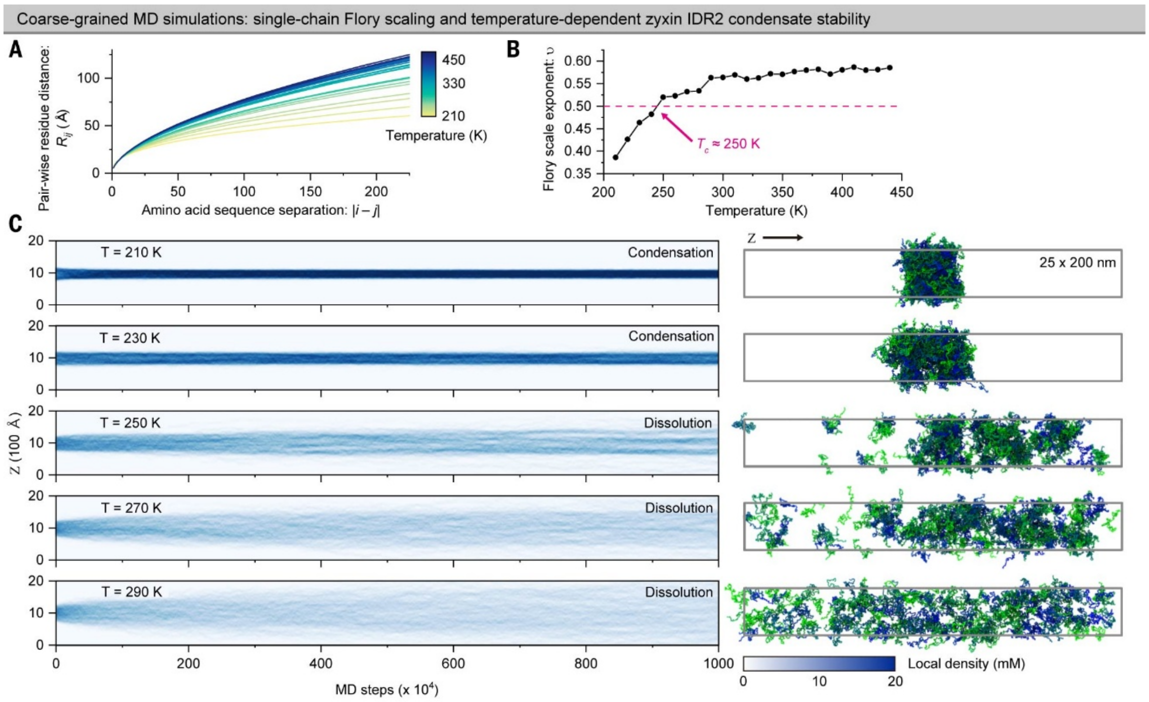


**Fig. S8. Coarse-grained MD simulations of zyxin IDR2.**

Coarse-grained (CG) molecular dynamics (MD) simulations of zyxin IDR2. Throughout this figure, 'temperature' denotes the CG-MD simulation (thermostat) temperature, a model control parameter, and is not directly comparable to experimental temperature. Also see Fig. 3, D to F.

(**A and B**) Single-chain conformational analysis of zyxin (IDR2) across temperatures. (A) Pairwise residue distance $R_{ij}$ plotted as a function of sequence separation $|i - j|$ for simulations performed from 210 K to 440 K (10-K intervals; CG simulation temperatures; see above). The results were fitted with a function, $R_{ij} \sim K|i - j|^{\nu}$, where $K$ and $\nu$ are fitting parameters ($\nu$ is the Flory scaling exponent). (B) Temperature dependence of $\nu$ extracted from the fits in (A). The red dashed line marks $\nu$ = 0.5, corresponding to the θ-condition. The simulation temperature at which $\nu$ approaches 0.5 is used as an estimate of the critical temperature ($T_c$) for phase separation.

(**C**) Temperature-dependent condensate density evolution of zyxin (IDR2) in CG MD simulations. (Left) Local density profiles along the z-axis for simulations at 210–290 K, shown as time-evolving traces. At lower simulation temperatures (210-230 K), densities remain localized, indicating a stable, compact condensate. With increasing simulation temperature, the density profile broadens, consistent with partial or complete dissolution of the condensate. (Right) Representative final snapshots at the corresponding temperatures. At low temperatures, chains form a compact droplet-like assembly; at higher temperatures, chains adopt a more dispersed state. Individual chains are colored from green to blue for visualization.

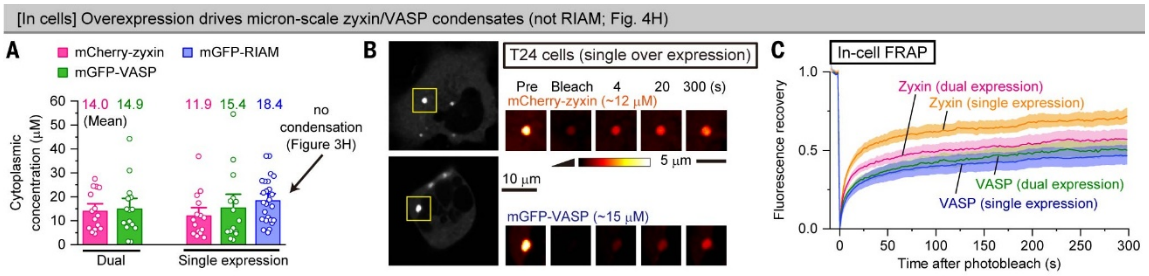


**Fig. S9. Overexpression-induced micron-scale co-condensates of zyxin and VASP in cells.**

In cells (T24 cells), overexpression of zyxin alone (mCherry-zyxin), VASP alone (mGFP-VASP), or both molecules induces micron-scale liquid-hydrogel-like condensates.

(**A**) Cytoplasmic concentrations of highly overexpressed mCherry-zyxin, mGFP-VASP, and mGFP-RIAM used for FRAP assays shown in (B) and (C), as well as in Figs. 3, B and C, and 9C (mean ± SEM, *n* = 15 cells). Concentrations were estimated by confocal microscopy, using purified mCherry-zyxin and mGFP-VASP in PBS as standards. Relative to the endogenous levels in fig. S10, zyxin and VASP are overexpressed by ≈19× and ≈24×, respectively.

(**B**) Typical FRAP image series for condensates of mCherry-zyxin alone (top) and mGFP-VASP alone (bottom).

(**C**) FRAP recovery curves (mean ± SEM, *n* = 15 condensates). Condensates of the zyxin-VASP mixture exhibited intermediate recovery between VASP-only and zyxin-only condensates, consistent with the observations made *in vitro*.

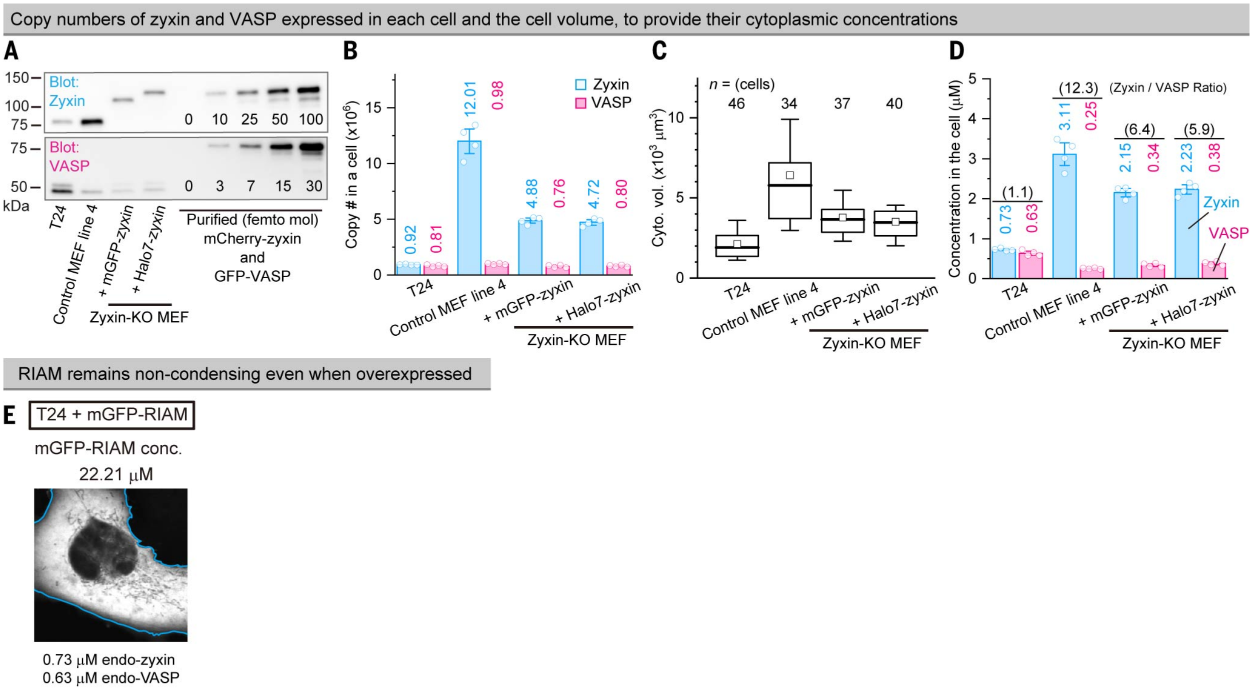


**Fig. S10. Quantification of endogenous and tagged zyxin/VASP concentrations and the result of RIAM overexpression.**

(**A**) Typical western blots used to estimate endogenous and tagged zyxin/VASP amounts. Absolute protein amounts were calibrated using purified mCherry-zyxin and mGFP-VASP standards.

(**B**) Total copy numbers per cell determined from the western blots (see A; mean ± SEM, $n = 4$ replicates). Endogenous VASP concentrations are consistent with a previous report (*190*).

(**C**) Cytoplasmic volume (cell volume minus nuclear volume) determined by confocal microscopy (mean ± SEM, $n = 4$ replicates).

(**D**) Cytoplasmic concentrations of endogenous and tagged zyxin and VASP (mean ± SEM).

(**E**) RIAM does not form detectable micron-scale condensates even at concentrations much higher than endogenous levels (confocal image).

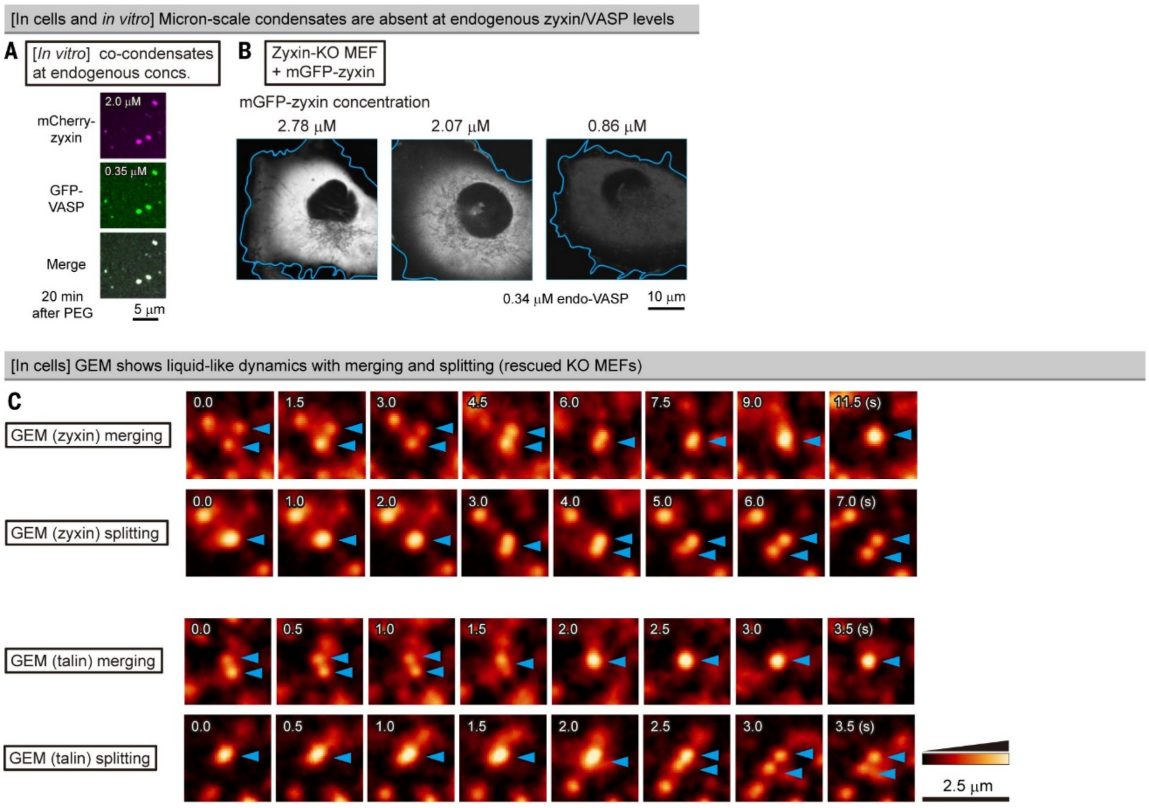


**Fig. S11. Physiological expression levels of zyxin do not form micron-scale condensates in cells and nanoscale GEMs undergo fusion and fission.**

(**A**) *In vitro*, zyxin and VASP form micron-scale co-condensates even at physiological endogenous concentrations in the presence of 1% PEG8K.

(**B**) mGFP-zyxin expressed at endogenous levels in zyxin-KO MEFs does not form detectable micron-scale condensates in cells.

(**C**) Typical images of nanoscale zyxin or talin clusters (GEMs) undergoing dynamic merging and splitting.

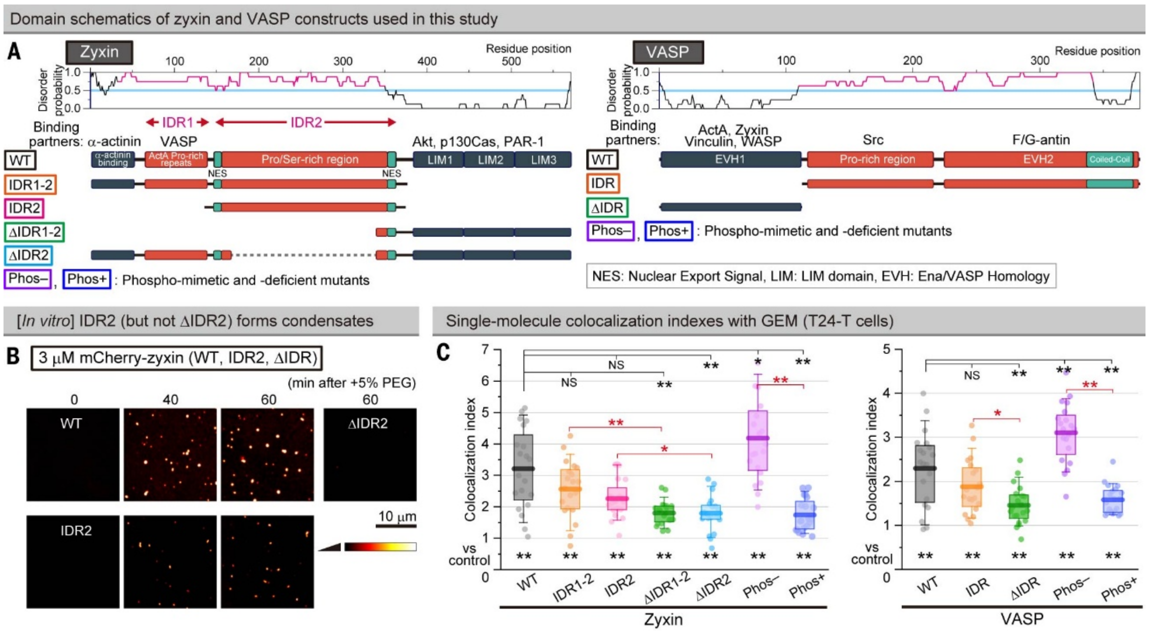


**Fig. S12. Zyxin and VASP's IDR constructs partition into GEMs.**

(**A**) Domain structures of zyxin, VASP, and their mutants employed in this study.

(**B**) Zyxin's IDR2, but not ΔIDR2, forms liquid-like micron-scale condensates *in vitro*.

(**C**) Single-molecule colocalization indexes of full-length and partial sequences of zyxin and VASP (± IDRs, Halo7-tagged) with GEM (talin-identified clusters) in T24-T cells.

Two-tailed Welch's t-test and Games-Howell's test for (C).

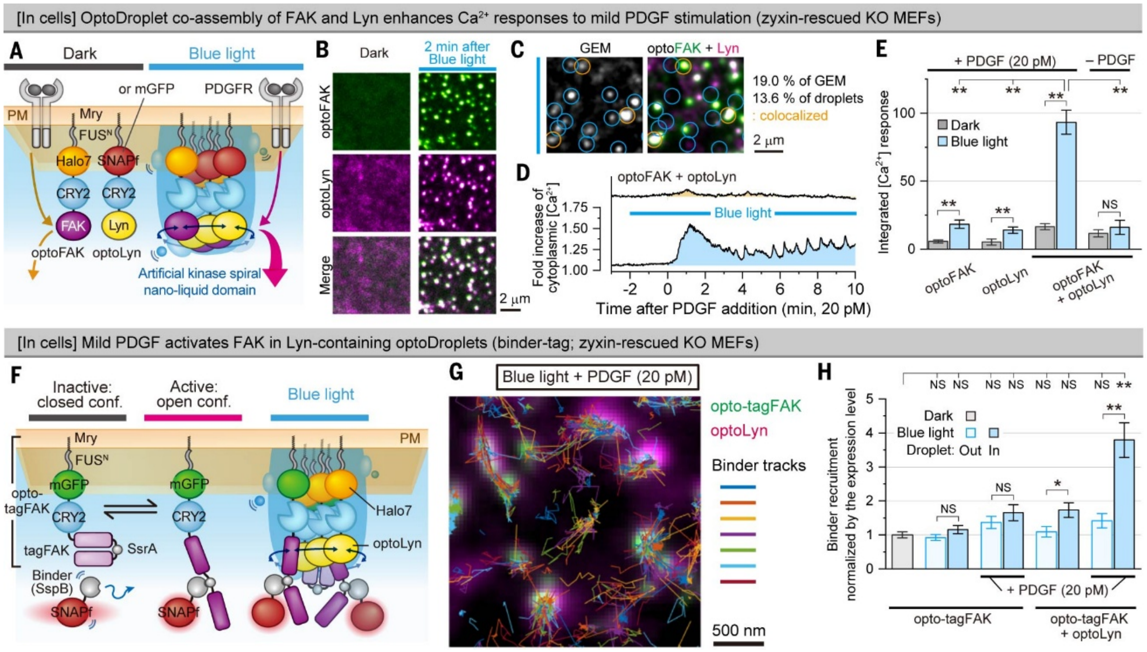


**Fig. S13. Figure S6. Mild PDGF stimulation leads to enhanced $Ca^{2+}$ mobilization under conditions where the mutual activation of FAK and SFK (Lyn) is optogenetically induced.**

(**A to E**) OptoDroplet-induced liquid-like co-assembly (*68*, *69*) of FAK and Lyn enhances $Ca^{2+}$ mobilization under mild PDGF stimulation, beyond simple additivity.

(A) Schematic of the optoDroplet experiments (*68*, *69*). OptoFAK and optoLyn (FAK and Lyn fused to a membrane-targeted optoDroplet cassette consisting of Src myristoylation signal peptide, FUS N-terminal IDR, a fluorescent tag protein, and the light-sensitive protein CRY2) were expressed in zyxin-KO MEFs rescued with mGFP-zyxin, and cytoplasmic $Ca^{2+}$ concentrations were monitored before and during blue light illumination with or without 20 pM PDGF (mild stimulation).

(B) Representative confocal micrographs of optoFAK (Halo7) and optoLyn (SNAPf) on the basal PM.

(**C**) Typical images of GEMs (talin clusters) and light-induced optoDroplets of optoFAK (Halo7) and optoLyn (SNAPf), showing partial colocalization (13.6 ± 1.3% of optoDroplets colocalized with GEMs, and 19.0 ± 2.4% of GEMs colocalized with optoDroplets).

(D) Typical $Ca^{2+}$ mobilization time course upon the addition of 20 pM PDGF.

(E) OptoDroplet formation (co-condensation) of optoFAK and optoLyn markedly enhances $Ca^{2+}$ responses upon mild PDGF stimulation. The graph shows the total $Ca^{2+}$ signal (AUC of traces as in D) (mean ± SEM, $n$ = 196 ~ 278 cells).

(**F to H**) FAK activation assay using a binder-tag readout. Enhancement of FAK activity via the formation of optoDroplets of optoFAK and optoLyn.

(F) Schematic of the binder-tag system to report FAK activation before and after optoDroplet induction with or without mild PDGF stimulation. In this binder-tag system, an SsrA peptide is inserted into the central region of FAK so that it can become accessible in the open/active conformation, enabling the binding of SNAPf-tagged SspB (*67*).

(G) Representative two-color image of opto-SsrA-FAK (green) and optoLyn (magenta) overlaid with trajectories of SspB after blue-light illumination and mild PDGF stimulation, indicating SspB recruitment to the optoDroplets.

(H) FAK activity increases ≈4-fold when both optoFAK and optoLyn are present in the same optoDroplet under mild PDGF stimulation, quantified by SspB recruitment normalized by the expression level (mean ± SEM, $n = 15$ cells).

Games-Howell's test for E and H.

The mild PDGF condition activates Lyn downstream of PDGFR but does not measurably activate FAK under these conditions (Fig. 5E). However, when light illumination induces the co-assembly of optoFAK and optoLyn into liquid-like optoDroplets, FAK becomes responsive to the mild PDGF input. Using a previously developed binder assay for the open, active conformation of FAK (*67*), we detected the robust recruitment of the SspB binder to the optoDroplets after mild PDGF stimulation (F to H). These observations suggest that, under the nanoscale co-confinement of Lyn and FAK in a liquid state, the Lyn initially activated by PDGFR can phosphorylate/activate FAK (which is not generally activated by the mild PDGF stimulation), which in turn can phosphorylate Lyn, inducing a chain (spiral-like) mutual activation reaction.

Meanwhile, without light activation, mild PDGF stimulation elicited only a slight $Ca^{2+}$ response, but upon the formation of a light-induced optoDroplet containing *both* optoFAK and optoLyn, the $Ca^{2+}$ response was enhanced by ≈6-fold relative to the dark state (D and E). Because $Ca^{2+}$ mobilization in this setting is downstream of PLCγ1–$IP_3$ signaling, these results are consistent with the idea that cooperative Lyn–FAK co-activation within the liquid-like optoDroplet promotes PLCγ1 recruitment/activation, thereby potentiating the $IP_3$–$Ca^{2+}$ output under otherwise subthreshold PDGF stimulation conditions.

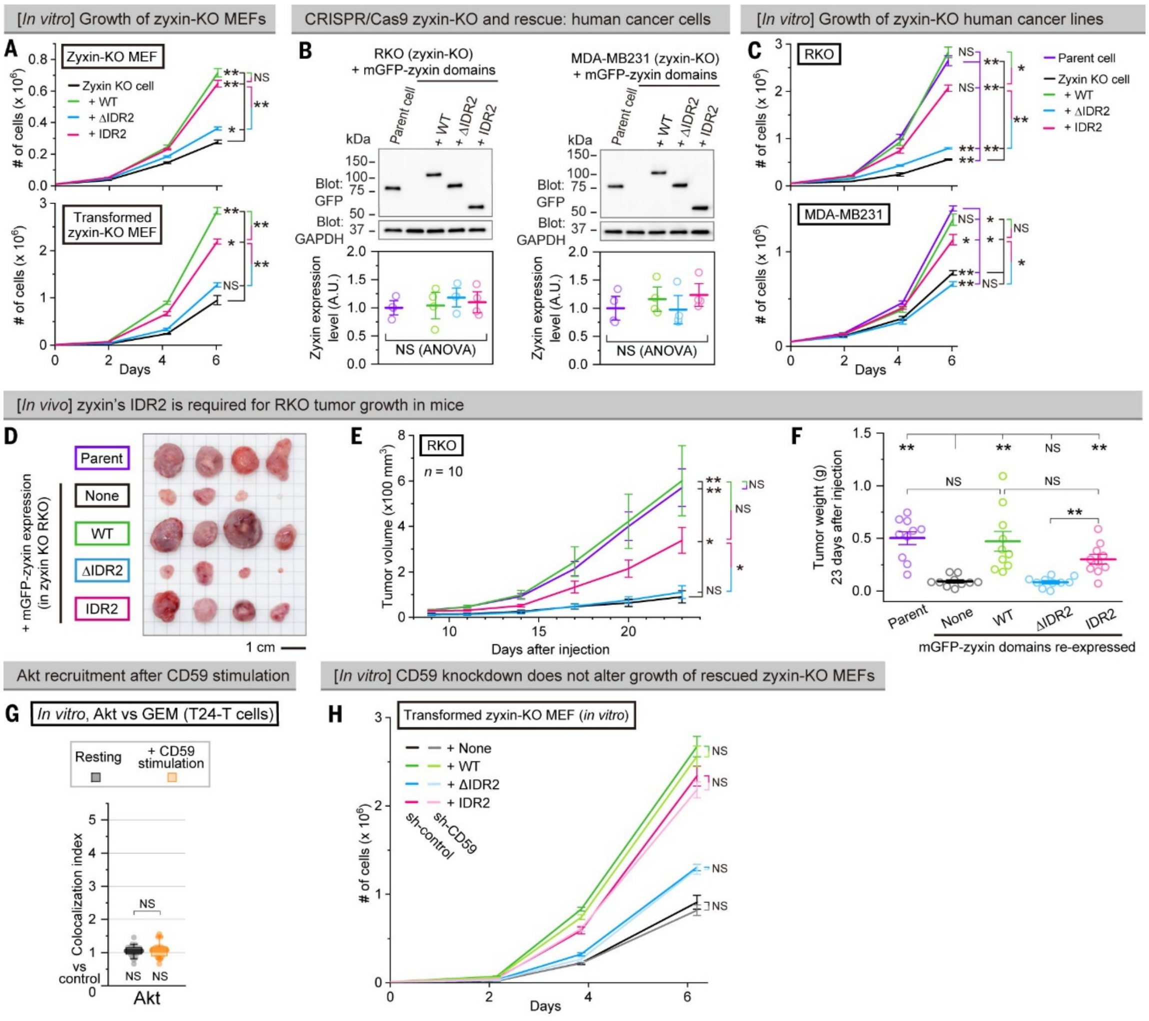


**Fig. S14. Cell proliferation comparison between *in vivo* and in culture with and without CD59 knockdown, examination using human cancer cell lines, and Akt recruitment to GEMs in cultured cells.**

(**A**) *In vitro* counterparts of Fig. 6, A to D. *In vitro* proliferation of zyxin-KO MEFs rescued with WT-zyxin or IDR2 is faster than that rescued with ΔIDR2 (mean ± SEM, $n$ = 3 replicates).

(**B**) Generation of zyxin-KO cell lines from human colon (RKO) and breast (MDA-MB231) cancer cell lines using CRISPR/Cas9 and re-expression of WT-zyxin, IDR2, or ΔIDR2 at levels comparable to endogenous zyxin in the respective parental lines. Typical western blot pattern (top) and band quantification results (bottom; normalized by the GAPDH band intensities; mean ± SEM, $n$ = 4 replicates) are shown.

(**C**) *In vitro* proliferation of zyxin-KO human cancer cell lines rescued with WT-zyxin or IDR2 is faster than when rescued with ΔIDR2 (mean ± SEM, $n$ = 3 replicates).

(**D to F**) Results obtained in mice showing that the presence of zyxin's IDR2 is critical for the growth of solid tumors of RKO cells in mice.

(D) Representative images of excised tumors.

(E) Tumor growth curves (mean ± SEM, $n$ = 10 mice).

(F) Weight of excised tumors (RKO cells) 29 days after injection (mean ± SEM, $n$ = 10 tumors).

(**G**) CD59 stimulation of cells in vitro (after serum starvation) does not induce Akt recruitment to GEMs ($n$ = 20 cells), at variance with the results in vivo (Fig. 6E).
(**H**) *In vitro* counterparts of Fig. 6, F to H. *In vitro* growth curves show that CD59 knockdown does not detectably affect proliferation under standard culture conditions, in contrast to the *in vivo* requirement for CD59 in tumor growth (mean ± SEM, $n$ = 3 replicates).
Games-Howell's test for (A, C, E, and F to H) and one-way ANOVA for (B).

**Movie S1. Visualizing the talin and zyxin clusters on the apical PM by rolling averaging.**

Movies recorded at video rate (**top**) and those after rolling averaging over 30 frames and background subtraction (**bottom**) are replayed at video rate. Halo7-talin expressed in a talin1/2-dKO MEF, Halo7-zyxin in a zyxin-KO MEF, and Halo7-farnesyl expressed in a talin2 KO MEF were labeled with the SaraFluor650-conjugated Halo-ligand and observed on the apical PM. After rolling averaging, the signals from the fast-diffusing molecules in the cytoplasm and on the PM, like Halo7-farnesyl, became averaged out. Meanwhile, immobile and slowly diffusing molecules and molecular clusters bound to the PM were visible even after rolling averaging (marked by circles in the bottom movies). The original data for Fig. 1A and fig. S1G. Representative movies among 20 movies.

**Movie S2. Transient recruitment of a CD59 cluster and a PDGF molecule to a GEM.**

**(Left)** Real-time movie of PDGFR molecules (magenta, with a trajectory; SaraFluor650 label) superimposed on the rolling-averaged movie of GEMs (observed as SNAPf-talin clusters in a Talin1/2-dKO MEF; green; TMR label) observed on the apical PM (recorded at video rate) after PDGF stimulation. PDGFR spots might represent PDGFR monomers or oligomers, but due to the presence of endogenous non-labeled PDGFR molecules, we could not differentiate these two possibilities. The original data for Fig. 1C, left. A representative movie among 20 movies.

**(Right)** Real-time movie of CD59 clusters formed by the addition of anti-CD59 mAb-coated fluorescent beads (magenta, with a trajectory), superimposed on the rolling-averaged movie of GEMs (observed as SNAPf-talin clusters in a Talin1/2-dKO MEF; green; TMR label) observed on the apical PM (recorded at video rate). The original data for Fig. 1C, right. A representative movie among 20 movies.

**Movie S3. Simultaneous triple-color single fluorescent particle/molecule tracking of GEMs, CD59 clusters, and PLCγ1 molecules.**

Real-time movie of engaged CD59 clusters (magenta, anti-CD59 mAb-coated fluorescent beads, with a trajectory) and single PLCγ1-Halo7 molecules (cyan; SaraFluor650 label) superimposed on the rolling-averaged movie of GEMs (observed as SNAPf-talin clusters; green; TMR label) on the apical PM of a T24-T cell, recorded at video rate. The original data for Fig. 1G. A representative movie among 30 movies.

**Movie S4. Transient recruitment of a FAK molecule and a PIP5K molecule to a GEM upon PDGF stimulation.**

Real-time movies of Halo7-PIP5K without stimulation (**left**) and Halo7-FAK after PDGF stimulation (**right**) (magenta, with a trajectory; SaraFluor650 label) superimposed on a rolling-averaged movie of GEMs (observed as SNAPf-talin clusters; green; TMR label) on the apical PM of a T24-T cell, recorded at video rate. The original data for Fig. 4B and fig. S4B bottom, respectively. A representative movie among 20 movies.

**Movie S5. Coarse-grained MD simulation of a zyxin IDR2 condensate.**
Movie of a coarse-grained MD simulation of a 200-chain zyxin IDR2 condensate at 210 K. The simulation time step was 0.1 ns, and the total simulated time was 1 ms ($10^7$ steps). The first $2 \times 10^6$ steps are shown. The original data for Fig. 3E.

**Data S1. (separate file)**
**Plasmids used in this study and their complete sequences.**
List of plasmids used in this study in an Excel table and plasmid sequence files in GenBank format with annotations.

**Data S2. (separate file)**
**Detailed experimental conditions and results including all statistical parameters.**
Detailed experimental conditions (including the employed tag proteins, fluorescent dyes, and cells) and results (including the raw data, the number of replicates, fitting results, and statistical test results).

### References and Notes

109. K. Naruse, X. Sai, N. Yokoyama, M. Sokabe, Uni-axial cyclic stretch induces c-src activation and translocation in human endothelial cells via SA channel activation. *FEBS Lett.* **441**, 111–115 (1998).

110. M. Ihara, H. Tomimoto, H. Kitayama, Y. Morioka, I. Akiguchi, H. Shibasaki, M. Noda, M. Kinoshita, Association of the cytoskeletal GTP-binding protein Sept4/H5 with cytoplasmic inclusions found in Parkinson's disease and other synucleinopathies. *J. Biol. Chem.* **278**, 24095–24102 (2003).

111. M. Hooper, K. Hardy, A. Handyside, S. Hunter, M. Monk, HPRT-deficient (Lesch–Nyhan) mouse embryos derived from germline colonization by cultured cells. *Nature* **326**, 292–295 (1987).

112. S. Xia, Y. B. Lim, Z. Zhang, Y. Wang, S. Zhang, C. T. Lim, E. K. F. Yim, P. Kanchanawong, Nanoscale architecture of the cortical actin cytoskeleton in embryonic stem cells. *Cell Rep.* **28**, 1251-1267.e7 (2019).

113. S. A. McKinney, C. S. Murphy, K. L. Hazelwood, M. W. Davidson, L. L. Looger, A bright and photostable photoconvertible fluorescent protein. *Nat. Methods* **6**, 131–133 (2009).

114. T. K. Fujiwara, S. Takeuchi, Z. Kalay, Y. Nagai, T. A. Tsunoyama, T. Kalkbrenner, K. Iwasawa, K. P. Ritchie, K. G. N. Suzuki, A. Kusumi, Development of ultrafast camera-based single fluorescent-molecule imaging for cell biology. *J. Cell Biol.* **222**, e202110160 (2023).

115. T. K. Fujiwara, T. A. Tsunoyama, S. Takeuchi, Z. Kalay, Y. Nagai, T. Kalkbrenner, Y. L. Nemoto, L. H. Chen, A. C. E. Shibata, K. Iwasawa, K. P. Ritchie, K. G. N. Suzuki, A. Kusumi, Ultrafast single-molecule imaging reveals focal adhesion nano-architecture and molecular dynamics. *J. Cell Biol.* **222**, e202110162 (2023).

116. D. S. Bindels, L. Haarbosch, L. van Weeren, M. Postma, K. E. Wiese, M. Mastop, S. Aumonier, G. Gotthard, A. Royant, M. A. Hink, T. W. J. Gadella, mScarlet: a bright monomeric red fluorescent protein for cellular imaging. *Nat. Methods* **14**, 53–56 (2017).

117. M. Maekawa, G. D. Fairn, Complementary probes reveal that phosphatidylserine is required for the proper transbilayer distribution of cholesterol. *J. Cell Sci.* **128**, 1422–1433 (2015).

118. B. B. Johnson, P. C. Moe, D. Wang, K. Rossi, B. L. Trigatti, A. P. Heuck, Modifications in perfringolysin O domain 4 alter the cholesterol concentration threshold required for binding. *Biochemistry* **51**, 3373–3382 (2012).

119. T. A. Tsunoyama, Y. Watanabe, J. Goto, K. Naito, R. S. Kasai, K. G. N. Suzuki, T. K. Fujiwara, A. Kusumi, Super-long single-molecule tracking reveals dynamic-anchorage-induced integrin function. *Nat. Chem. Biol.* **14**, 497–506 (2018).

120. Y. L. Nemoto, R. J. Morris, H. Hijikata, T. A. Tsunoyama, A. C. E. Shibata, R. S. Kasai, A. Kusumi, T. K. Fujiwara, Dynamic Meso-Scale Anchorage of GPI-Anchored Receptors in the Plasma Membrane: Prion Protein vs. Thy1. *Cell Biochem Biophys* **75**, 399–412 (2017).

121. D. Ilić, E. A. C. Almeida, D. D. Schlaepfer, P. Dazin, S. Aizawa, C. H. Damsky, Extracellular matrix survival signals transduced by focal adhesion kinase suppress p53-mediated apoptosis. *J. Cell Biol.* **143**, 547–560 (1998).

122. H.-S. Lee, P. Anekal, C. J. Lim, C.-C. Liu, M. H. Ginsberg, Two modes of integrin activation form a binary molecular switch in adhesion maturation. *Mol. Biol. Cell* **24**, 1354–1362 (2013).

123. D. Tsuruta, M. Gonzales, S. B. Hopkinson, C. Otey, S. Khuon, R. D. Goldman, J. C. R. Jones, Microfilament-dependent movement of the β3 integrin subunit within focal contacts of endothelial cells. *FASEB J.* **16**, 866–868 (2002).

124. J. Okrut, S. Prakash, Q. Wu, M. J. S. Kelly, J. Taunton, Allosteric N-WASP activation by an inter-SH3 domain linker in Nck. *Proc. Natl Acad. Sci. U.S.A.* **112**, E6436–E6445 (2015).

125. Y. Sun, N. Thapa, A. C. Hedman, R. A. Anderson, Phosphatidylinositol 4,5-bisphosphate: targeted production and signaling. *Bioessays* **35**, 513–522 (2013).

126. L. E. Diamond, K. R. McCurry, E. R. Oldham, M. Tone, H. Waldmann, J. L. Platt, J. S. Logan, Human CD59 expressed in transgenic mouse hearts inhibits the activation of complement. *Transpl. Immunol.* **3**, 305–312 (1995).

127. C. M. Johannessen, J. S. Boehm, S. Y. Kim, S. R. Thomas, L. Wardwell, L. A. Johnson, C. M. Emery, N. Stransky, A. P. Cogdill, J. Barretina, G. Caponigro, H. Hieronymus, R. R. Murray, K. Salehi-Ashtiani, D. E. Hill, M. Vidal, J. J. Zhao, X. Yang, O. Alkan, S. Kim, J. L. Harris, C. J. Wilson, V. E. Myer, P. M. Finan, D. E. Root, T. M. Roberts, T. Golub, K. T. Flaherty, R. Dummer, B. L. Weber, W. R. Sellers, R. Schlegel, J. A. Wargo, W. C. Hahn, L. A. Garraway, COT drives resistance to RAF inhibition through MAP kinase pathway reactivation. *Nature* **468**, 968–972 (2010).

128. N. P. Jones, J. Peak, S. Brader, S. A. Eccles, M. Katan, PLCγ1 is essential for early events in integrin signalling required for cell motility. *J. Cell Sci.* **118**, 2695–2706 (2005).

129. S. Ishida, T. Matsu-ura, K. Fukami, T. Michikawa, K. Mikoshiba, Phospholipase C-β1 and β4 contribute to non-genetic cell-to-cell variability in histamine-induced calcium signals in HeLa cells. *PLoS One* **9**, e86410 (2014).

130. F. P. Lindberg, H. D. Gresham, E. Schwarz, E. J. Brown, Molecular cloning of integrin-associated protein: an immunoglobulin family member with multiple membrane-spanning domains implicated in alpha v beta 3-dependent ligand binding. *J. Cell Biol.* **123**, 485–496 (1993).

131. K. G. Suzuki, R. S. Kasai, K. M. Hirosawa, Y. L. Nemoto, M. Ishibashi, Y. Miwa, T. K. Fujiwara, A. Kusumi, Transient GPI-anchored protein homodimers are units for raft organization and function. *Nat. Chem. Biol.* **8**, 774–83 (2012).

132. A. Nakamura, C. Oki, K. Kato, S. Fujinuma, G. Maryu, K. Kuwata, T. Yoshii, M. Matsuda, K. Aoki, S. Tsukiji, Engineering orthogonal, plasma membrane-specific SLIPT systems for multiplexed chemical control of signaling pathways in living single cells. *ACS Chem. Biol.* **15**, 1004–1015 (2020).

133. H. Sun, R. Taneja, Analysis of transformation and tumorigenicity using mouse embryonic fibroblast cells. *Methods Mol. Biol.* **383**, 303–310 (2007).

134. C. L. Harris, S. M. Hanna, M. Mizuno, D. S. Holt, K. J. Marchbank, B. P. Morgan, Characterization of the mouse analogues of CD59 using novel monoclonal antibodies: tissue distribution and functional comparison. *Immunology* **109**, 117–126 (2003).

135. I. Štefanová, I. Hilgert, H. Krištofová, R. Brown, M. G. Low, V. Hořejši, Characterization of a broadly expressed human leucocyte surface antigen MEM-43 anchored in membrane through phosphatidylinositol. *Mol. Immunol.* **26**, 153–161 (1989).

136. N. Madore, K. L. Smith, C. H. Graham, A. Jen, K. Brady, S. Hall, R. Morris, Functionally different GPI proteins are organized in different domains on the neuronal surface. *EMBO J.* **18**, 6917–6926 (1999).

137. D. W. Mason, A. F. Williams, The kinetics of antibody binding to membrane antigens in solution and at the cell surface. *Biochem. J.* **187**, 1–20 (1980).

138. J. H. Hanke, J. P. Gardner, R. L. Dow, P. S. Changelian, W. H. Brissette, E. J. Weringer, B. A. Pollok, P. A. Connelly, Discovery of a novel, potent, and Src family-selective tyrosine kinase inhibitor. *J. Biol. Chem.* **271**, 695–701 (1996).

139. J. K. Slack-Davis, K. H. Martin, R. W. Tilghman, M. Iwanicki, E. J. Ung, C. Autry, M. J. Luzzio, B. Cooper, J. C. Kath, W. G. Roberts, J. T. Parsons, Cellular characterization of a novel focal adhesion kinase inhibitor. *J. Biol. Chem.* **282**, 14845–14852 (2007).

140. A. Kosenko, N. Hoshi, A change in configuration of the calmodulin-KCNQ channel complex underlies $Ca^{2+}$-dependent modulation of KCNQ channel activity. *PLoS One* **8**, e82290 (2013).

141. K. A. Johnson, M. R. Budicini, N. Bhattarai, T. Sharma, S. Urata, B. S. Gerstman, P. P. Chapagain, S. Li, R. V. Stahelin, PI(4,5)P2 binding sites in the Ebola virus matrix protein VP40 modulate assembly and budding. *J. Lipid Res.* **65**, 100512 (2024).

142. S. V. Ulianov, A. K. Velichko, M. D. Magnitov, A. V. Luzhin, A. K. Golov, N. Ovsyannikova, I. I. Kireev, A. S. Gavrikov, A. S. Mishin, A. K. Garaev, A. V. Tyakht, A. A. Gavrilov, O. L. Kantidze, S. V. Razin, Suppression of liquid–liquid phase separation by 1,6-hexanediol partially compromises the 3D genome organization in living cells. *Nucleic Acids Res.* **49**, 10524–10541 (2021).

143. P. Zhou, R. S. Kasai, W. Fujita, T. A. Tsunoyama, H. Neyama, H. Ueda, T. Yokoyama, M. Sakamoto, S. Pigolotti, T. K. Fujiwara, A. Kusumi, Single-molecule characterization of opioid receptor heterodimers reveals soluble μ-δ dimer blocker peptide alleviates morphine tolerance. *Nat. Commun.* **16**, 9859 (2025).

144. T. Fujiwara, K. Ritchie, H. Murakoshi, K. Jacobson, A. Kusumi, Phospholipids undergo hop diffusion in compartmentalized cell membrane. *J. Cell Biol.* **157**, 1071–1082 (2002).

145. I. Koyama-Honda, K. Ritchie, T. Fujiwara, R. Iino, H. Murakoshi, R. S. Kasai, A. Kusumi, Fluorescence imaging for monitoring the colocalization of two single molecules in living cells. *Biophys. J.* **88**, 2126–2136 (2005).

146. A. C. Shibata, T. K. Fujiwara, L. Chen, K. G. Suzuki, Y. Ishikawa, Y. L. Nemoto, Y. Miwa, Z. Kalay, R. Chadda, K. Naruse, A. Kusumi, Archipelago architecture of the focal adhesion: membrane molecules freely enter and exit from the focal adhesion zone. *Cytoskeleton* **69**, 380–92 (2012).

147. S. J. Sahl, M. Leutenegger, M. Hilbert, S. W. Hell, C. Eggeling, Fast molecular tracking maps nanoscale dynamics of plasma membrane lipids. *Proc. Natl Acad. Sci. U.S.A.* **107**, 6829–6834 (2010).

148. P. Zhou, T. A. Tsunoyama, R. S. Kasai, K. M. Hirosawa, Z. Kalay, A. Aladag, T. Fujiwara, S. Pigolotti, A. Kusumi, Single-molecule detection of transient dimerization of opioid receptors 1: Homodimers' effect on signaling and internalization. *bioRxiv*, 2024.07.25.605080 (2024).

149. S. L. Veatch, B. B. Machta, S. A. Shelby, E. N. Chiang, D. A. Holowka, B. A. Baird, Correlation functions quantify super-resolution images and estimate apparent clustering due to over-counting. *PLoS One* **7**, e31457 (2012).

150. K. Xu, G. Zhong, X. Zhuang, Actin, spectrin, and associated proteins form a periodic cytoskeletal structure in axons. *Science* **339**, 452–456 (2013).

151. K. A. K. Tanaka, K. G. N. Suzuki, Y. M. Shirai, S. T. Shibutani, M. S. H. Miyahara, H. Tsuboi, M. Yahara, A. Yoshimura, S. Mayor, T. K. Fujiwara, A. Kusumi, Membrane molecules mobile even after chemical fixation. *Nat Methods* **7**, 865–866 (2010).

152. S.-H. Lee, J. Y. Shin, A. Lee, C. Bustamante, Counting single photoactivatable fluorescent molecules by photoactivated localization microscopy (PALM). *Proc. Natl Acad. Sci. U.S.A.* **109**, 17436–17441 (2012).

153. E. D. Zitter, D. Thédié, V. Mönkemöller, S. Hugelier, J. Beaudouin, V. Adam, M. Byrdin, L. V. Meervelt, P. Dedecker, D. Bourgeois, Mechanistic investigation of mEos4b reveals a strategy to reduce track interruptions in sptPALM. *Nat. Methods* **16**, 707–710 (2019).

154. M. Ovesný, P. Křížek, J. Borkovec, Z. Švindrych, G. M. Hagen, ThunderSTORM: a comprehensive ImageJ plug-in for PALM and STORM data analysis and super-resolution imaging. *Bioinformatics* **30**, 2389–2390 (2014).

155. S. Berg, D. Kutra, T. Kroeger, C. N. Straehle, B. X. Kausler, C. Haubold, M. Schiegg, J. Ales, T. Beier, M. Rudy, K. Eren, J. I. Cervantes, B. Xu, F. Beuttenmueller, A. Wolny, C. Zhang, U. Koethe, F. A. Hamprecht, A. Kreshuk, ilastik: interactive machine learning for (bio)image analysis. *Nat. Methods* **16**, 1226–1232 (2019).

156. M. Dundr, T. Misteli, Measuring dynamics of nuclear proteins by photobleaching. *Curr. Protoc. Cell Biol.* **18**, 13.5.1-13.5.18 (2003).

157. N. Otsu, A threshold selection method from gray-level histograms. *IEEE Trans. Syst. Man Cybern.* **9**, 62–66 (1979).

158. N. J. Anthis, G. M. Clore, Sequence-specific determination of protein and peptide concentrations by absorbance at 205 nm: sequence-specific protein concentration at 205 nm. *Protein Sci.* **22**, 851–858 (2013).

159. M. M. Tomayko, C. P. Reynolds, Determination of subcutaneous tumor size in athymic (nude) mice. *Cancer Chemother. Pharmacol.* **24**, 148–154 (1989).

160. C. Tan, J. Jung, C. Kobayashi, D. U. L. Torre, S. Takada, Y. Sugita, Implementation of residue-level coarse-grained models in GENESIS for large-scale molecular dynamics simulations. *PLoS Comput. Biol.* **18**, e1009578 (2022).

161. J. Jung, K. Yagi, C. Tan, H. Oshima, T. Mori, I. Yu, Y. Matsunaga, C. Kobayashi, S. Ito, D. U. L. Torre, Y. Sugita, GENESIS 2.1: High-performance molecular dynamics software for enhanced sampling and free-energy calculations for atomistic, coarse-grained, and quantum mechanics/molecular mechanics models. *J. Phys. Chem. B* **128**, 6028–6048 (2024).

162. J. Jung, C. Tan, Y. Sugita, GENESIS CGDYN: large-scale coarse-grained MD simulation with dynamic load balancing for heterogeneous biomolecular systems. *Nat. Commun.* **15**, 3370 (2024).

163. J. Schindelin, I. Arganda-Carreras, E. Frise, V. Kaynig, M. Longair, T. Pietzsch, S. Preibisch, C. Rueden, S. Saalfeld, B. Schmid, J.-Y. Tinevez, D. J. White, V. Hartenstein, K. Eliceiri, P. Tomancak, A. Cardona, Fiji: an open-source platform for biological-image analysis. *Nat. Methods* **9**, 676–682 (2012).

164. U. Praekelt, P. M. Kopp, K. Rehm, S. Linder, N. Bate, B. Patel, E. Debrand, A. M. Manso, R. S. Ross, F. Conti, M.-Z. Zhang, R. C. Harris, R. Zent, D. R. Critchley, S. J. Monkley, New isoform-specific monoclonal antibodies reveal different sub-cellular localisations for talin1 and talin2. *Eur. J. Cell Biol.* **91**, 180–191 (2012).

165. G. D. Paolo, L. Pellegrini, K. Letinic, G. Cestra, R. Zoncu, S. Voronov, S. Chang, J. Guo, M. R. Wenk, P. D. Camilli, Recruitment and regulation of phosphatidylinositol phosphate kinase type 1γ by the FERM domain of talin. *Nature* **420**, 85–89 (2002).

166. I. de Curtis, Biomolecular condensates at the front: cell migration meets phase separation. *Trends Cell Biol.* **31**, 145–148 (2021).

167. C.-C. Lin, K. M. Suen, J. Lidster, J. E. Ladbury, The emerging role of receptor tyrosine kinase phase separation in cancer. *Trends Cell Biol.* **34**, 371–379 (2024).

168. K. Graham, A. Chandrasekaran, L. Wang, N. Yang, E. M. Lafer, P. Rangamani, J. C. Stachowiak, Liquid-like condensates mediate competition between actin branching and bundling. *Proc. Natl. Acad. Sci. U.S.A.* **121**, e2309152121 (2024).

169. Y. Wang, C. Zhang, W. Yang, S. Shao, X. Xu, Y. Sun, P. Li, L. Liang, C. Wu, LIMD1 phase separation contributes to cellular mechanics and durotaxis by regulating focal adhesion dynamics in response to force. *Dev. Cell* **56**, 1313-1325.e7 (2021).

170. P. Liang, Y. Wu, S. Zheng, J. Zhang, S. Yang, J. Wang, S. Ma, M. Zhang, Z. Gu, Q. Liu, W. Jiang, Q. Xing, B. Wang, Paxillin phase separation promotes focal adhesion assembly and integrin signaling. *J. Cell Biol.* **223**, e202209027 (2024).

171. J. Zhu, Q. Zhou, Y. Xia, L. Lin, J. Li, M. Peng, R. Zhang, M. Zhang, GIT/PIX condensates are modular and ideal for distinct compartmentalized cell signaling. *Mol. Cell* **79**, 782-796.e6 (2020).

172. T. Litschel, C. F. Kelley, X. Cheng, L. Babl, N. Mizuno, L. B. Case, P. Schwille, Membrane-induced 2D phase separation of the focal adhesion protein talin. *Nat. Commun.* **15**, 4986 (2024).

173. C. M. Lim, A. G. Díaz, M. Fuxreiter, F. W. Pun, A. Zhavoronkov, M. Vendruscolo, Multiomic prediction of therapeutic targets for human diseases associated with protein phase separation. *Proc. Natl. Acad. Sci. U.S.A.* **120**, e2300215120 (2023).

174. A. Tulpule, J. Guan, D. S. Neel, H. R. Allegakoen, Y. P. Lin, D. Brown, Y.-T. Chou, A. Heslin, N. Chatterjee, S. Perati, S. Menon, T. A. Nguyen, J. Debnath, A. D. Ramirez, X. Shi, B. Yang, S. Feng, S. Makhija, B. Huang, T. G. Bivona, Kinase-mediated RAS signaling via membraneless cytoplasmic protein granules. *Cell* **184**, 2649-2664. e18 (2021).

175. K. Zhou, Q. Chen, J. Chen, D. Liang, W. Feng, M. Liu, Q. Wang, R. Wang, Q. Ouyang, C. Quan, S. Chen, Spatiotemporal regulation of insulin signaling by liquid–liquid phase separation. *Cell Discov.* **8**, 64 (2022).

176. X. K. Gao, X. S. Rao, X. X. Cong, Z. K. Sheng, Y. T. Sun, S. B. Xu, J. F. Wang, Y. H. Liang, L. R. Lu, H. Ouyang, H. Ge, J. Guo, H. Wu, Q. M. Sun, H. Wu, Z. Bao, L. L. Zheng, Y. T. Zhou, Phase separation of insulin receptor substrate 1 drives the formation of insulin/IGF-1 signalosomes. *Cell Discov.* **8**, 60 (2022).

177. J. Sampson, M. W. Richards, J. Choi, A. M. Fry, R. Bayliss, Phase-separated foci of EML4-ALK facilitate signalling and depend upon an active kinase conformation. *EMBO Rep.* **22**, EMBR202153693 (2021).

178. L. Zeng, I. Palaia, A. Šarić, X. Su, PLCγ1 promotes phase separation of T cell signaling components. *J. Cell Biol.* **220**, e202009154 (2021).

179. N. Ma, F. Wu, J. Liu, Z. Wu, L. Wang, B. Li, Y. Liu, X. Dong, J. Hu, X. Fang, H. Zhang, D. Ai, J. Zhou, X. Wang, Kindlin-2 phase separation in response to flow controls vascular stability. *Circ. Res.* **135**, 1141–1160 (2024).

180. K. Sala, A. Corbetta, C. Minici, D. Tonoli, D. H. Murray, E. Cammarota, L. Ribolla, M. Ramella, R. Fesce, D. Mazza, M. Degano, I. de Curtis, The ERC1 scaffold protein implicated in cell motility drives the assembly of a liquid phase. *Sci. Rep.* **9**, 13530 (2019).

181. K. Sala, A. Raimondi, D. Tonoli, C. Tacchetti, I. de Curtis, Identification of a membrane-less compartment regulating invadosome function and motility. *Sci. Rep.* **8**, 1164 (2018).

182. K. Guo, J. Zhang, P. Huang, Y. Xu, W. Pan, K. Li, L. Chen, L. Luo, W. Yu, S. Chen, S. He, Z. Wei, C. Yu, KANK1 shapes focal adhesions by orchestrating protein binding, mechanical force sensing, and phase separation. *Cell Rep.* **42**, 113321 (2023).

183. D. A. Calderwood, I. D. Campbell, D. R. Critchley, Talins and kindlins: partners in integrin-mediated adhesion. *Nat. Rev. Mol. Cell Biol.* **14**, 503–517 (2013).

184. J. Yang, L. Zhu, H. Zhang, J. Hirbawi, K. Fukuda, P. Dwivedi, J. Liu, T. Byzova, E. F. Plow, J. Wu, J. Qin, Conformational activation of talin by RIAM triggers integrin-mediated cell adhesion. *Nat. Commun.* **5**, 5880 (2014).

185. M. Krause, E. W. Dent, J. E. Bear, J. J. Loureiro, F. B. Gertler, ENA/VASP proteins: regulators of the actin cytoskeleton and cell migration. *Cell Dev. Biol.* **19**, 541–564 (2003).

186. A. M. López-Colomé, I. Lee-Rivera, R. Benavides-Hidalgo, E. López, Paxillin: a crossroad in pathological cell migration. *J. Hematol. Oncol.* **10**, 50 (2017).

187. T. Takenawa, S. Suetsugu, The WASP–WAVE protein network: connecting the membrane to the cytoskeleton. *Nat. Rev. Mol. Cell Biol.* **8**, 37–48 (2007).

188. F. J. Sulzmaier, C. Jean, D. D. Schlaepfer, FAK in cancer: mechanistic findings and clinical applications. *Nat. Rev. Cancer* **14**, 598–610 (2014).

189. M. E. Amri, U. Fitzgerald, G. Schlosser, MARCKS and MARCKS-like proteins in development and regeneration. *J. Biomed. Sci.* **25**, 43 (2018).

190. J. D.- Guercio, L. Kurzawa, J. Mueller, G. Dimchev, M. Schaks, M. Nemethova, T. Pokrant, S. Brühmann, J. Linkner, L. Blanchoin, M. Sixt, K. Rottner, J. Faix, Loss of Ena/VASP interferes with lamellipodium architecture, motility and integrin-dependent adhesion. *eLife* **9**, e55351 (2020).
